# Discovery of a small peptide that increases yeast lifespan by enhancing the function of APC^Cdh1^ in nondividing quiescent cells

**DOI:** 10.64898/2026.02.17.706436

**Authors:** R.E. Harris, S.D. Postnikoff, N.K. Shukla, A.H. Harkness, H.M. Brooks, A. Khawaja, M.E. Malo, C.Z. Verdugo, B.M. Waddell, M. Wu, T.A.A. Harkness

## Abstract

Aging is accompanied by molecular hallmarks conserved from yeast to humans. One such hallmark, senescence, is an irreversible nondividing state that promotes aging. We previously reported that mutants of the yeast Anaphase Promoting Complex (APC) shorten the lifespan of both dividing and nondividing cells. Here, we propose that activation of the APC will promote the maintenance of quiescent yeast cells, thereby delaying senescence and aging. We observed that APC activity in aging quiescent cells becomes increasingly impaired, as the APC substrates Clb1 and Mps1 accumulated within aging cells reintroduced back into the cell cycle. To identify peptide activators of the APC, we used a yeast 2-hybrid screen to recover peptides that interacted with the APC subunit Apc10. Recovered peptides were tested for their effects on the replicative lifespan (RLS) of dividing cells, and the chronological lifespan (CLS) of nondividing cells. Several peptides increased the RLS of wild type cells, but only one peptide (C43-4) increased CLS, which requires the APC Cdh1 co-activator, but not the Cdc20 co-activator. In cells expressing C43-4, Clb1 and Mps1 levels remained low when aging quiescent cells were reintroduced back into the cell cycle. The addition of a synthetic C43-4 to aging quiescent cells increased CLS even when added at late stages of aging. Mutations to APC subunits and co-activators blocked the ability of C43-4 to extend CLS. Htz1, which shares sequence homology with C43-4, was found to suppress APC mutant phenotypes and stabilize Apc10. We individually mutated the 6 amino acids in C43-4 that were shared with Htz1; one of the mutants disrupted the C43-4-Apc10 2-hybrid interaction and abolished the C43-4-dependent increase in CLS. C43-4 action is evolutionarily conserved, as C43-4 increased *C. elegans* lifespan in a *daf-16*- and *aak-2*-dependent manner. Our results describe the discovery of an evolutionarily conserved translational peptide that is the first of its kind to specifically activate the anti-aging APC^Cdh1^ complex.

## INTRODUCTION

Loss of genomic stability and proteostasis are key aging hallmarks, observed from yeast to humans, that push quiescent cells (also referred to as G0 or stationary phase cells) into senescence (López-Otín et al. 2023). As genomic stability and proteostasis erodes in aging cells, DNA mutations and aneuploidy, as well as misfolded and aggregated proteins, begin to accumulate (Kennedy et al. 2014; Harkness et al. 2018; Guo et al. 2022; Sanada et al. 2025). The accumulation of cellular damage typically results in the transition from quiescence to senescence, followed by the release of molecules that damage neighboring cells, and finally, death through apoptosis. Senescence is a major driver of aging, with senescent cells accumulating in the aging population (Ovadya et al. 2018; Song et al. 2020). In recent decades, medical improvements and lifestyle changes have driven an increase in the mean human lifespan (Li et al. 2018; Olshansky et al. 2024). However, the extra years of life are often fraught with a progressive increase in disease (Niccoli and Partridge 2012), suggesting that without a means to prolong the life of quiescent cells and slow cellular aging, it is unlikely that the human health span will continue to increase. Therefore, to increase human health span, to live longer without disease, it is vital to discern how to protect quiescent cells from the inherent damage that occurs to DNA and proteins during aging.

Proteostasis, the combined processes that work together to maintain protein folding, degradation, and trafficking, is vital for the maintenance of cellular integrity (Morimoto and Cuervo 2014). The ubiquitin-proteasome system (UPS) orchestrates the selective degradation of key regulatory proteins (Kevei et al. 2017; Sampaio-Marques and Ludovico 2018) and plays a pivotal role in maintaining the appropriate complement of proteins in the cell. As the cell ages, proteostasis erodes, largely due to impaired proteasome- and autophagy-dependent removal of misfolded and aggregated proteins (Kaushik and Cuervo 2015; Akbergenovet al. 2025). Among the critical regulators of proteostasis during the cell cycle is the Anaphase Promoting Complex (APC), an essential and highly conserved ubiquitin ligase (E3) that targets protein substrates for degradation. The APC ensures proper exit from mitosis, entrance to G1/G0, and protection against chromosomal instability (Greil et al. 2022; Hu et al. 2022; Zhang et al. 2025). The significance of APC function extends beyond cell cycle control, as recent evidence indicates its involvement in stress response regulation during the yeast quiescent stationary phase and its potential role in aging processes (Postnikoff et al. 2012; Chen et al. 2012; Harkness 2018; Hu et al. 2022). Elevated levels of APC substrates have been observed in various cancers, underscoring the impact of APC dysregulation on disease and cellular aging (VanGenderen et al. 2020; Arnason et al. 2022; Jeong et al. 2023; Lubachowski et al. 2024; VanGenderen et al. 2026).

The APC takes the form of two temporally distinct complexes, APC^Cdc20^ and APC^Cdh1^ (APC^Cdc20^ in yeast, APC^CDC20^ in mammalian cells) APC^Cdc20^ is responsible for entrance to and transition through mitosis, while APC^Cdh1^ triggers mitotic exit and maintains the G1/G0 state until cells are ready for DNA replication (Greil et al. 2022; Hu et al. 2022; Zhang et al. 2025). Since APC^Cdh1^ is critical for entrance into G0 and maintenance of quiescent cells (Hu et al. 2011; Chen et al. 2012; Eguren et al. 2013; Lafranchi et al. 2014), we propose that activation of APC^Cdh1^ will prolong the quiescent state, delaying the transition into pathogenic senescence. APC^CDC20^ and APC^CDH1^ activity is tightly regulated by several competing intercellular factors, such as RAS/PKA, MAPK, PLK, and SIRT2 (Yamashita et al. 1996; Kotani et al. 1998; Zhou et al. 2016; Harkness 2018; VanGenderen et al. 2020). A major APC inhibitor is the Spindle Assembly Checkpoint (SAC) that sequesters Cdc20 away from the APC during prometaphase until all spindles are attached to kinetochores at the metaphase plate (Kapanidou et al. 2017). Once all chromosomes are attached to the metaphase plate, the SAC is satisfied and Cdc20 is released, allowing it to interact with the APC. Defects in SAC signalling and function are associated with cancer and aging (Brown and Geiger 2018). Current research indicates that premature activation of APC^Cdc20^ via SAC inhibition in cancer cells promotes rapid entry into mitosis, causing cells with high levels of aneuploidy to undergo mitotic catastrophe and apoptosis (Choi et al. 2017; Szymiczek et al. 2017; VanGenderen et al. 2020; Lubachowski et al. 2024). SAC inhibitors have been developed as anti-cancer therapeutics and are currently entering clinical trials (Lima et al. 2017). However, SAC inhibitors activate APC^CDC20^, but not APC^CDH1^. Interestingly, mutations to APC subunits have been found to generate resistance to SAC inhibitors *in vitro*, providing evidence that an active APC is required for these inhibitors to work (Thu et al. 2018).

While SAC inhibition is currently considered as therapy against cancer, activation of the proteasome is also considered to have therapeutic potential to delay aging and promote cell health (Upadhyay and Joshi 2024). Inhibition of the proteasome also targets cancer cells for death (Zhou et al. 2023), highlighting the fine line proteostasis plays in cancer cell growth. Based on observations that mammalian APC^CDH1^ contributes to tumor suppression (Sansregret et al. 2017; Garzón et al. 2017; Thu et al. 2018), and yeast APC^Cdh1^ is required for stress response and longevity (Harkness et al. 2004; Postnikoff et al. 2012; Menzel et al. 2014; Malo et al. 2016), we focused on the development of APC^Cdh1^ activators. To do so, we used a yeast 2-hybrid (Y2H) screen (VanGenderen et al. 2026) to discover small peptides that interacted with the APC subunit Apc10. We subcloned yeast-derived APC-binding peptide constructs into a human expression vector and discovered that one in particular, called C43-4, was capable of stalling MDR cancer growth *in vitro*, and in mice growing breast cancer cells *in vivo*. In the study described here, we characterized Apc10-binding peptides for those that could increase yeast lifespan and reverse impairment to activity APC as cells age. We show that C43-4 is the only peptide tested that could activate APC^Cdh1^.

## MATERIALS AND METHODS

### Yeast strains, media, and basic methods

*Saccharomyces cerevisiae* strains used in this study were all haploid, and are listed in **Table 1**. Yeast cells were cultured at 30°C in rich medium (YPD - 1% yeast extract, 2% peptone, and 2% glucose (dextrose) [Fisher Scientific, BD 242820]) or in synthetically defined (SD) medium (0.17% yeast nitrogen base without amino acids [Fisher Scientific, BD 233520], 0.5% ammonium sulfate, and 0.13% essential amino acids, with the carbon source maintained at 2%, either with 2% glucose, sucrose, galactose, or mixtures of sucrose and galactose, as indicated). Omission of specific amino acids provided selection pressure for the maintenance of transformed plasmids containing auxotrophic markers (plasmids used in this study are shown in **Table 2**). Complete media (CM) was the same as SD except all amino acids were included. Minimal media (MM) was prepared the same as SD, except with only 0.013% supplementation of necessary amino acids, using a ten-fold reduction in available amino acids to increase cellular stress and shorten lifespan. Selection of the KanMX module, used to replace wild type genes, required 0.2 mg/mL of geneticin (G418 sulfate; Gibco, 11811023) added to YPD. XL1-Blue chemically competent cells (Agilent #200249) were cultured in LB broth (0.5% yeast extract [Gibco, 212750], 1% tryptone [Gibco 211705], and 1% NaCl [Fisher, BP358-1]) for basic plasmid extraction from bacterial cells. To make solid media, 2% agar was added to the liquid medium prior to autoclaving. Molten agar was cooled to approximately 55-60°C (2% sugars were added to media) before pouring into petri dishes (VWR, 25384-302). For long-term storage, cells were grown to log phase in YPD, suspended in 1.5% glycerol, and stored at -80°C. Yeast transformations were performed using the LiAc method as previously described (Gietz and Schiestl 2007). Yeast genomic DNA extraction was conducted using a LiAc-SDS method according to a published protocol (Lõoke et al. 2011). Bacterial DNA preps, western blots, and PCR was performed according to previous methods (Malo et al. 2016). Replicative lifespan was conducted as published previously (Menzel et al. 2014; Postnikoff and Harkness 2014).

**Table 1.**
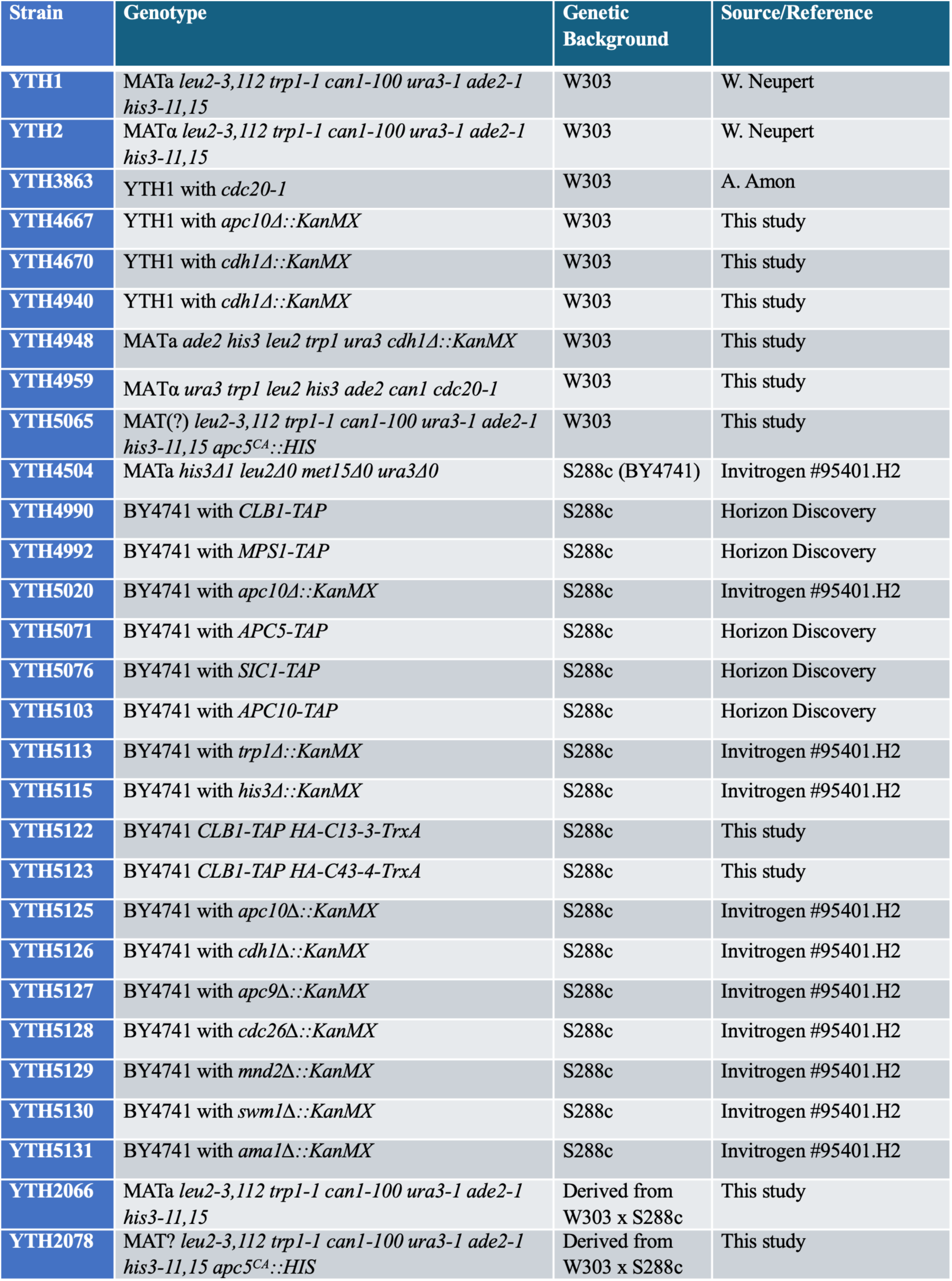
List of yeast strains used in this study.

**Table 2.** List of plasmids used in this study.

| Plasmid | Vector | Insert | Source/Reference |
| --- | --- | --- | --- |
| pJG4-5 | *2μ-TRP1* | *GAL1prom* | C.R. Geyer |
| pJG4-5-C2-4B | *2μ-TRP1* | *GAL1prom-HA-C2-4B* | VanGenderen et al. 2026 |
| pJG4-5-C3-1 | *2μ-TRP1* | *GAL1prom-HA-C3-1* | VanGenderen et al. 2026 |
| pJG4-5-C3-1B | *2μ-TRP1* | *GAL1prom-HA-C3-1B* | VanGenderen et al. 2026 |
| pJG4-5-C4-3 | *2μ-TRP1* | *GAL1prom-HA-C4-3* | VanGenderen et al. 2026 |
| pJG4-5-C9-5 | *2μ-TRP1* | *GAL1prom-HA-C9-5* | VanGenderen et al. 2026 |
| pJG4-5-C13-3 | *2μ-TRP1* | *GAL1prom-HA-C13-3* | VanGenderen et al. 2026 |
| pJG4-5-C43-4 | *2μ-TRP1* | *GAL1prom-HA-C43-4* | VanGenderen et al. 2026 |
| pRS416 | *CEN-URA3* |  | ATCC #87521 |
| pRS416-C43-4 | *CEN-URA3* | *GAL1prom-C43-4-HA* | This study |
| YEplac181 | *2μ-LEU2* |  | W. Neupert |
| YEplac181-C43-4 | *2μ-LEU2* | *GAL1prom-HA-C43-4* | This study |
| pYEX-4T-1 | *2μ-URA3* | *CUP1prom* | Clontech #6199-1 |
| pYEX-4T-1-APC5 | *2μ-URA3* | *CUP1prom-GST-APC5* | Clontech #6199-1 |
| pYEX-4T-1-APC10 | *2μ-URA3* | *CUP1prom-GST-APC10* | Clontech #6199-1 |
| pYEX-4T-1-HTZ1 | *2μ-URA3* | *CUP1prom-GST-HTZ1* | Clontech #6199-1 |
| pMWU-C43-4-GFP | pDEST-eft-3p | SV40-HA-TrxA-C43-4-GFP | This study |

### Yeast NaOH protein extraction

Prior to sample preparation, 4 X sample loading buffer was prepared (0.2 M Tris base [Sigma-Aldrich, 252859], 0.4 M DTT [Fisher, FERR0861], 277 mM SDS [BioShop, SDS999.1], 6 mM Bromophenol blue [Sigma-Aldrich, 114391], and 4.3 M glycerol [Fisher, BP229-1]), stored at - 20°C, and diluted to 1 X as needed. The absorbance at an optical density of 600 nm (OD_600_) of all yeast cultures was measured using a SmartSpec (Bio-Rad SmartSpec Plus Spectrophotometer), and 1-5 mL of culture was centrifuged at 5000 X g for 3 minutes. Up to 5 mL was taken if the initial OD_600_ reading was below 1.0 to have enough protein for extraction, dependent on strains grown and absorbance recorded. Supernatant was removed and pellets were resuspended in 500 μL of ice-cold 0.5 M NaOH (Sigma-Aldrich, SX0607N). Cell suspensions were incubated on ice for 10 min and centrifuged at 13,000 X g for 2 minutes at 4°C before removing the supernatant. Samples were resuspended in 1 X sample loading buffer as follows: 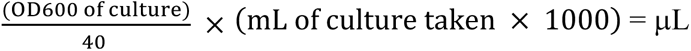 of 1 X sample loading buffer to add. Samples were frozen at -20°C overnight, and then thawed, mixed, and heated at 80°C for 10 min. After heating, samples were chilled on ice for 2 minutes and centrifuged at 13,000 X g for 5 minutes at 4°C before transferring protein-containing supernatant to a new tube. Samples were used for subsequent analysis and stored at -80°C.

### Western blotting

Westerns were performed according to our published methods (Menzel et al. 2014; Malo et al. 2016), with updates provided. 15 μL of protein lysates were separated through 10% SDS-PAGE gels with a BLUelf pre-stained protein ladder (FroggaBio, PM008-0500-G) in 1 X SDS running buffer. Electrophoresis was performed at 150 V for 60-90 minutes depending on the thickness of the gel cast. Total protein was visualized by placing the gel cast with 0.5% Trichloroethanol (TCE) on a benchtop UV transilluminator (VWR Compact UV Transilluminator) and irradiating for 1-2 minutes to induce covalent modification of tryptophan proteins to shift fluorescent emission to the visible range (Ladner et al. 2004). Gels were imaged with a Bio-Rad VersaDoc MP molecular imager to capture total protein loaded in each lane. Separated proteins on these gels were transferred for 30-50 minutes at 20 V to a nitrocellulose membrane (Bio-Rad, 1620115), which was preactivated by a 10-minute incubation in 1 X transfer buffer. After transfer, if transilluminated total protein load was not captured prior to transfer, membranes were submerged for 1-2 minutes in Ponceau S stain to visualize and scan total protein. Ponceau S stain was washed away with phosphate buffered saline with 1% Tween-20 (PBST), and membranes were blocked for 30-60 minutes on an orbital shaker (Corning LSE Platform Rocker) at room temperature (RT) with blocking buffer (5% skim milk in PBST). Blocked membranes were then submerged in primary antibodies against TAP (Thermo Fisher, CAB1001), HA (Abcam, ab9110), GAPDH (Proteintech, 60004-1-Ig), histone H3 (Proteintech, 17168-1-AP), Clb2 (Y180, Santa Cruz, SC-9071), GST (J90, described in Turner et al. 2010), or FITC (Proteintech, 80003-1-RR) diluted 1:1000 in blocking buffer (except GAPDH, which was diluted 1:10,000), and incubated for 3 hours at room temperature or overnight at 4°C. Following incubation, membranes were washed with PBST (0.1% Tween 20) for three 10-minute washes prior to incubation with HRP-conjugated secondary antibodies diluted 1:10,000 with blocking buffer for 30 minutes at RT (Rabbit HRP conjugated antibody: BioRad, 1706515; Mouse HRP conjugated antibody: BioRad, 1706516). Another three 10-minute PBST washes were performed before membranes were imaged with a Bio-Rad VersaDoc MP molecular imager, utilizing Western ECL substrate (Bio-Rad, 170-5060) to detect chemiluminescence. Images were analyzed and quantified using ImageJ software, performed by measuring the signal in a region of interest, with the background signal subtracted. Quantified protein abundance was normalized to total protein using either TCE or Ponceau S images, or using GAPDH as a loading control.

### Chronological Lifespan using methylene blue

All aging stationary phase cultures were initiated by seeding a single colony of yeast into 5 mL of YPD or SD medium for auxotrophic selection with overnight shaking at 30°C. Cultures were resuspended the next day into fresh media (typically 5-15 ml) in flasks with a liquid-air ratio of 1:4, with incubation at 30°C and vigorous shaking. When a smaller volume was needed, when using the C43-4 synthetic peptide, 2-5 ml cultures were maintained in 15 ml conical tubes or 14 ml snap tubes. Cells continued dividing until reaching peak cell density resulting in a post-diauxic shift, typically after 1-2 days. Flasks were left shaking to maintain this condition for 1-2 weeks, with aliquots removed every second day to measure cell viability using methylene blue (Bapat et al. 2006; Kwolek-Mirek et al. 2014). 10-20 μL of cells were removed each time and mixed 1:1 with 0.1% (w/v) methylene blue solution (0.7 g sodium acetate trihydrate, 0.3 g glacial acetic acid, 50 mg methylene blue [Fisher Scientific; M291-25], add water to 50 ml) before plating on a hemocytometer. Yeast cells contain an enzyme that reduces methylene blue; viable cells remain colourless, while dead yeast cells stain dark blue. Viability was calculated as:

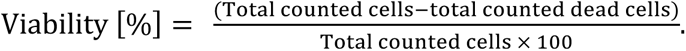

### Aging Arrest Assays

50 ml cultures were set up, with 6 mL collected every second day from the chronologically aging cells. Cells were resuspended in fresh YPD to allow cells to return to the cell cycle, in the presence of 15 μg/mL of nocodazole (Fisher Scientific; 48792810MG) to arrest cells in their first mitosis after being released from stationary phase. Cell cycle arrest was verified via microscopy, ensuring budding cells were visible before proceeding. The budding index was also determined. 2 mL of this culture was collected for the M phase time point, and the remaining 4 mL were washed with Ultrapure water (18.2MΩ-cm) before resuspending in fresh YPD medium, with the addition of new arresting agents. 2 mL were aliquoted with 60 μM α-factor (GenScript; RP01002) to arrest cells in the first “G1” phase after mitosis, or with 300 mM hydroxyurea (Fishert Scientific; AAA1083106) to arrest cells in the first “S” phase after mitosis. Cultures were left shaking for 2 hours at 30°C and spun down to pellet for lysate preparation. Lysates were prepared and separated by polyacrylamide gel electrophoresis for western blot analysis of protein abundance in each phase of the cell cycle for 8 days.

### Cycloheximide (CHX) Chase Assay

CHX inhibits ribosome function to observe intracellular protein stability or degradation (Schneider-Poetsch et al. 2010). Cells were inoculated in 5 mL of SD medium and grown overnight at 30°C shaking. Overnight cultures were diluted to an OD_600_ of 0.5 in 6 mL of fresh SD medium with 15 μg/mL of nocodazole to arrest cells in mitosis (approximately 2 hours). At the 2 hour time point, the t=0 sample was collected (samples arrested in mitosis). Samples were spun down and rinsed twice with Ultrapure water (18.2MΩ-cm) before resuspending in fresh SD media with 10 μg/mL of CHX. Aliquots were collected every 30 minutes for 90 minutes, freezing each pellet immediately after harvesting. Lysates were prepared and separated by polyacrylamide gel electrophoresis for western blot analysis of protein abundance at each time point.

### Integration of peptide constructs into the genome

To incorporate the HA-TrxA-C13-3 or HA-TrxA-C43-4 peptide constructs directly into the genome, two sets of PCR reactions were performed using Phusion™ High–Fidelity DNA Polymerase (Thermo, F530S). Two primers (listed in **Table 3**) were used on the Trp+ pJG4-5 vectors, which have the HA-TrxA-C13-3 or HA-TrxA-C43-4 constructs cloned into the multiple cloning site (MCS). The first integration primer (#1) was designed with an 18 bp sequence upstream of the TrxA peptide sequence and included a 60 bp overhang with homology to the *TRP1* promoter. The downstream primer (#2) contained 18 bp of the final TrxA gene including the stop codon, with another 60 bp overhang of homology 100 bp upstream of the LEU2 promoter. The next set of primers were designed for use with genomic DNA harboring the *LEU2* gene. The first primer in this set (#3) shared 18 bp of homology 100 bp upstream of the *LEU2* gene, with a 60 bp overhang of the final TrxA gene including the stop codon. The final primer (#4) shared 18 bp of homology to *LEU2* from the stop codon and had a 60 bp overhang downstream of *TRP1*. The schematic shown in **Supplemental Figure 1** outlines the plan for integration of these PCR fragments into the Clb1-TAP (YTH4990) strain that lack a functional *LEU2* gene (*leu2Δ0*). The integration fragments were used to incorporate a functional *LEU2* gene with the peptide construct into the *TRP1* locus, disrupting the *TRP1* gene.

**Table 3.**
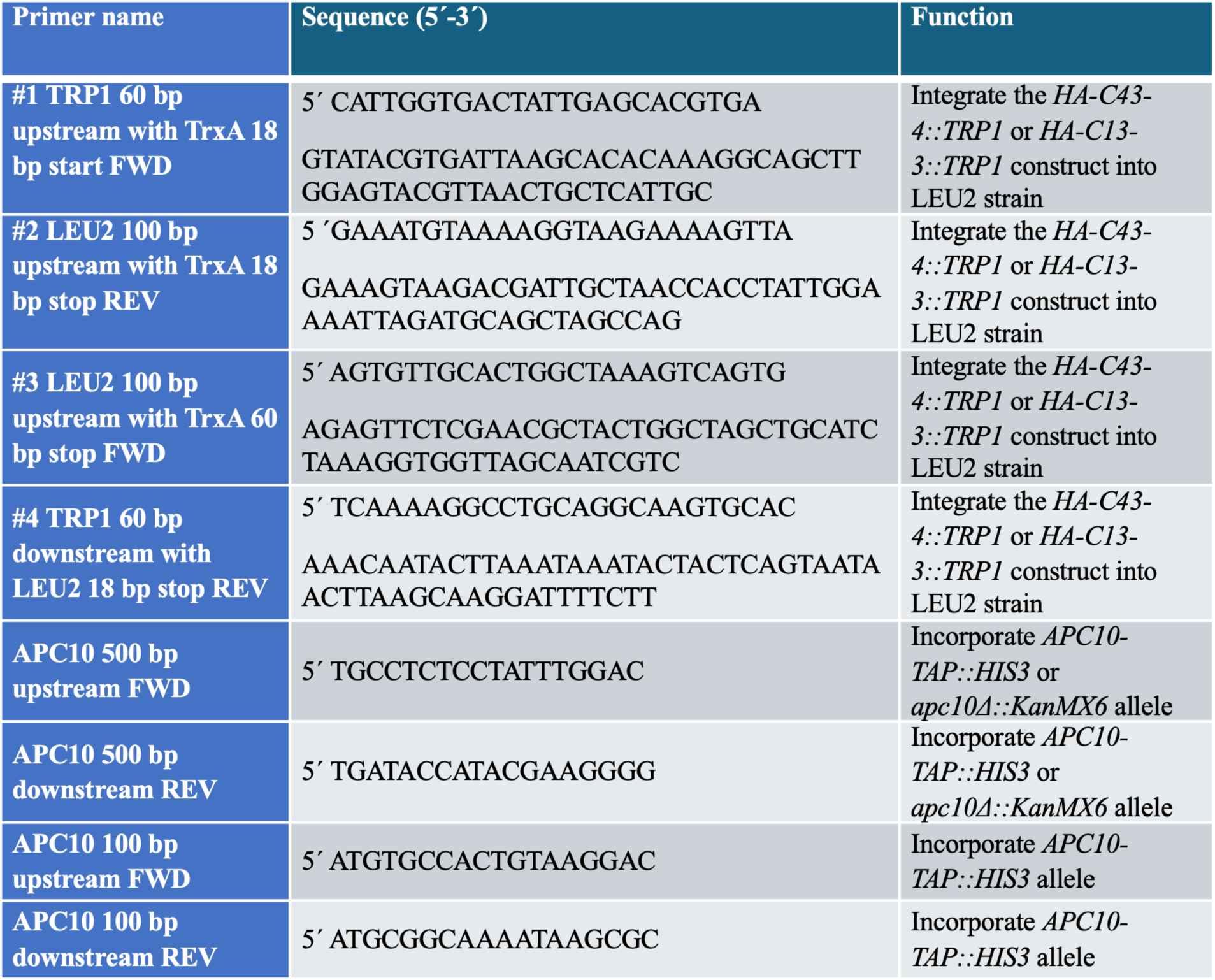
List of primers used in this study.

### Complex formation between FITC labelled peptides and TAT2

To allow complex formation between C43-4^FITC^ and the membrane permeable peptide TAT2 (peptides obtained from BioMatik), we first diluted 10 μL of a 10 μg/μL TAT2 solution in 90 μL water. 5 mg of the powdered C43-4 peptide was then resuspended in 5 mL water, with 10 μL of that solution diluted with 90 μL of water. We then mixed 10 μL of the diluted TAT2 and peptide solutions, with 80 μL water added. Next, we incubated the mixture at room temperature for 15 minutes for complex formation. 100,000 yeast cells were then pelleted, leaving 100 μL of supernatant. 100 μL of the peptide/TAT solution was added, gently mixed, then incubated for 15 minutes. Cells were then used for experiments.

### Nuclear fractionation in yeast

Nuclear fractionation was conducted to visualize the localization of the HA-TrxA-C43-4 plasmid-expressed peptide and the synthesized C43-4^FITC^ peptide. Yeast cultures were grown overnight at 30°C in YPD, then pelleted and washed twice using spheroplasting buffer (1.2 M sorbitol [Sigma-Aldrich; S1876-500G], 0.1 M KH_2_PO_4_ [Sigma-Aldrich; P5379-100G]). The cells were resuspended in spheroplasting buffer with 0.2 mg/mL zymolyase (Amsbio, 120491-1) and incubated at 30°C for one hour to rupture the cell wall. Cells were monitored for spheroplasting using a 100 X lens (Olympus CX41). The spheroplasted cells were then pelleted, washed, and resuspended in 1 M sorbitol, with the addition of NP-40 lysis buffer (20 mM Tris-HCl, pH 7.4, 20 mM NaCl, 10 mM MgCl_2_, 1 mM EDTA, 1% NP-40) containing protease and phosphatase inhibitor tablets (Sigma-Aldrich, 04693159001, PHOSS-RO). The cells were then left to lyse on ice for 15 minutes. The culture was centrifuged at 500 X g for 5 minutes at 4°C. The protein-containing supernatant was then centrifuged at 13,000 X g for 15 minutes at 4°C. The subsequent supernatant consisted of the cytoplasmic fraction, while the final pellet was resuspended in lysis buffer as the nuclear fraction. All samples were then sonicated (Branson Sonifer 250) at 30% duty pulse cycle for 5 pulses for additional lysis, with protein concentrations determined using a Bradford assay (Bio-Rad, 5000006). Samples were then heated at 80°C for 5 minutes with 1 X sample loading buffer (5 X loading buffer [250 mM Tris HCl, pH 6.8, 10% w/v SDS], 25% v/v 2-mercaptoethanol, 50% v/v glycerol, and 0.5% w/v bromophenol blue).

### C. elegans strains

The following strains were used: N2 bristol wildtype, MWU96 *cwwIs3*[*eft-3p-TrxA(APC)::GFP; myo-2p::tdTomato*], MWU128 *cwwIs3*; *daf-16(mu86),* CF1038 *daf-16(mu86)*, MWU 143 *cwwIs3; aak-(ok524)*, RB755 *aak-2(ok524)*, MWU146 *cwwIs3; akt-1(ok525)*, RB759 *akt-1(ok525)*. CF1038, RB755, and RB759 were outcrossed with N2 six times prior to generating MWU128 and MWU143 for the lifespan assay. Worms were cultured at 20°C using standard methods as described (Brenner 1974).

### *C. elegans* transgenic construction

To generate the C43.4::GFP strain, a synthetic DNA fragment containing the SV40 NLS:HA:TrxA:C43.4 sequence was first synthesized by Genestrand (Eurofin Genomics) and fused with GFP using the PCR methods as described by (Hobert 2002). The GFP fusion fragment was amplified with *attB1* and *attB2* overhangs and cloned using the Gateway method into the pDEST-eft-3p plasmid that drives ubiquitous gene expression, flanked by the *unc-54* 3’ UTR. The constructed plasmid was microinjected into *C. elegans* at 25 ng/µL along with the *myo-* 2::tdTomato co-injection marker (2 ng/µL). Transgenic strains expressing the extrachromosomal array were then integrated to generate stable expression using the UV irradiation technique, followed by six rounds of outcrossing with the N2 wild type prior to use.

### *C. elegans* lifespan assays

Lifespan assays were conducted as previously described (Tabarraei et al. 2023). *C. elegans* were synchronized using the hypochlorite treatment and grown on *HT115* empty vector *E. coli* as the food source at 20°C. Worms were picked manually to segregate from the offspring during the reproductive period. Day one is considered the first day of adulthood. Three independent trials were performed for all lifespan assays with statistics for each trial listed in **Supplemental Tables 7-10**. Lifespan data was analyzed using the log-rank test using the OASIS2 software (Han et al. 2014; 2016).

### Statistical analyses

Statistical data calculations for RLS experiments were determined using OASIS2, as described previously (Gutierrez et al. 2026). Log-rank tests were applied. Area under the curve (AUC) calculations were performed for CLS experiments for each strain ± SEM and analyzed with a one-way ANOVA and Dunnett’s multiple comparisons *post-hoc* test using GraphPad Prism. Spot dilution normalization of the 100-fold dilution (third column) for each strain ± SEM was determined using a one-way ANOVA and Dunnett’s multiple comparisons *post-hoc* test, as described previously (Petropavlovskiy et al. 2020). One- or two-way ANOVA tests were used to compare means of unmatched groups. Multiple comparisons tests were performed *post-hoc* as indicated to calculate *p*-values and assess significance between strains or treatments. Tukey’s *post-hoc* testing was used after one- or two-way ANOVA tests to identify which pairs of group means were different. Dunnett’s *post-hoc* testing was used to compare multiple treatment groups to a single control group. Šídák’s *post-hoc* testing was used to compare specific and independent group means. Significance is denoted as ‘ns’ for non-significant, *p*<0.05*, *p*<0.01**, *p*<0.001***, and *p*<0.0001****. Values represent ± standard error of the mean (SEM).

## RESULTS

### The APC loses function in aging quiescent yeast cells

Mutations to subunits of the Anaphase Promoting Complex (APC) exhibit short replicative (APC^Cdc20^- and APC^Cdh1^-dependent) and chronological (APC^Cdc20^-dependent only) lifespans (Harkness et al. 2004). Therefore, we tested whether wild type (WT) APC^Cdh1^ loses function in aging quiescent yeast cells. This is important as once cells enter quiescence, they can accumulate damage that eventually leads to senescence, a major hallmark of aging. Therefore, if APC^Cdh1^ activation can prolong the health of quiescent yeast cells via maintenance of repair pathways (APC^CDH1^ is associated with DNA repair and maintenance of genome stability; García-Higuera et al. 2008; Wäsch et al. 2010; Greil et al. 2016; Ha et al. 2017; Harkness 2018), it is expected that APC^Cdh1^ activation may extend the metabolic activity of quiescent cells, thereby delaying senescence and aging. To test this idea, we grew WT yeast cells expressing endogenous TAP-tagged APC substrates (Clb1-TAP and Mps1-TAP) to stationary phase, a nondividing quiescent state, and then aged them chronologically. Clb1 and Mps1 are targeted by APC^Cdc20^ during mitosis, and then APC^Cdh1^ during G1 (Baumer et al. 2000; Cui et al. 2010). As a control, we used TAP-tagged Sic1, which is targeted by the Skp, Cullin, F-box (SCF) ubiquitin ligase at the G1/S junction for degradation (**Figure 1A)** (Orlicky et al. 2003). We ensured that the APC functioning in these replicating cells was aged by arresting chronologically aging cells in the first M and G1 phases they encountered after being released into fresh YPD media. If the APC loses function in aging cells, then the APC^Cdc20^ substrates would not be as effectively targeted and accumulate in the subsequent G1. On the other hand, Sic1, targeted by the SCF at the end of G1, should remain low during S phase in aging cells (**Figure 1B**). We show that in our hands, Clb1-TAP and Mps1-TAP accumulate in M phase in log phase cells, and are degraded by G1, whereas Sic1-TAP is highest in G1 and removed at S, as expected (**Figure 1C**). We grew cells expressing the 3 TAP-tagged proteins to stationary phase, which relies on APC^Cdh1^ (Skaar and Pagano 2008; Cappell et al. 2016), removed samples every other day for 8 days, and then allowed them to re-enter the cell cycle in YPD liquid media supplemented with nocodazole. Once a mitotically arrested sample was removed, nocodazole was washed away with α−factor or hydroxyurea added to arrest the cells in G1 or S, respectively. The APC substrates Clb1 and Mps1 accumulated at mitosis in the aging cells, while Sic1 accumulated at G1 (**Figures 1D-1F**). We observed both Clb1-TAP and Msp1-TAP accumulated over the 8-day time course in G1 as the quiescent cells chronologically aged (**Figures 1D, 1E**), while protein levels in mitotically arrested cells remained stable. The G1:M ratio increased over time in aging cells (**Figures 1D**, **1E**), supporting our hypothesis that the APC loses function with age. Opposed to this, our results show that the Sic1-TAP levels remained low during S phase in the chronologically aging cells (**Figure 1F**). This indicates that while the APC appears to lose function in aging cells, targeting of Sic1 by the SCF is not impaired with aging.

**Figure 1.**
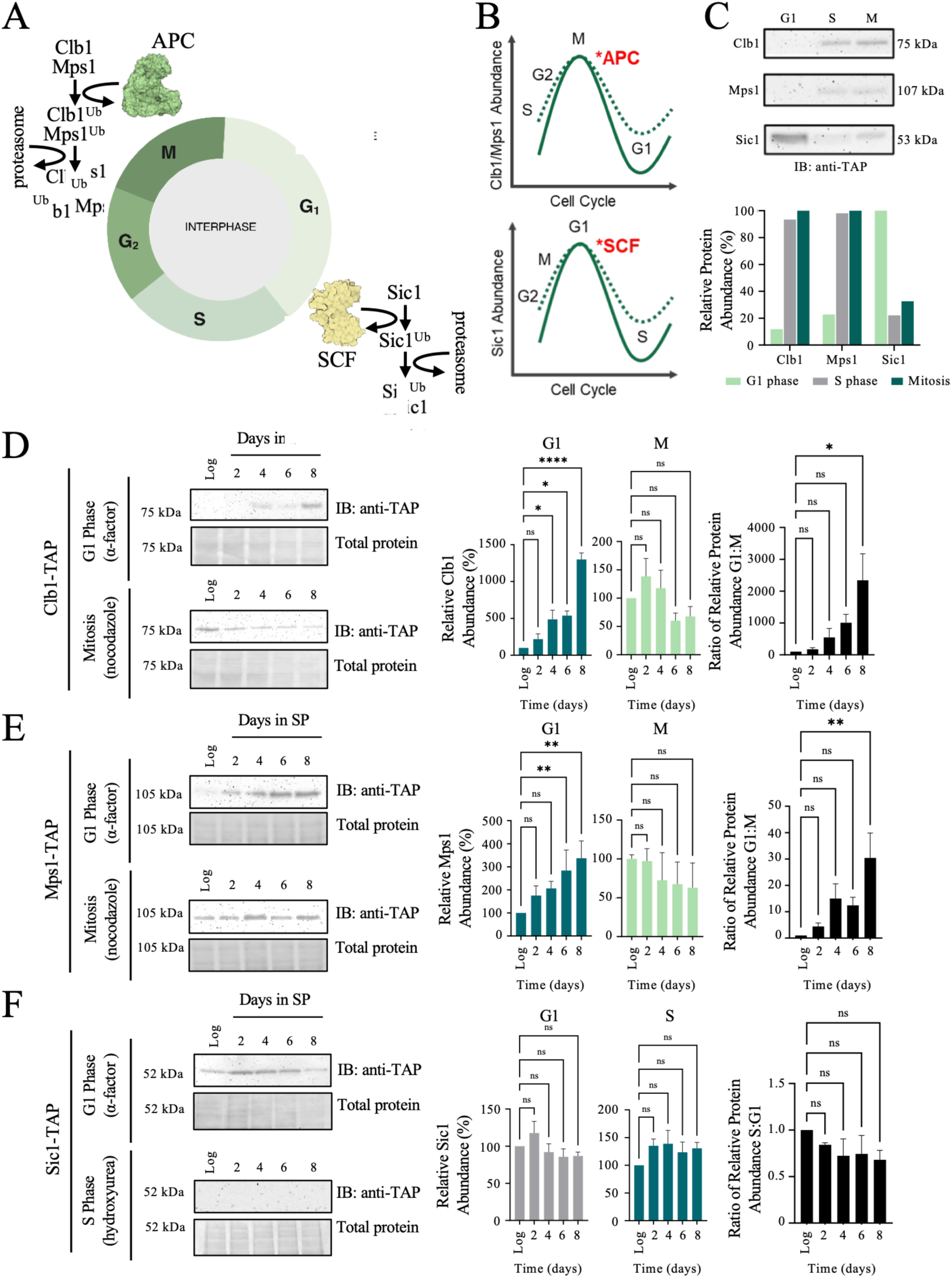
APC function decreases in aging cells. **(A)** Schematic of APC and SCF activity during the cell cycle, with examples of substrates targeted for ubiquitination at these time points. **(B)** Typical abundance of selected mitotic APC (top) or G1 SCF (bottom) substrates during the cell cycle in young, healthy cells when effectively targeted for ubiquitination. Dotted lines are expected changes in abundance of G1 APC (top) or S phase SCF (bottom) substrates during the cell cycle in chronoslogically aged cells when ineffectively targeted for ubiquitination. **(C)** Representative western blot analyses on overnight samples of young, log phase cells arrested with α-factor (G1), hydroxyurea (S), or nocodazole (M) to determine normal accumulation of proteins during the cell cycle. n=1. Western blots were normalized to total protein load (Ponceau S stained membranes) before each strain was normalized to M phase (100%), excluding Sic1-TAP which was normalized to G1 (100%). Uncropped blots are shown in **Supplemental Figure 2**. Western blot analyses were performed on lysates harvested from Clb1-TAP **(D)**, Mps1-TAP **(E),** or Sic1-TAP **(F)** cells in log and stationary phase. The budding index is shown in **Supplemental Figure 3** and full uncropped blots are shown in **Supplemental Figure 4**. Chronologically aging cells in stationary phase were resuspended in nutrient-rich media, supplemented with α−factor, hydroxyurea, or nocodazole to arrest cells in G1, S, or M phase, respectively, to determine trends in protein accumulation. Densitometry was used to normalize proteins to the Ponceau S stained membranes to determine relative abundance in the indicated cell cycle stages to determine G1:M ratios for Clb1 and Mps1, and S/G1 ratios for Sic1. Values are presented as mean ± SEM for n=3 and were analyzed with a one-way ANOVA and Dunnett’s multiple comparisons *post hoc* test where *p*<0.05*, *p*<0.001***, and *p*<0.0001**** between log phase and each day of stationary phase. Normalized graphs were used to compare relative ratios between cell cycle phases over time (black bar graphs, right), analyzed with a one-way ANOVA and Dunnett’s multiple comparisons *post hoc* test where *p*<0.05*.

### Treatment of aging dividing and nondividing cells with the small chemical APC activator M2I-1 increases RLS, but does not increase CLS

To further validate whether the APC is required for lifespan, we performed CLS assays using a methylene blue method to compare how long APC mutants can remain metabolically active during stationary phase. In all cases, the APC mutants *apc5^CA^*, *apc10Δ*, *cdc20-1*, and *cdh1Δ*, had a significantly shorter CLS (**Figure 2A**; stats shown in **Supplemental Figure 5A**). We previously showed that APC subunit mutants had a short CLS and RLS (*apc5^CA^*, *apc9Δ*, *apc10Δ* and *cdc26Δ*; see **Table 1** for the strain list; Harkness et al. 2004). *APC9*, *APC10*, *CDC26*, and *CDH1* are not essential genes and can be deleted (denoted by Δ), whereas *APC5* and *CDC20* are essential. The *apc5^CA^* mutant was isolated in a screen for chromatin assembly (CA) mutants (Harkness et al. 2002). The *apc5^CA^*allele contains a deletion of nucleotides 36 and 37, generating a novel amino acid sequence after amino acid 12, with a stop codon 18 amino acids later, producing a temperature-sensitive phenotype at 37°C. When the *apc5^CA^*allele was TAP tagged on the C-terminus, it resulted in a shorter, weakly expressed protein (Menzel, J. MSc thesis, 2014), indicating the *apc5^CA^* phenotype was likely due to translational slippage. The *cdc20-1* allele was isolated by the Amon lab (Visintin et al. 1997). Here, we show that both APC co-activators, Cdc20 and Cdh1, are required for normal CLS (**Figure 2A**).

**Figure 2.**
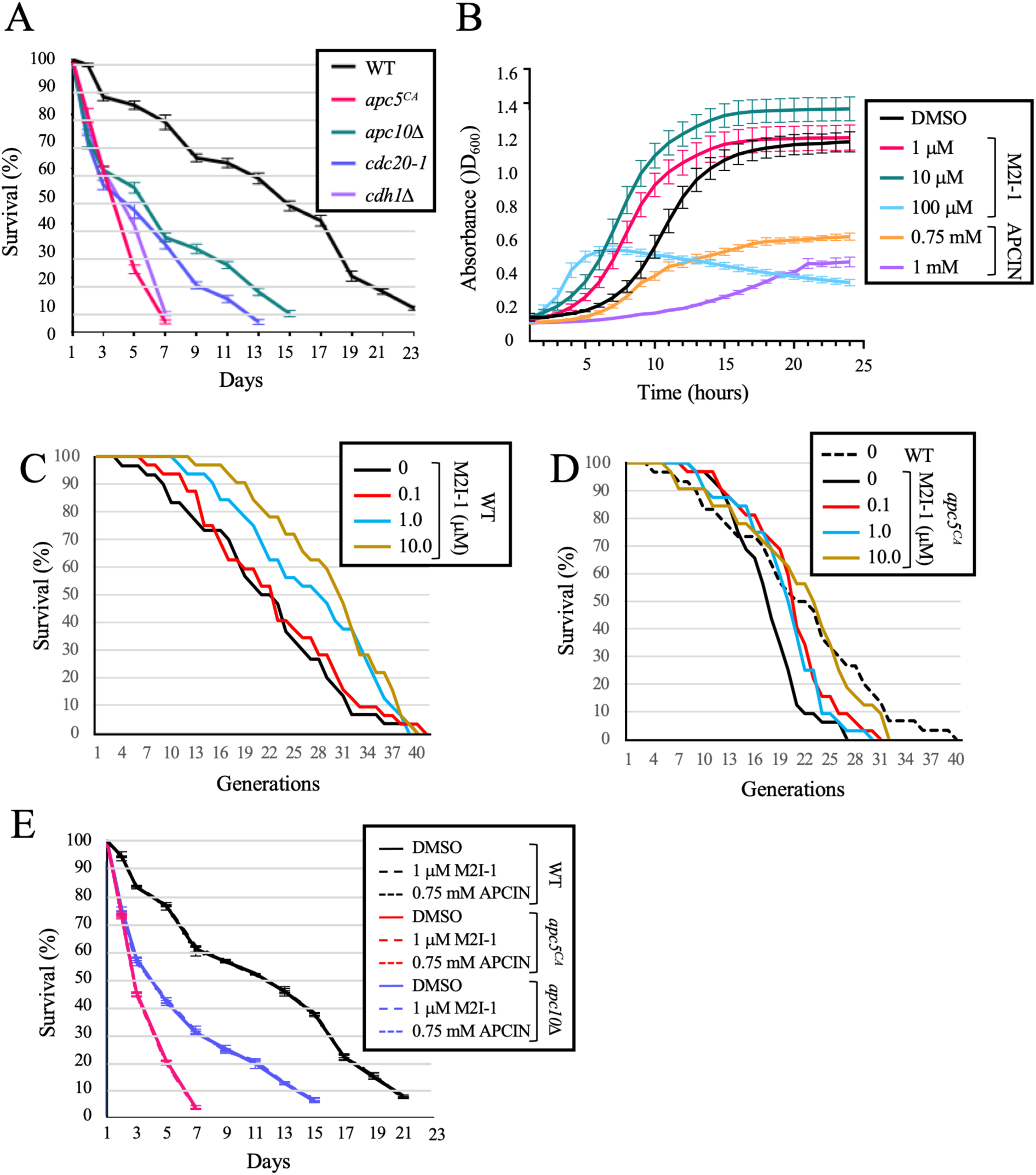
APC activation increases growth rate and RLS, but not CLS. **(A)** Chronological lifespan assays of WT (YTH2) and W303-based strains harboring APC mutants *apc5^CA^*(YTH5065), *apc10Δ* (4667), *cdc20-1* (YTH3863), and *cdh1Δ* (4670), compared in SD media. Survival was assessed using methylene blue cell counts every second day starting Day 1 until survival reached <10%. Values are presented as mean ± SEM for n=3. The quantitation is shown in **Supplemental Figure 5A**, with statistical analyses presented in **Supplemental Figure 5B**. The CLS for all APC mutants was significantly different from WT, with *p*<0.0001****. **(B)** 96-well growth assays were performed using YTH1 W303-based cells treated with the doses of the APC activator M2I-1 and the APC inhibitor APCIN shown, in SD media supplemented with 2% sucrose. 1% DMSO was used as the vehicle control, which corresponded to the concentration of DMSO in the 100 μM M2I-1 sample. The OD_600_ reading was measured every hour for 24 hours. Values are presented as mean ± SEM for n=4. Quantitation and significance are shown in **Supplemental Figure 5C**. The growth differences in the presence of M2I-1 and APCIN were statistically significant, with *p*<0.0001****. **(C)** Replicative lifespan of WT (YTH2066) and **(D)** *apc5^CA^* (YTH2078) cells exposed to the doses of M2I-1 shown. Cells were maintained on YPD plates supplemented with the appropriate M2I-1 doses. Statistical data is shown in **Supplemental Tables 1 and 2**. **(E)** W303-based WT (YTH1), *apc5^CA^* (YTH5065), and *apc10Δ* (YTH4667) strains were treated with the indicated doses of M2I-1 or APCIN, grown in SD medium with 2% sucrose. 1% DMSO was used as the vehicle control. Cells were grown to stationary phase and then chronologically aged. Survival was assayed by methylene blue cell counts every second day starting Day 1 until survival reached <10%. Values are presented as mean ± SEM for n=3 for each strain and condition. The treatments had no effect on CLS, therefore the overlap of the 3 CLS experiments for each strain looked like a single CLS experiment. Quantitation and significance is shown in **Supplemental Figure 5D**. The M2I-1 and APCIN CLS curves were not significantly different from the controls.

To test if the lifespan defects in APC mutants are due to a loss of APC function, we treated cells with increasing doses of M2I-1, thereby prematurely activating the APC (Kastl et al. 2015). First, we determined if chemical APC activation or inhibition impacted the growth rate of cells. We observed that 1 and 10 μM M2I-1 increased the growth rate of cells, while the higher dose of 100 μM impaired cell growth (**Figure 2B**; stats shown in **Supplemental Figure 5C**). We predicted that if APC activation increased cell growth, then APC inhibition, using APCIN (APC inhibitor) (Sackton et al. 2014), a chemical that blocks the ability of Cdc20 to interact with the APC, should decrease the growth rate of cells. We observed that 0.75 and 1 μM APCIN impaired cell growth (**Figure 2B**; stats shown in **Supplemental Figure 5C**). These observations suggest that increasing APC activity using M2I-1 at 1 and 10 μM increases yeast growth rate.

To test the effects of M2I-1 on RLS, we maintained mother cells on plates with increasing doses of M2I-1 (0.1, 1.0, and 10.0 μM). We found that treatment of WT cells with M2I-1 increased the mean RLS by roughly 50% at the highest dose used (**Figure 2C**; statistical analyses and *p*-values shown in **Supplemental Table 1**). Similarly, when the RLS of *apc5^CA^*cells was determined following increased exposure to M2I-1, it was observed that M2I-1 restored *apc5^CA^* short lifespan back to WT levels (**Figure 2D**; statistical analyses and *p*-values shown in **Supplemental Table 2**). Next, we assessed whether M2I-1 could also increase the CLS of quiescent WT and APC mutants. Only APC^Cdh1^ is present in stationary phase cells (Skaar and Pagano 2008; Cappell et al. 2016), whereas both APC^Cdc20^ and APC^Cdh1^ are required in cycling cells. We observed that 1 μM M2I-1 had no effect on the CLS of the cells (**Figure 2E**; quantitation shown in **Supplemental Figure 5D**). APCIN also had no impact on the CLS of WT and mutant cells. This is consistent with both M2I-1 and APCIN only impacting cycling cells, through interactions with Cdc20, and not Cdh1 (Sackton et al. 2014; Li et al. 2019).

### Characterization of peptides that bind to Apc10

To identify a method to activate APC^Cdh1^, we characterized the phenotypes of peptides expressed from the *E. coli* Thioredoxin A (TrxA) backbone that bound to the APC subunit Apc10 using a 2-hybrid screen (VanGenderen et al. 2026; **Figure 3A**; see **Table 2** for plasmids used). One of these peptides, C43-3, resensitized drug-resistant breast cancer cells to therapy, both *in vitro* and *in vivo*. First, a select group of peptides were expressed in W303-based wild type cells (**Figure 3B**). The transformed cells were spot diluted onto SD -Trp media supplemented with either 2% sucrose or 1% galactose/1% sucrose to induce the TrxA-peptide constructs that were under the control of the *GAL1* promoter, and grown at 30°C for 3 days. We chose to use sucrose as a non-inducing agent, as this sugar has less impact on galactose promoter repression than glucose (Neumann and Lampen 1967; Johnston et al. 1994). We observed that when induced with 1% galactose, C9-5, C13-3, and C43-4 slowed the growth of cells (**Figure 3B**; quantified in **Figure 3C**). Next, we used a plate reader with cells growing in 96-well plates, where the OD_600_ was measured every hour for 24 hours to assess whether the peptides shown above continued to slow growth in liquid media (**Figure 3D**). Peptide C9-5 slowed the growth to a greater extent than C13-3 and C43-3 when expressed with 1% galactose. We prepared lysates from the cells and performed western blots against the HA motif cloned into the TrxA-peptide construct. The peptides were all expressed, as expected (**Figure 3E**). Peptides C9-5, C13-3, and C43-4 were cloned in-frame with TrxA, generating larger bands than C2-4B, which was not cloned in-frame, creating an early stop codon rather than the C-terminal half of TrxA.

**Figure 3.**
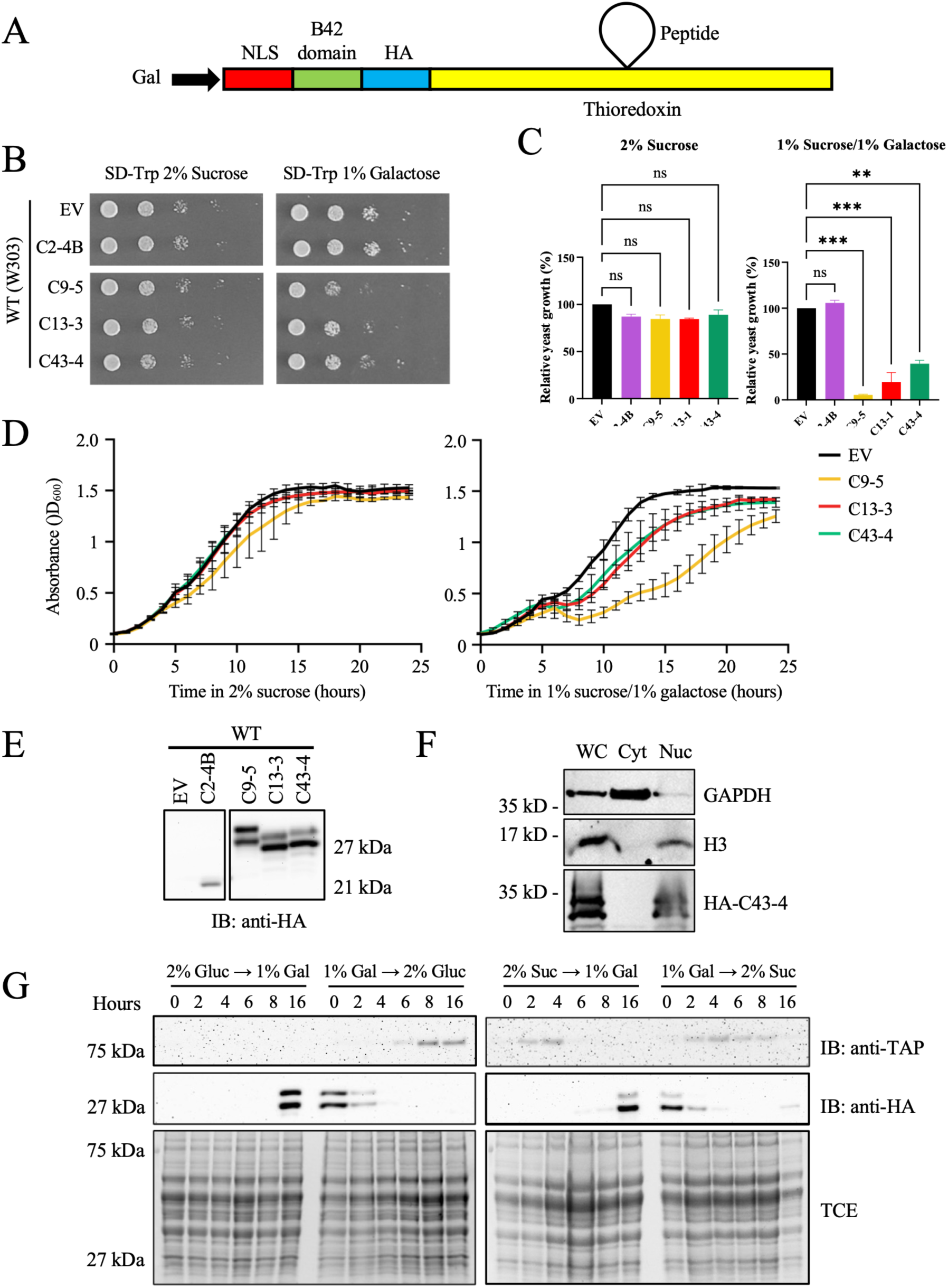
Phenotypic analyses following induction of the different peptide constructs. **(A)** Structure of the Thioredoxin A (TrxA) peptide display cassette. The peptide was cloned into the TrxA backbone, which was fused to a nuclear localization signal (NLS), a transcriptional transactivation domain (B42) and an HA tag, all under the galactose-inducible promoter, as previously described (VanGenderen et al. 2026). **(B)** Spot dilution assay for the WT W303-based strain (YTH1) transformed with the pJG4-5 empty vector (EV) and 4 different peptide constructs. Cells were cultured overnight, adjusted based on optical density, with ten-fold serial dilutions in 5 μl then spotted onto SD -Trp solid agar plates containing the indicated concentrations of carbon sources to induce peptide expression. Plates were incubated at 30°C for three days before imaging. Two sets of independent cultures were tested, and selected representative images are shown for n=2. **(C)** Normalization of 100-fold dilution (third column) for each strain ± SEM were analyzed with a one-way ANOVA and a Dunnett’s multiple comparisons *post-hoc* test. No significant differences were observed between the EV control strains and peptides in 2% sucrose, whereas in galactose, the differences between the EV control and C9-5, C13-3 and C43-4 was significant; *p*<0.01** and *p*<0.001***. **(D)** Growth curves of the YTH1 strain transformed with an EV control or the 4 different peptide constructs shown above, grown in 2% sucrose (left) or 1% sucrose/1% galactose (right) to induce peptide expression. Cells were cultured overnight, adjusted based on optical density, and maintained in a 96-well plate for OD_600_ readings every hour for 24 hours using a microplate reader. Values are presented as mean ± SEM for n=3. Curves were analyzed with a two-way ANOVA and a Dunnett’s multiple comparisons *post-hoc* test where no significance was observed between strains in 2% sucrose, and *p*<0.01** between the EV and C9-5 peptide in 1% galactose/sucrose. **(E)** Western blot analysis of HA-tagged peptide constructs to confirm yeast transformations; representative lysates harvested from cells grown in 1% galactose/sucrose are shown. Uncropped blots shown in **Supplemental Figure 6A**. **(F)** Clb1-TAP cells expressing an integrated TrxA-HA-C43-4 (also contains an NLS) were grown overnight in 2% galactose to induce the cassette. Cells were fractionated into nuclear and cytosolic lysates. Whole cell (WC), cytosolic (Cyt), and nuclear (Nuc) fractions were assessed by western blots using antibodies against GAPDH, histone H3, and HA. Uncropped blots shown in **Supplemental Figure 6B**. **(G)** Western blot analyses of Clb1-TAP cells with galactose-inducible HA-TrxA-C43-4 integrated into the genome, grown initially overnight in 2% glucose (left) or 2% sucrose (right). The following day, samples were resuspended in 2% galactose (left) or 1% galactose/1% sucrose (right) supplemented media. Samples were collected every 2 hours for 8 hours, and another sample was taken the next morning after 16 hours. Cells were then washed and resuspended in 2% glucose or 2% sucrose with samples again taken every 2 hours for 8 hours, and the last sample taken after 16 hours. Lysates were prepared for western blot analysis using anti-HA and anti-TAP to detect HA-TrxA-C43-4 and Clb1 abundance, respectively. Uncropped blots are shown in **Supplemental Figures 6C and 6D**.

If the peptides do in fact activate the APC, then the peptides should be localized to the nucleus. We used HA-TrxA-C43-4 to determine whether the peptides could be localized to the nucleus. Therefore, we fractionated cells expressing HA-TrxA-C43-4 integrated into the genome of Clb1-TAP cells (see **Supplemental Figure 1** for the integration scheme) grown overnight in 2% galactose to recover nuclear and cytoplasmic fractions. We showed that histone H3 and GAPDH signals were exclusively in the nucleus and cytoplasm, respectively. As expected, HA-TrxA-C43-4 was exclusively in the nucleus (**Figure 3F**). To confirm whether C43-4 activated the APC, we used Clb1-TAP cells with the integrated HA-TrxA-C43-4 cassette. Upon APC activation, we expect that Clb1-TAP protein levels in cells will decrease, as shown previously with chemical activation of the APC in human cells (Arnason et al. 2022; Lubachowski et al. 2024; VanGenderen et al. 2026). Cells were grown overnight in complete SD media supplemented with 2% sucrose, then washed and resuspended in 1% galactose/1% sucrose to induce C43-4, with samples taken as shown. The cells were once again washed and resuspended in 2% sucrose to slow C43-4 overexpression. Samples were again removed over time. We observed that C43-4 was expressed over time with galactose, with levels going down when galactose was removed (**Figure 3G**). In support of C43-4 activating the APC, Clb1-TAP levels were reduced upon C43-4 induction and returned once C43-4 induction was stopped.

### C43-4 slows cell growth in a Cdh1-dependent manner

Consistent with increased APC activity slowing yeast growth, we previously reported that overexpression of *APC5*, or the *FKH1* and *FKH2* members of the APC stress response network, reduced yeast cell growth (Harkness et al. 2004; Postnikoff et al. 2012). We further characterized peptide effects on yeast growth by expressing C2-4B, C3-1, C3-1B, C4-3, C9-5, C13-3, and C43-4 in the WT W303-based genetic background (**Supplemental Figure 7A**), or C2-4B, C9-5, C13-3, and C43-4 in the S288c-based genetic background (**Supplemental Figure 7B**). The transformants were spot diluted onto solid SD -Trp media supplemented with either 2% glucose or 2% galactose. To test effects of the peptides on heat stress, the plates were grown at 30°C and 37°C for 2-3 days, and then scanned. C2-4B, C3-1, C3-1B, and C4-3 had little impact on glucose-or galactose-grown cells at 30° or 37°C (**Supplemental Figure 7**). C13-3, C9-5, and C43-4 slowed the growth of both W303- and S288c-based cells, with the effect more prominent with 2% galactose induction (compare spots shown in **Supplemental Figure 7A** with those in **Supplemental Figure 7C**; see **Supplemental Figure 7D** for quantification of the spots shown in **Supplemental Figure 7C** on 1% galactose). We speculate that C9-5, C13-3 and C43-4 may be increasing APC activity to high levels, which we have shown may slow cell growth (Harkness et al. 2004). This was also observed with M2I-1, which increased growth at lower doses, but decreased it at higher doses (**Figure 2B**).

To test whether the APC is required for the peptide slow growth phenotype, we transformed W303-based WT and *apc5^CA^* cells with the EV or TrxA-HA-C43-3, and spot-diluted the cells onto SD - Trp plates supplemented with either 2% sucrose or 1% galactose/1% sucrose (**Figure 4A**). C43-4 continued to slow the growth of *apc5^CA^*cells, but not as dramatically as observed in WT cells (**Figure 4A**, quantified in Figure **4B**), supporting our observations that C43-4 requires a functional APC to assert its activity. To gain direct evidence that C43-4 requires either the Cdc20 or Cdh1 co-activators to activate the APC, we first transformed W303-based cells harboring a *CDH1* deletion with C2-4B, C13-3, C43-4, and the EV control (**Figure 4C**). The cells were spot diluted onto SD -Trp media supplemented with either 2% glucose or 2% galactose, with the plates grown at 30° or 37°C. WT cells expressing C13-3 or C43-4 grew slowly when induced with 2% galactose, whereas C2-4B expressing cells grew at the same rate the EV control cells. The effect for C43-4 was enhanced when grown at 37°C on 2% galactose plates (**Figure 4C**; quantified in **Figure 4D**).

**Figure 4.**
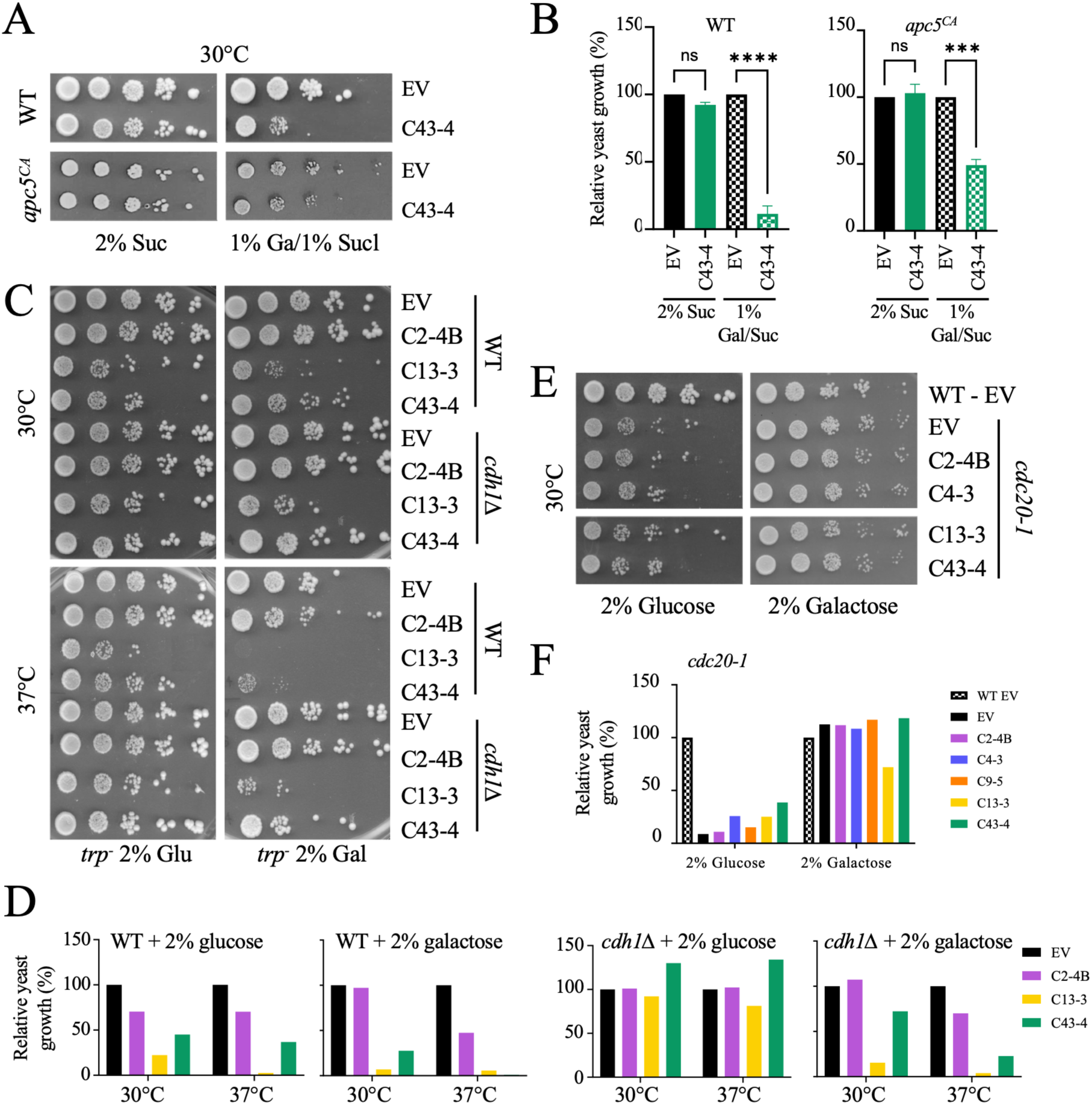
C43-4 growth phenotypes require both *CDC20* and *CDH1*. **(A)** WT (YTH1) and *apc5^CA^* (YTH5065) cells expressing either the EV or HA-TrxA-C43-4 were cultured overnight, adjusted based on optical density, then ten-fold serial dilutions were spotted onto SD -Trp plates supplemented with the sugars shown. Plates were incubated at 30°C for three days before imaging. Two sets of independent cultures were tested, and selected representative images are shown for n=2. **(B)** Normalization of 100-fold dilution (third column) for each strain ± SEM were analyzed with a one-way ANOVA and a Dunnett’s multiple comparisons *post-hoc* test where no significance was observed between the EV control strains and different peptides in 2% sucrose, and *p* < 0.001***, and p < 0.0001**** in 1% galactose. (C) Spot dilutions of W303 WT (YTH1) and *cdh1Δ* (YTH4940) strains transformed with an EV control or 3 different peptide constructs were performed. Cells were grown in 2% glucose (left) or 2% galactose (right). The top panels were grown at 30°C, while the lower panels were grown at 37°C. Cells were grown for 3-5 days, then imaged. **(D)** Normalization of 100-fold dilution (third column) for each strain was performed and plotted. n=1. **(E)** Spot dilutions of W303 WT (YTH1) cells transformed with an EV, and *cdc20-1* (YTH3863) cells transformed with an EV or the 4 peptide constructs shown are performed. Cells were grown in 2% glucose (left) or 2% galactose (right) to induce peptide expression. Plates were incubated for 3-5 days, then imaged. **(F)** Normalization of 100-fold dilution (third column) for each strain was performed and plotted; n=1.

If Cdh1 was required for the peptide phenotypes, then we should no longer observe slowed growth when induced with 2% galactose in *cdh1Δ* cells. In cells expressing C13-3, growth remained significantly suppressed, indicating that Cdh1 does not affect C13-3 function. However, in *cdh1Δ* cells expressing C43-4, cell growth was largely restored. Opposed to C13-3, C43-4 appears to require Cdh1 for function. No substantial growth phenotype was observed for C2-4B, and thus we could not determine if it required Cdh1 for function. Next, we expressed the peptides shown in temperature-sensitive *cdc20-1* cells (**Figure 4E**; quantified in **Figure 4F**). We observed that with all the peptides tested, Cdc20 is required for activity, as all transformants grew to the same extent as the control cells expressing an empty vector when induced with galactose. It should be noted that growth on galactose suppressed the slow growth of *cdc20-1* cells observed on glucose. It has been described previously that glucose inhibits APC activity, while alternative carbon sources, such as glycerol, galactose, and raffinose, can suppress APC mutations (Baumer et al. 2000; Harkness et al. 2004). Taken together, these results support the idea that C43-4 enhanced both APC^Cdc20^ and APC^Cdh1^ activity. C13-3 appears to activate only APC^Cdc20^, with the remaining peptides requiring testing in future studies.

### APC-binding peptides increase the replicative lifespan of WT cells

We have previously shown that the APC is required for longevity in yeast (Harkness et al. 2004; Postnikoff et al. 2012; Menzel et al. 2014; Malo et al. 2016). Therefore, we predicted that peptide activation of the APC would increase yeast RLS, as M2I-1 did (**Figures 2C**, **2D**). We transformed the peptides shown into WT cells (YTH2066) for RLS (**Figure 5A**; stats, mothers scored, and mean RLS shown in **Supplemental Table 3**). The cells were maintained on SD -Trp media supplemented with 2% glucose for low-level expression, as we have shown previously that 2% glucose results in low, but detectable levels of expression from galactose-inducible promoters (Turner et al. 2010; Postnikoff et al. 2012; Menzel et al. 2014). Under these conditions, the mean RLS of WT cells expressing an empty vector (EV) or most of the peptides was shorter than normally observed on YPD (**Supplemental Table 3**). However, peptides C2-4B and C13-3 increased the mean RLS by ∼50% of that for control cells. We then tested the RLS of the cells maintained on YPD (**Figure 5B**). The mean RLS of control cells was 20.87 +/- 1.45 generations (**Supplemental Table 4**), as expected for YPD. Additionally, C2-4B continued to increase RLS, whereas C13-3 did not. The increase in RLS using C2-4B, however, did not reach significance (**Supplemental Table 4**). Utilizing the C43-4 peptide, we measured RLS when the peptide construct-expressing cells and controls were induced with YP 0.5% galactose/1.5% sucrose, and compared to cells maintained on YP 2% glucose (YPD). We used cells expressing the HA-TrxA-C43-4 cassette that was integrated into the genome of Clb1-TAP cells (as used in **Figures 3F, 3G**) to avoid the potential inconsistent effects of transforming plasmids. We observed that cells expressing integrated C43-4 induced with 0.5% galactose lived the longest (**Figure 5C**; **Supplemental Table 5**). The mean RLS of WT + YPD was 22 +/- 1.74 generations, while the mean RLS of WT + C43-4 + 0.5% galactose was 29.66+/- 1.97 generations, with a *p*–value of 0.0057. This indicates that C43-4 requires galactose induction to increase yeast RLS.

**Figure 5.**
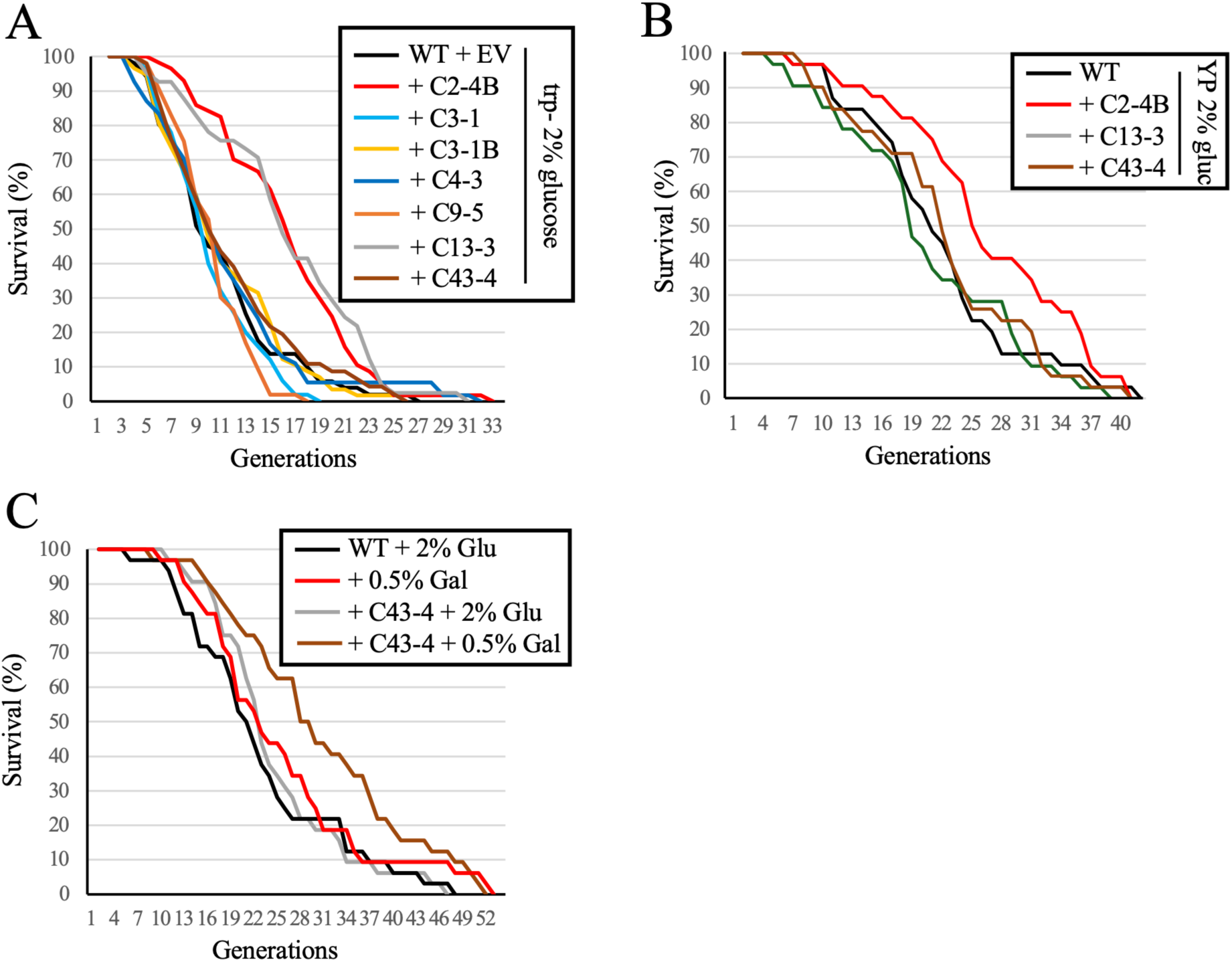
The peptides have different effects on RLS in cells. **(A)** WT cells (YTH2066) were transformed with the peptides shown for RLS assays. Mother cells were placed on SD -Trp plates for the entirety of the experiment. Daughter cells were picked from each mother cell, counted and discarded. Counting continued until the last mother stopped dividing. Statistical data was generated using OASIS2 and is shown in **Supplemental Table 3**. *p*–values for WT control vs WT + C2-4B was 9.4 × 10^−6^**\*\*\*\*** and for WT control vs WT + C13-3 is 2.5 × 10^−5^****. **(B)** Peptides C2-4B, C13-3 and C43-4 were selected for further study. The Y2H strain (YTH3640), co-transformed with *APC10-LEXA* and the peptides shown, were used for RLS. Mother cells were placed on YPD for the entirety of the RLS experiment. Daughters were picked from each mother until all mothers stopped dividing. Statistical data is shown in **Supplemental Table 4**. **(C)** Clb1-TAP cells and Clb1-TAP cells expressing integrated HA-TrxA-C43-4 were used for RLS in the presence of YPD or YP 0.5% galactose/1.5% sucrose. Statistical data is shown in **Supplemental Table 5**. The *p*-value for WT 2% glucose vs WT + C43-4 + 0.5% galactose is 0.0057**. All other comparisons were not significant.

### The C43-4 peptide increases CLS

RLS measures how many cell divisions a mother cell can go through prior to death, which is controlled by APC^Cdc20^ during mitosis and APC^Cdh1^ during mitotic exit and G1. To determine if the peptides can activate a function reliant on APC^Cdh1^ alone, we used CLS. CLS measures how long a population of quiescent cells can remain metabolically active. Entrance and maintenance of G0 is dependent upon APC^Cdh1^ activity (Skaar and Pagano 2008; Cappell et al. 2016). It is important to identify APC^Cdh1^ activators, as this version of the APC is associated with tumor suppression and maintenance of genome stability (García-Higuera et al. 2008; Wäsch et al. 2010; Greil et al. 2016; Ha et al. 2017). To the best of our knowledge, there are no known APC^Cdh1^ activators. As shown above, the commercial APC activator M2I-1 increased RLS, but did not increase CLS (**Figures 2C-2E**). This was not surprising since M2I-1 works specifically to release Cdc20 from the SAC, and is therefore not expected to impact an APC^Cdh1^-dependent activity, such as CLS. We tested cells expressing plasmid-borne peptides grown to stationary phase in SD -Trp media supplemented with either 2% sucrose or 1% galactose/1% sucrose for CLS. Peptides C2-4B, C9-5, and C13-3 did not increase CLS when induced, while C43-4 did (**Figures 6A-D**; AUC calculations shown to the right of each panel; mean and max lifespans and 95% confidence intervals shown in **Supplemental Table 6**; all samples were done at the same time with the same WT control). The HA-TrxA-C43-4 cassette was cloned into a 2μ *URA3* plasmid, which also increased CLS in WT cells (**Figure 6E**). We next assessed the CLS of Clb1-TAP cells harboring integrated copies of HA-TrxA-C13-3 and HA-TrxA-C43-3. Only the C43-4 expressing cells, when induced with 1% galactose/1% sucrose exhibited increased CLS (**Supplemental Figure 8**). We then assessed CLS in response to increased galactose concentrations. Using Clb1-TAP cells +/- the integrated HA-TrxA-C43 cassette, we found that the CLS was unaltered by carbon source usage in cells lacking C43-4 (**Figure 6F**; AUC quantified on the right). However, in the presence of C43-4, the CLS increased proportionally to the galactose concentration (**Figure 6G**; AUC quantified on the right). Lastly, W303-based WT (YTH1), *apc5^CA^*(YTH5065), and *apc10Δ* (YTH4667) cells were transformed with the HA-TrxA-C43-4 plasmid, followed by CLS using MM SD -Trp media supplemented with either 2% sucrose or 1% galactose/1% sucrose (**Figure 6H**; quantitation and mean, max, and stats shown in **Supplemental Figure 9**). We observed that while C43-4 induction increased the CLS of WT cells (note that CLS is longer in SD media compared to MM; compare the CLS in MM in **Figure 6A** with CLS in SD media in **Figure 6H**), it did not increase the CLS of the APC mutants tested. The results shown here indicate that C43-4 is the only peptide, expressed from the TrxA cassette, to increase CLS in an APC-dependent manner.

**Figure 6.**
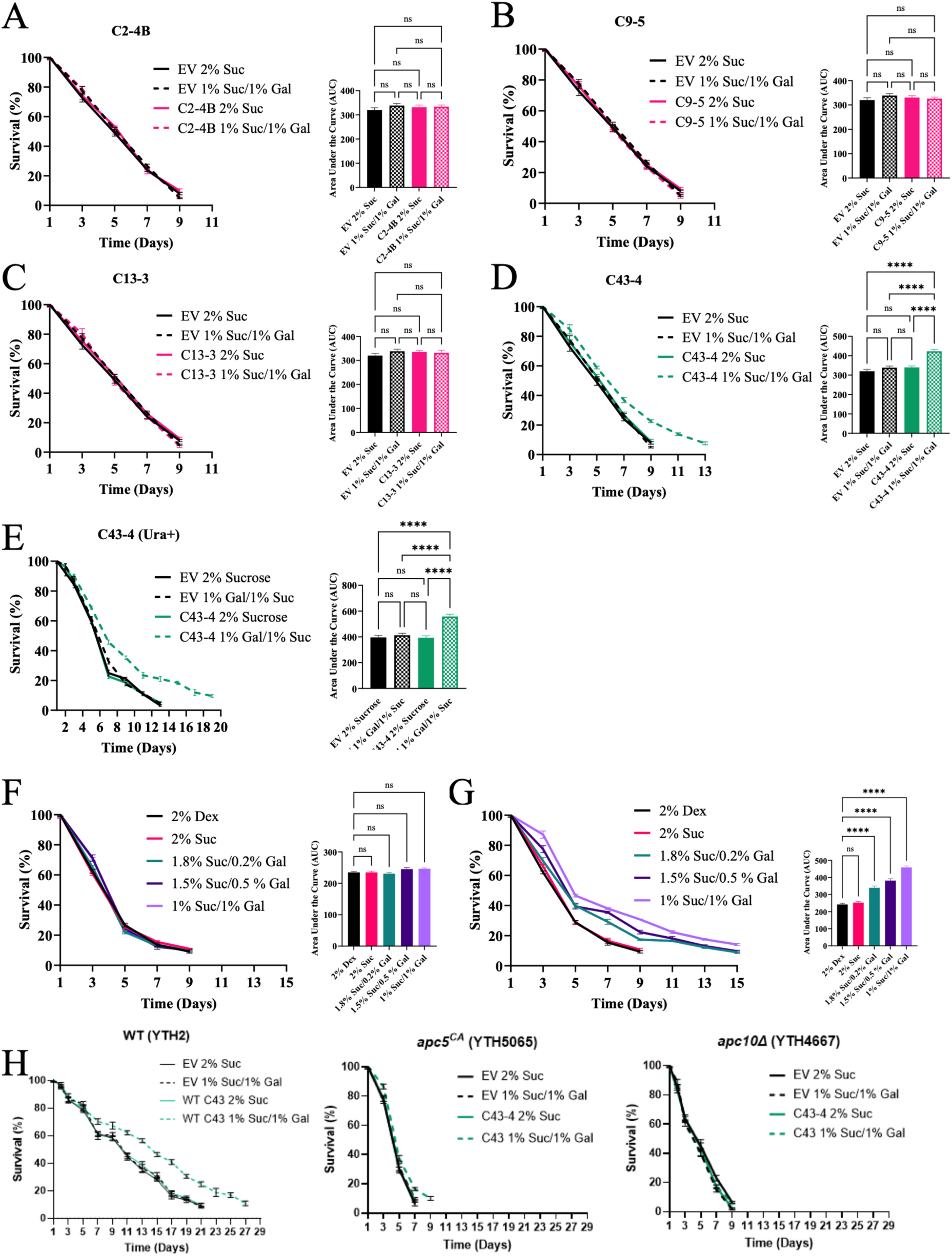
Only C43-4 increases CLS. CLS assays using W303-based cells transformed with the +Trp pJG4-5 EV control and four peptide constructs: **(A)** C2-4B, **(B)** C9-5, **(C)** C13-3, and **(D)** C43-4, grown in MM -Trp media with 2% sucrose or 1% galactose/sucrose. Values are presented as mean ± SEM for n=2. **(E)** CLS assays using WT S288c-based cells (YTH4504) transformed with the Ura+ pRS416 EV or the pRS416*-*C43-4 peptide construct in depleted MM -Ura media. Survival in the experiments above was assayed by methylene blue cell counts every second day starting on Day 1 until survival reached <10%. Values are presented as mean ± SEM for n=3. Area under the curve (AUC) calculations of each strain ± SEM were analyzed with a one-way ANOVA and Tukey’s multiple comparisons *post-hoc* test where *p*<0.0001**** between strains containing the C43-4 peptide in galactose and the control strains. Graphs are shown in the panels to the right. Mean and max lifespans and 95% confidence intervals are shown in **Supplemental Table 6**. **(F)** CLS experiments with Clb1-TAP (YTH4990) or **(G)** Clb1-TAP cells with integrated HA-TrxA-C43-4 (YTH5123) were used for CLS experiments. CLS on sucrose was compared to CLS on glucose (Dex). AUC quantitation is shown to the right. **(H)** CLS was performed on W303-based WT (YTH1), *apc5^CA^* (YTH5065), and *apc10Δ* (YTH4667) cells using the methylene blue method, when maintained in depleted SD -Trp media. Quantitation and stats are shown in **Supplemental Figure 6**.

### The APC-binding peptide C43-4 increases APC activity in chronologically aging cells

At the beginning of this study, we showed that the APC loses activity as cells age (**Figure 1**). To test whether APC activity can be enhanced in quiescently aging cells, we used plasmid-borne HA-TrxA-C43-4 and the EV as a control, transformed into Clb1-TAP BY4741 cells, as done above (**Figure 1**). Cells were removed on the days shown and allowed to re-enter the cell cycle in the presence of nocodazole, then arrested in G1 using α-factor, (budding index shown in **Supplemental Figure 10**). We expected that aging cells re-entering the cell cycle would have to rely on the activity of the APC currently in the aging cells. If the APC is impaired, then targeted degradation of Clb1-TAP at G1 would be deficient, leading to Clb1-TAP levels rising in G1 as cells aged (**Figure 7A**, first panel from the left; increased G1:M ratio, rightmost panel). We observed this accumulation in cells expressing the EV in glucose (**Figures 7A**), similar to **Figure 1D**. Cells expressing the EV maintained in 2% galactose showed the same effect (**Figure 7B**), with the G1:M ratio increasing as cells aged (**Figure 7B**, rightmost panel). When cells expressing C43-4 were maintained in glucose while aging, we again observed a rise in the G1:M ratio (**Figure 7C**; G1:M ratio shown in rightmost panel), indicating that the APC remains impaired while aging when C43-4 is not induced. However, when we maintained C43-4 expressing cells under inducing conditions, we observed that Clb1-TAP G1 levels were low throughout the aging experiment (**Figures 7D**; G1:M ratio shown in rightmost panel). C43-4 abundance increased in aging cells under inducing conditions, indicating that C43-4 remains inducible throughout the aging experiment. The low G1:M ratio indicates that in the presence of induced C43-4, the APC remains active.

**Figure 7.**
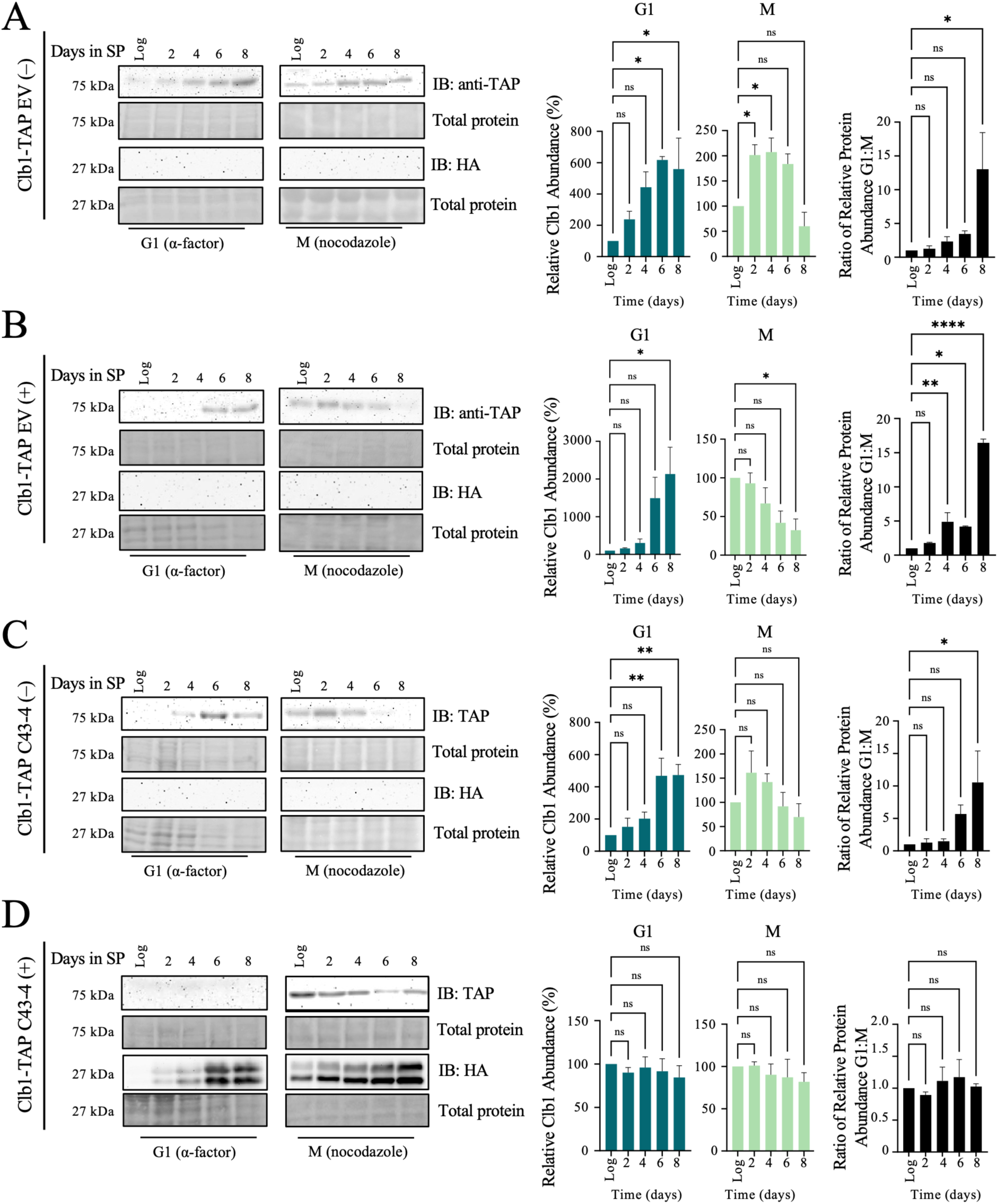
Degradation of APC substrate Clb1 is increased in aging cells upon APC activation by the C43-4 peptide construct. Western blot analyses performed on lysates harvested from Clb1-TAP strains containing an EV or the integrated HA-TrxA-C43-4 construct in log and stationary phase over 8 days, maintained in depleted MM -Trp media with 2% glucose (-; **A** and **C),** or 2% galactose (+; **B** and **D),** to induce C43-4 expression. On the days indicated, samples were resuspended in nutrient-rich media to re-enter the cell cycle in the presence of nocodazole or α-factor to arrest cells in mitosis or G1, respectively. Full uncropped blots are shown in **Supplemental Figure 11**. Densitometry was used to normalize proteins to the total protein Ponceau S stains to determine relative abundance in the indicated stage of the cell cycle in aging cells between M and G1. Normalized graphs were used to compare relative G1:M ratios over time (black bar graphs, right). Values are presented as mean ± SEM for n=3 and were analyzed with a one-way ANOVA and Dunnett’s multiple comparisons *post-hoc* test where *p*<0.05*, *p*<0.01** and *p*<0.001*** between log phase and each day of stationary phase.

To get a view of whether the C43-4 peptide can be used therapeutically to activate APC^Cdh1^, we synthesized the 20 amino acid C43-4 fused to an FITC motif (C43-4^FITC^) and a scrambled control, also fused to FITC (scrambled^FITC^). We tested whether C43-4^FITC^ required a carrier peptide to cross the cell wall and plasma membrane. We incubated C43-4^FITC^ and scrambled^FITC^ with the cell permeable TAT2 peptide, which consists of 2 repeats of the membrane-penetrating sequence RKKRRQRRR. The TAT2 cell penetrating peptide is derived from the HIV-1 transactivator protein (Torchilin 2008), to allow complex formation. The TAT2 peptide has been observed to enter yeast cells, but the efficiency is far less compared to mammalian cells due to the yeast cell wall (Isogai et al. 2025). We performed CLS in BY4741 cells with 5 μg/ml C43-4^FITC^ or scrambled^FITC^ added to the growth media at the beginning of culturing, in the presence or absence of 5 μg/mL complexed TAT2. The results show that C43-4^FITC^ increased CLS in the presence or absence of TAT2 (**Figure 8A**). Neither the addition of TAT2 alone nor the addition of scrambled peptide influenced CLS. This confirmed that the C43-4 peptide sequence, and not the TrxA backbone, is necessary for extending lifespan. Next, we added increasing amounts of C43-4^FITC^ or scrambled^FITC^ in the absence of TAT2 to the cell media. We added 1, 5 or 10 μg/mL peptide at the beginning of culturing, or added 5 μg/mL at the beginning of culture and again at Day 1 of stationary phase. We observed that increasing the amount of scrambled^FITC^ had no effect, but increasing C43-4^FITC^ steadily increased the maximum CLS of cells (**Figure 8B**). It is noteworthy that a single C43-4^FITC^ dose at the beginning of the experiment is sufficient to increase CLS. Also, the addition of 5 μg/mL at the beginning and again upon entering stationary phase was equivalent to adding one dose of 10 μg/mL at the beginning of the experiment, indicating that C43-4 aids both dividing and non-dividing cells. We were next interested in determining if C43-4^FITC^ could increase the CLS of already aging cells. For this experiment, 5 μg/mL C43-4^FITC^ was added either at the beginning of culturing, or at Days 1, 3, 5 or 7 of stationary phase (**Figure 8C**). The greatest effect on lifespan was observed when adding C43-4^FITC^ to the mitotically growing cells prior to entry into stationary phase. As can be seen in **Figure 8C**, the CLS also increased when C43-4^FITC^ was added to aging cells, even at Day 7. This suggested that even later in stationary phase, lifespan could still be extended, and that APC impairment with age can be reversed. To determine whether C43-4^FITC^ required either of the APC co-activators, Cdc20 or Cdh1, we added C43-4^FITC^ to W303-based WT, *cdc20-1* or *cdh1Δ* cells at the beginning of culture (**Figure 8D**). We observed that both *cdc20-1* and *cdh1Δ* cells had a short CLS, as observed in **Figure 2A**. C43-4^FITC^ increased the CLS of W303 WT cells, but could not increase CLS in the mutant backgrounds, indicating that C43-4 requires Cdc20 and Cdh1 to function as a CLS extender.

**Figure 8.**
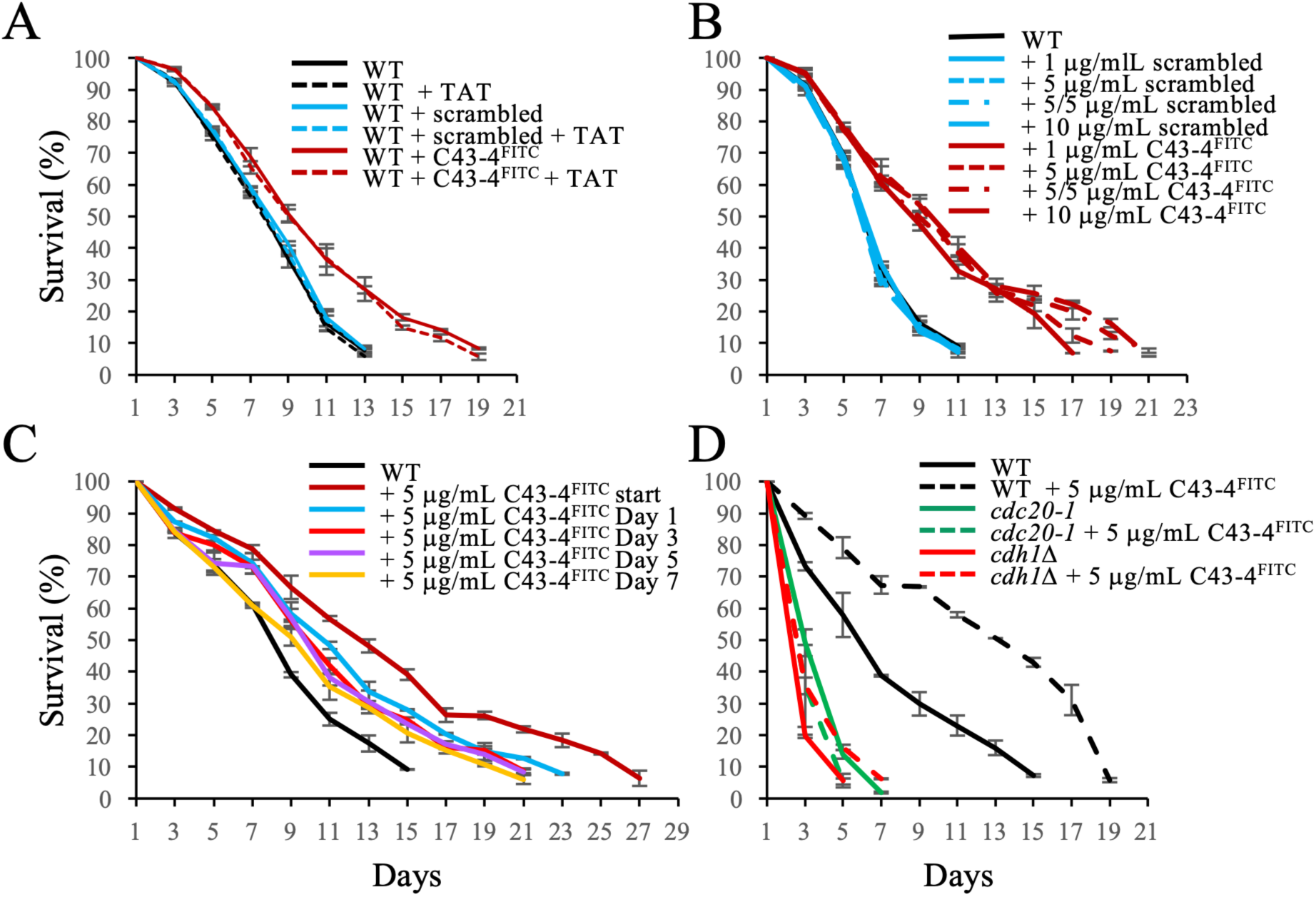
A synthesized 20 amino acid C43-4 fused with FITC (C43-4^FITC^) increases lifespan. **(A)** WT BY4741 cells (YTH4504) were grown in the presence or absence of the 5 μg/mL synthetic C43-4^FITC^. Scrambled peptide fused to FITC and the TAT2 permeable peptides were used as controls. Cells were grown to stationary phase, with samples removed every other day to assess viability using methylene blue to identify dead cells. **(B)** BY4741 cells were grown to stationary phase in the presence of 1, 5, or 10 μg/mL C43-4^FITC^ added to the culture at the beginning of growth. In another sample, 5 μg/mL C43-4^FITC^ was added at the beginning, and again when cells stopped dividing. Scrambled peptide was used in the same manner. Cell survival was determined using methylene blue. **(C)** BY4741 cells were grown to stationary phase. C43-4^FITC^ was added at the beginning of growth, then at the days shown in different samples while the cells were chronologically aging. Cell survival was assessed, as above, with methylene blue. **(D)** W303-based WT, *cdc20-1*, and *cdh1Δ* cells were grown to stationary phase and aged chronologically, in the presence or absence of C43-4^FITC^. Survival was assessed using the methylene blue method. Error bars represent mean ± SEM; n=3.

To confirm that C43-4^FITC^ was reaching the nucleus, we used an antibody that recognized the FITC motif. First, we observed that C43-4^FITC^ was present in whole cell lysates following overnight incubation and multiple washes, but the scramble peptide was not (**Supplemental Figure 12A**). To ensure the scrambled peptide could be recognized by the FITC antibody, we spiked lysates with 1 μg C43-4^FITC^ or scrambled^FITC^. We observed that both FITC labelled peptides could be recognized by the antibody (**Supplemental Figure 12B**). We then complexed the scrambled peptide with TAT2 and added this to cells overnight. We still did not observe scrambled^FITC^ in cells (**Supplemental Figure 12C**). As observed with CLS, where the TAT peptide was not required to extend C43-4-dependent CLS (**Figure 8A**), the TAT carrier peptide was not required for translocation of C43-4 across the plasma membrane (**Supplemental Figure 12C**). Using lysates fractionated into cytosolic and nuclear fractions, we observed that C43-4^FITC^, like HA-TrxA-C43-4, did indeed enter the cell and the nucleus (**Figure 9A**, uncropped blots shown in **Supplemental Figure 12D**). However, since the scrambled peptide did not enter the cell, we required a control peptide that entered the cell and nucleus, but did not increase CLS.

**Figure 9.**
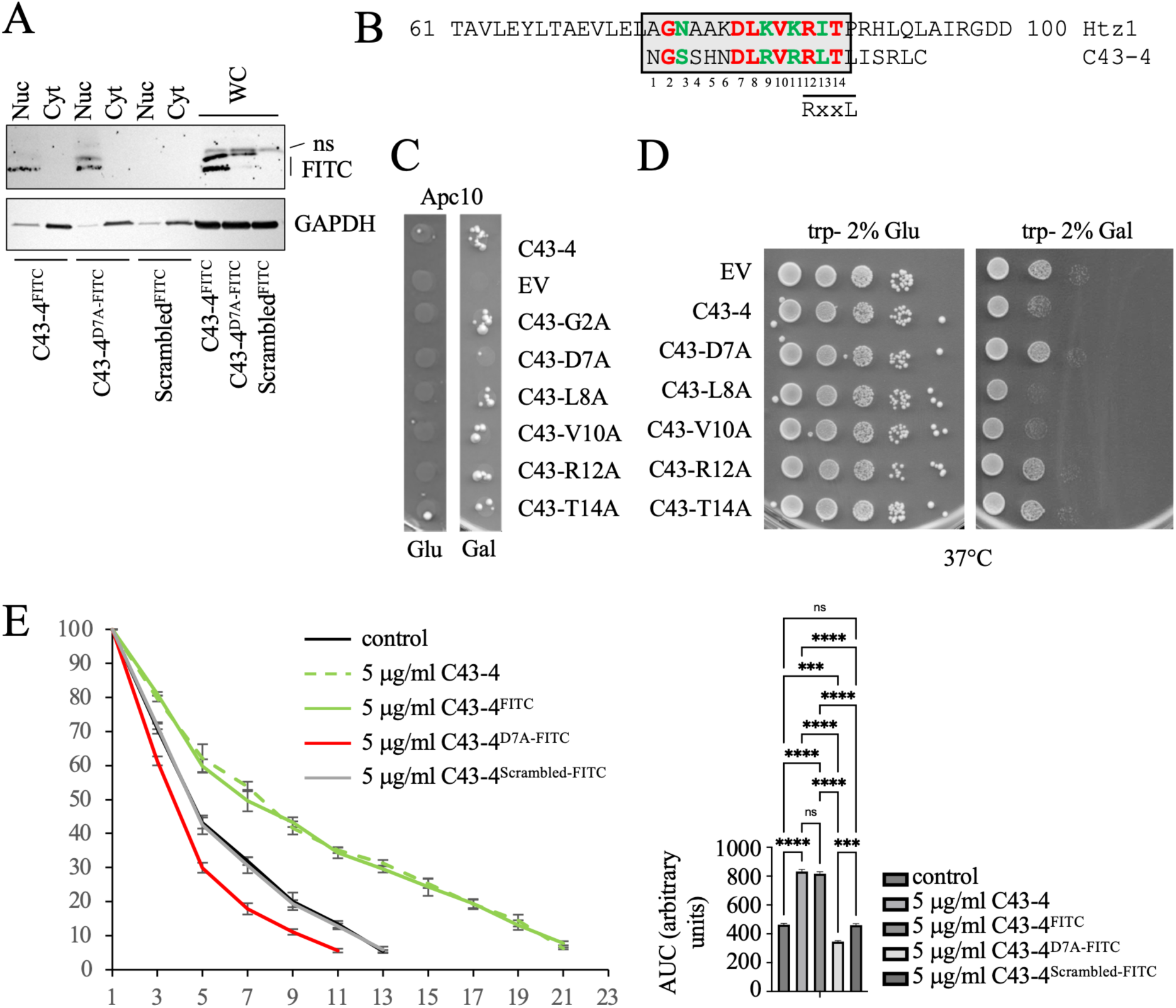
Identification of amino acids required for C43-4 function. **(A)** Cells incubated with the C43-4^FITC^ or untreated control cells were fractionated into cytosol and nuclear fractions. Antibodies against FITC, histone H3, and GAPDH were used. C43-4, a mutated C43-4^D7A^, and the scrambled peptide were incubated with cells overnight. Full uncropped blots are shown in **Supplemental Figure 12D**. **(B)** Homology of C43-4 with a region of Htz1, with shared amino acids shaded in red and amino acids of the same class shaded in green. The RxxL mofit is noted. The amino acids shaded in red in C43-4 were mutated to alanine. **(C)** A 2-hybrid test using the mutated C43-4 constructs and LexA-Apc10. Cells were spotted onto -Trp/His/Ade supplemented with either 2% glucose or galactose. **(D)** The C43-4 mutants were expressed in W303-based cells and grown at 37°C on media containing 2% glucose or 2% galactose. The plates were grown at 37°C for 2-3 days. **(E)** The peptides shown were added to BY4741 cells at the beginning of the CLS experiment. The cells were maintained in depleted CM for the duration of the experiment. Counts using methylene blue staining were stopped once the cells alive were below 10%. Three biological repeats were performed for each sample with mean ± SEM shown. Area under the curve (AUC) was calculated for each curve and plotted.

To generate a control for the synthetic C43-4 peptide, we mutated C43-4. In order to determine which amino acids to mutate, we performed a BLAST search of the peptide amino sequence against the yeast proteome. We observed that C43-4 shared sequence with a number of yeast proteins (**Supplemental Figure 13A**). C43-4 encoded the D-box RxxL motif, which is observed in many APC substrates (Arnold et al. 2015). Proteins encoding the RxxL motif are often recruited to the APC by interacting with the Apc10-Cdc20/Cdh1 co-receptor (Carroll et al. 2005; da Fonseca et al. 2011). C43-4 shared sequence homology with the proteins Cst9 and Rpt1, both of which express an RxxL, but are not known to serve as APC targets. We chose Htz1, a stress-responsive histone H2A variant (Srivatsan et al. 2018; Saatchi et al. 2019; Brewis et al. 2025) as our target, as we previously demonstrated that the APC interacts genetically with histones during mitosis (Harkness et al. 2002; Harkness et al. 2005; Turner et al. 2010; Islam et al. 2011). Expression of galactose-inducible *GST-HTZ1* (kind gift from A. Emili, U of Toronto) in WT, *apc5^CA^*, and *apcΔ10* cells, even at low levels in glucose, showed that *HTZ1* could suppress APC mutant defects (**Supplemental Figure 13B**). Since Htz1 is described as stress-responsive, we asked if GST-Htz1 protein levels are also responsive to APC activity. *GST-HTZ1* expressed in the mutants shown above were grown in 0.5% galactose for 16 hours to synthesize GST-Htz1. The cells were then arrested in mitosis with nocodazole, followed by treatment with cycloheximide (CHX) to inhibit the ribosome. Samples were taken every 30 minutes for 90 minutes (**Supplemental Figure 13C**). Antibodies against Clb2, an APC substrate, were used as a control. We observed that Clb2 levels were elevated and stable in *apc5^CA^*and *apc10Δ* cells, as expected. Htz1 levels were also elevated in *apc5^CA^*cells, and more so in *apc10Δ* cells. Htz1 was partially stabilized in the APC mutants, indicating that the APC, as well as another E3, likely control Htz1 protein levels. Lastly, we asked if Htz1 levels may impact the protein levels of APC subunits. We expressed *GST-HTZ1* in Apc5-TAP and Apc10-TAP cells. As a control, we expressed a vector expressing only GST. Cells were grown overnight in glucose, then switched to 0.5% galactose-supplemented media, with samples recovered at 0, 8 and 16 hours for western analyses. We observed that Apc5-TAP levels were unchanged throughout the experiment, whereas Apc10-TAP was weakly expressed at the beginning of galactose induction, and decreased after 16 hours in control GST cells (**Supplemental Figure 13D**). However, in the presence of GST-Htz1, Apc10-TAP levels increased over the time course. This suggests that Apc10 is an unstable protein that is stabilized by Htz1. This provides a potential mechanism whereby C43-4 may mimic Htz1 activity by interacting with the APC, possibly stabilizing Apc10, and promoting stress-responsive APC activity.

We therefore mutated all amino acids in C43-4 that were shared with Htz1 individually to alanine (amino acids shaded in red in **Figure 9B**). We then expressed the mutant versions of C43-4 in yeast 2-hybrid cells co-expressing LexA-Apc10 to determine if any of the mutations disrupt the C43-3-Apc10 2-hybrid interaction. We observed that all mutant peptides, except C43-4^D7A^ still interacted with Apc10 (**Figure 9C**). Next, we spot diluted W303-based cells expressing the C43-4 mutants on SD -Trp- media at 37°C supplemented with either 2% glucose or 2% galactose. We observed that the WT C43-4 peptide slowed the growth of cells, as did the L8A and V10A variants (**Figure 9D**), indicating that L8 and V10 are not required for C43-4 function. The D7A, R12A and T14A mutants all grew like the empty vector control, indicating that these amino acids are required for function. Notably, D7 is required for the 2-hybrid interaction, while R12 and T14 make up the RxxL domain. We therefore synthesized the C43-4^D7A^ peptide fused to FITC. When C43-4^D7A^-^FITC^ was added to cells overnight, we observed that the peptide was localized to the nucleus following cell fractionation (**Figure 9A**). We then added the peptide to cells at the beginning of log growth for CLS studies. We observed that C43-4^D7A^-^FITC^ did not increase CLS, but slightly reduced it (**Figure 9E**). Therefore, the D7A mutation disrupts Apc10-peptide interactions, C43-4 function, and abolishes extended CLS, supporting our observations that C43-4 interacts with the APC and prolongs CLS.

Next, to gather additional evidence that a functional APC is required for C43-4^FITC^ to increase CLS, we added 5 μg/mL C43-4^FITC^ to cultures of BY4741 cells harboring APC mutations (**Figures 10A-H**; AUC and statistical calculations shown in **Figure 10I**). C43-4^FITC^ could not increase the CLS of any of the APC mutants used, and could not increase the CLS of APC mutants in W303- or BY4741-based cells, indicating that the effect is unlikely due to the genetic background. We also used a strain lacking Ama1, a meiosis-specific APC activator that is involved in spore formation (Cooper et al. 2000). We expected that *ama1Δ* cells would have a short CLS, as stationary phase often leads to meiosis under the right conditions (Reinders et al. 1998). Indeed, the deletion of *AMA1* shortened CLS, which C43-4^FITC^ could not restore. This indicates that C43-4^FITC^ requires a functional APC to increase CLS in a strain background independent manner.

**Figure 10.**
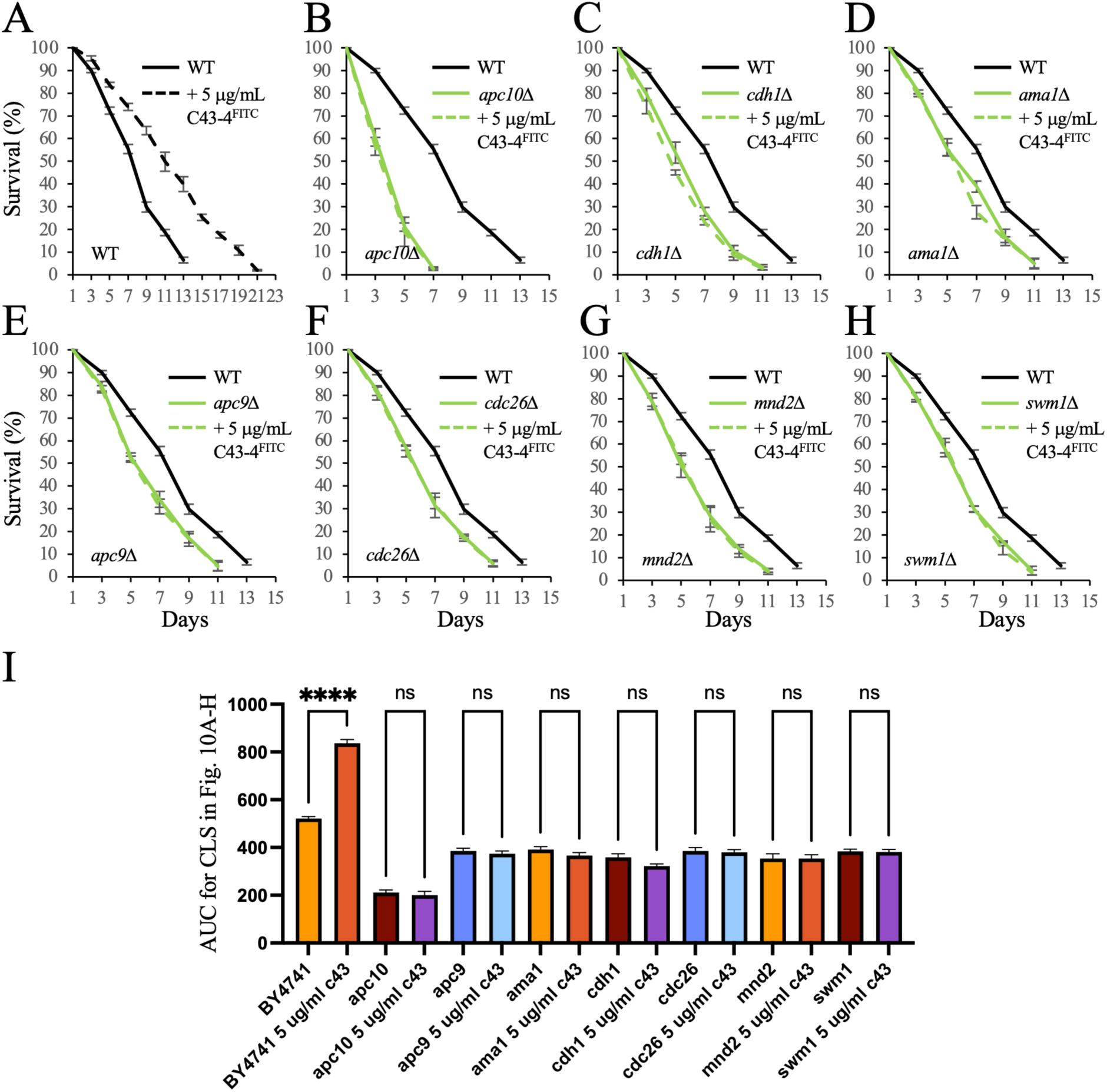
A functional APC is required for increased C43-4-dependent CLS. **(A)** WT BY4741-based cells (YTH4505) were grown to stationary phase and allowed to chronologically age, in the presence or absence of 5 μg/ml C43-4^FITC^. Cell survival was determined and plotted using the methylene blue technique. **(B)** – **(H)** *apc10Δ* (YTH5020), *cdh1Δ* (YTH5126), *ama1Δ* (YTH5131), *apc9Δ* (YTH5127), *cdc26Δ* (YTH5128), *mnd2Δ* (YTH5129), and *swm1Δ* (YTH5130) cells, all from the BY4741 deletion collection, were grown to stationary phase in the presence or absence of 5 μg/ml C43-4^FITC^. Cell survival was measured using the methylene blue method and compared to WT curves. The experiment was done together with the same WT with biological triplicates. In all cases, the mean ± SEM is shown. **(I)** AUC was calculated for the CLS experiments shown above.

### C43-4 increases lifespan in *C. elegans* in a *daf-16* and *aak-2*-dependent manner

Lastly, we asked if the lifespan function of C43-4 is conserved in higher eukaryotes. We chose to test this in the worm *Caenorhabditis elegans.* The construct shown in **Figure 11A** was cloned to express C43-4 fused to GFP under the ubiquitous *eft-3* gene promoter. Stable integration of the C43-4 transgene showed consistent expression as visualized by GFP fluorescence (**Figure 11B**). Post-mitotic lifespan was then carried out on these worms. We observed that C43-4 continued to increase lifespan in worms (**Figure 11C**; stats calculations shown in **Supplemental Table 7;** *p* < 0.001). Our work in yeast demonstrated that the APC works in a feedforward loop with the stress response factors Fkh1/Fkh2 and Snf1 (Harkness et al. 2004; Postnikoff et al. 2012; Jiao et al. 2015; Malo et al. 2016), which are members of the evolutionarily conserved FOXO3A and AMPK families, respectively. We introduced the loss-of-function mutant alleles of *daf-16* and *aak-2* into the integrated C43-4-GFP construct and performed lifespan assays. Increased lifespan induced by C43-4 partially required *daf-16* (**Figure 11D**; stats calculations shown in **Supplemental Table 8;** *p* < 0.00001 for all combinations). Increased C43-4 lifespan also required *aak-2* (**Figure 11E**; stats calculations shown in **Supplemental Table 9;** *p*<0.0001 on average for all combinations, except N2 vs *aak-2* C43-4 [0.3218, 0.0033, and 1.1 × 10^−7^ in the three trials, and *aak-2* vs *aak-2* C43-4, which was ns in 2 out of 3 trials]). This indicates further evolutionary conservation, as the yeast APC requires the Fkh proteins or Snf1 to sense stress and increase lifespan (Harkness et al. 2004; Postnikoff et al. 2012; Jiao et al. 2015). As a final validation, we expressed C43-4 in worms with a loss-of-function mutation to *akt-1*. These worms naturally have a longer lifespan, similar to what is observed in yeast, as *sch9Δ* mutants live long (Fabrizio et al. 2001). In *akt-1* mutant worms expressing C43-4, we observed that *akt-1* loss-of-function was epistatic to C43-4, as they continued to experience *akt-1*-dependent long life and did not show a further extension (**Figure 11F**; stats calculations shown in **Supplemental Table 10;** *p*<0.0001 for all comparisons, except *akt-1* vs *akt-1* C43-4, which was ns in all trials). This indicates that induction of longer life by C43-4 may involve inhibition of the Akt-1 protein. Taken together, our data indicates that we have discovered the first APC^Cdh1^ activator that increases the lifespan of quiescent cells in yeast and worms.

**Figure 11.**
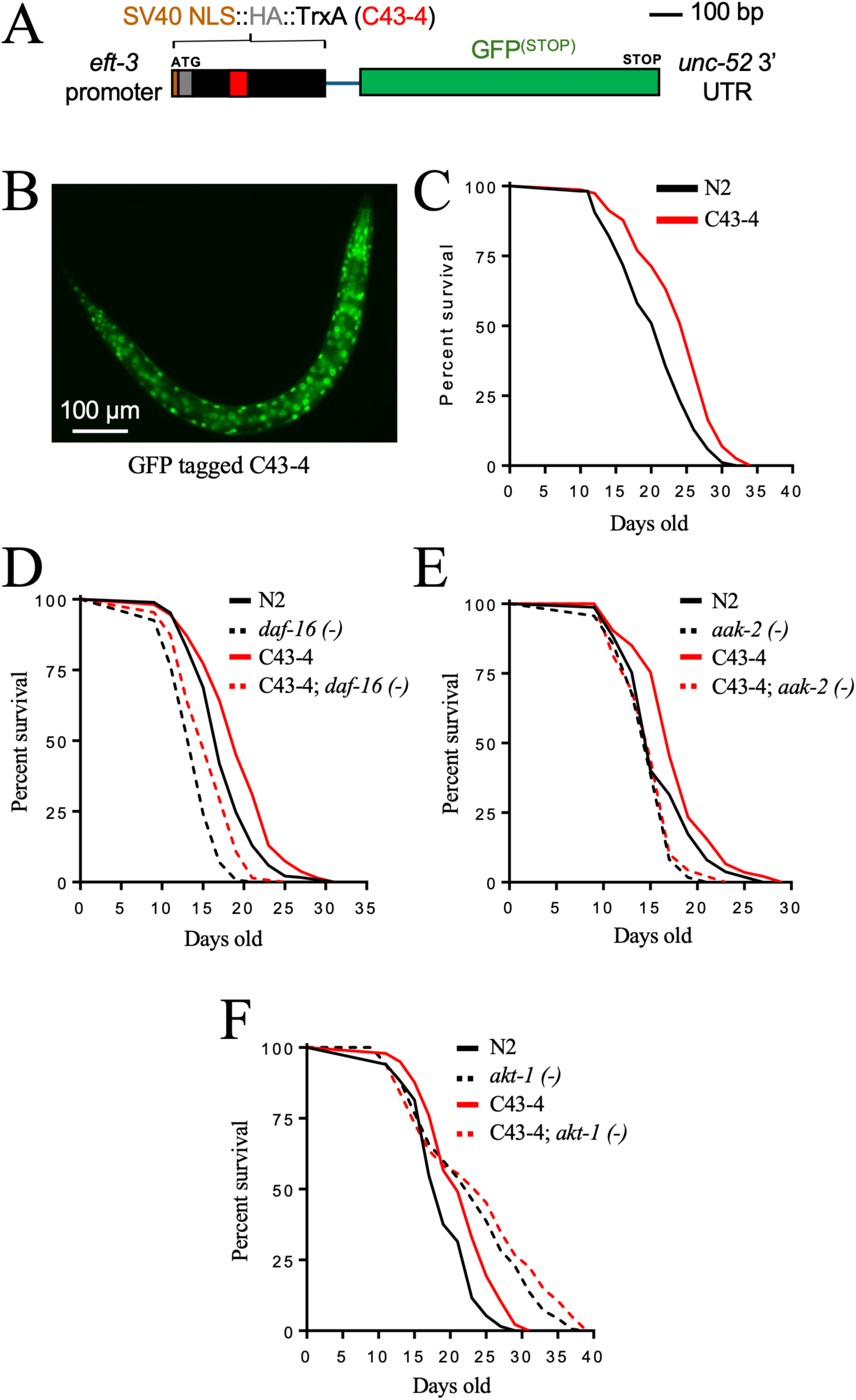
Effects of C43.4 expression in *C. elegans*. **(A)** Schematic of the C43-4 plasmid built for *C. elegans* expression. **(B)** C43-4-GFP expression pattern in an L4 *C. elegans*. **(C)** Expression of C43-4 extends *C. elegans* lifespan compared to the N2 wildtype. N=98-124 worms scored for each condition; trial 3 is shown. **(D)** Effects of C43-4 on lifespan in the *daf-16(mu86)* loss-of-function background, n=129-186 worms measured per condition; trial 1 is shown. The *daf-16* gene is the yeast *FKH1* orthologue. **(E)** Effects of C43-4 on lifespan in the *aak-2(ok524)* loss-of-function background, n=146-175 worms measured per condition; trial 3 is shown. The *aak-2* gene is the yeast *SNF1* orthologue. **(F)** Effects of C43-4 on lifespan in the *akt-1(ok525)* loss-of-function background, n=133-184 worms measured per condition; trial 1 is shown. The *akt-1* gene is the yeast *SCH9* orthologue. For all lifespan assays, 3 independent trials were performed with full results and statistics for all trials shown in **Supplemental Tables 7**-**10**.

## DISCUSSION

In the study presented here, we sought to discover small peptides that directly activate APC^Cdh1^. Mammalian APC^CDH1^ activation can occur via CDK4/6 inhibitors that block phosphorylation of CDH1, allowing APC^CDH1^ to remain active and prevent S phase transit (Mouery et al. 2024). However, these inhibitors are indirect and impact many other processes. Chemicals that indirectly activate APC^CDC20^ through SAC inhibition are available, but the one tested in this study, M2I-1, does not increase the lifespan of non-dividing quiescent cells. Thus, even though SAC inhibitors are moving towards clinical trials, with unsatisfactory results (Zhou et al. 2016; Novais et al. 2012; Pinto et al. 2023), we predict that these chemicals will not increase the cell health of quiescent differentiated cells. A key example of this is the effect CDH1 has on the proper function of neurons (Almeida et al. 2005; Bobo-Jimenez et al. 2017), where CDH1 is involved in neuronal differentiation and the developing brain. A loss of APC7 was recently described to be at the heart of an intellectual disability syndrome in humans, suggesting that the APC subunit, APC7 is required for the developing mammalian brain (Ferguson et al. 2022). In these experiments, the APC was shown to target the proliferation marker, Ki-67, for ubiquitin-dependent degradation when associated with heterochromatin. Conditional knockout of CDH1 in mice, but not CDC20, led to the accumulation of Ki-67, linking APC^CDH1^, but not APC^CDC20^, with brain development. In another study, the genetic ablation of CDH1 caused the death of neural progenitor cells and microencephaly in mice (Delgado-Esteban et al. 2013). The same group showed that a missense mutation in CDH1 in a young boy was associated with severe antenatal microencephaly, psychomotor retardation, and refractory epilepsy (Rodríguez et al. 2019). Impaired CDH1 also leads to memory loss and impaired learning (Li et al. 2008; Bobo-Jimenez et al. 2017). Not surprisingly, APC^CDH1^ was linked with Alzheimer’s, as it has been shown that the amyloid-β peptide causes phosphorylation of CDH1, leading to APC^CDH1^ disassembly and loss of activity (Lapresa et al. 2022). Therefore, the need for methods to target activation of APC^CDH1^ in the aging brain cannot be understated.

In a recent study, using a yeast 2-hybrid screen, we discovered small peptides that bind to the yeast Apc10 subunit of the APC (VanGenderen et al. 2026). Apc10 is the processivity factor for the APC, required for the recruitment of substrates and the building of long K48-linked ubiquitin chains (Passmore et al. 2003; Buschhorn et al. 2011; Meyer and Rape 2011). Apc10, from yeast to humans, forms a co-receptor with Cdc20/Cdh1 to recruit substrates that contain a D-box motif, composed of the RxxL core sequence (da Fonseca et al. 2011; Qin et al. 2019). We chose to use Apc10 as our bait due to its pivotal role in recruiting and ubiquitinating substrate targets. Its broad interaction domains suggest that there is ample opportunity to manipulate Apc10 protein-protein interactions through small peptide binding.

We had shown previously that APC activity is impaired in multiple drug-resistant (MDR) breast cancer cells (Arnason et al. 2022; Lubachowski et al. 2024). We also showed using the APC binding peptides that MDR cancer could be reversed (VanGenderen et al. 2026). The peptides C13-3 and C43-4, stably expressed from the TrxA plasmid construct, demonstrated the best conserved activity of the peptides in human cells, showing a significant decrease in APC substrates. C13-3 and C43-4 both increased the pAPC1^S355^ activation signal, and APC E3 activity, and this was associated with increased DNA damage, apoptosis, and mitotic catastrophe. Both C13-3 and C43-4 resensitized MDR breast cancer cells growing in culture to DOX, but only C43-4 stalled tumor growth in a mouse model growing MDA-MB-231 triple negative breast cancer cells stably expressing HA-TrxA-C43-4. M2I-1 also stalled tumor growth in the mouse model and acted at least additively with C43-4, indicating that M2I-1 and C43-4 activate the APC through different mechanisms.

We hypothesized in the study presented here that the APC loses function in aging cells. The rationale for this hypothesis was based on our observations in yeast that APC mutants have shorter RLS and CLS (Harkness et al. 2004), and that the APC is responsible for chromatin assembly and histone post-translational modifications during mitosis (Harkness et al. 2002; Arnason et al. 2005; Harkness et al. 2005; Turner et al. 2010; Islam et al. 2011). We also observed that the APC worked with the Snf1/Fkh1 stress response pathway (Harkness et al. 2004; Postnikoff et al. 2012; Menzel et al. 2014; Malo et al. 2016; Harkness 2018), all indicative that the APC would impact longevity.

In this study, we validated this hypothesis by showing that the APC cannot properly target the APC substrates Clb1 or Mps1 when derived from aging cells (**Figure 1**). We tested whether APC^Cdc20^ activation using the M2I-1 SAC inhibitor could increase yeast lifespan. It increased RLS but it did not increase CLS (**Figure 2**). Since M2I-1 only works on APC^CDC20^, and entrance to quiescence requires APC^CDH1^ (Marescal and Cheeseman 2020), it was not surprising that M2I-1 did not increase CLS. C43-4 increased the RLS of WT cells (**Figure 5A**), and was the only peptide to increase the CLS of WT cells that we tested (**Figure 6**). C43-4 required *CDH1* to induce a slow growth phenotype and increase CLS (Figures **4C****, 4D-H**, **10C**). C43-4 was used to test whether it could maintain APC function in aging cells using the assay shown in **Figure 1**. Only when C43-4 was induced with galactose in aging cells did we observe the maintenance of low levels of Clb1-TAP (**Figure 7**), indicating that the APC remained active throughout the aging process.

We next synthesized the 20 amino acid C43-4 fused to an FITC motif and found that it could increase CLS by itself (**Figure 8**). This indicated that C43-4 did not require the TrxA backbone it was expressed from in the plasmid, nor did it require the restrained confirmation that was offered with the TrxA backbone. Using westerns and cellular fractionation, we observed that both the HA-TrxA-C43-4 construct and the synthetic C43-4^FITC^ peptide were localized to the nucleus (**Figures 3F**, **9A**). HA-TrxA-C43-4 was fused to a nuclear localization signal, but C43-4^FITC^ was not. We asked if C43-4^FITC^ required the cell penetrating TAT2 peptide to transport C43-4 into the cells, but C43-4^FITC^ increased the CLS of cells in the presence or absence of TAT2 (**Figure 8A**), and C43-4^FITC^ still entered the cell and the nucleus in the absence of the cell penetrating TAT2 peptide (**Figure 9A**). Use of the scrambled peptide did not increase CLS. However, while C43-4^FITC^ was observed in the nucleus, the scrambled^FITC^ peptide did not enter the cell (**Figure 9A**). Both C43-4 and scrambled^FITC^ are predicted to have a net positive charge of approximately 3 (bachem.com; biosynth.com) and to be amphipathic, which are features of cell-penetrating peptides (Svirina and Terterov 2021; Egberink et al. 2023). An enrichment of positively charged amino acids typically defines a nuclear localization signal. C43-4 and the scrambled peptide both contain 4 arginines, with three of them clustered in C43-4 (RVRR), but not in the scrambled peptide. The clustered stretch of arginines in C43-4 may be sufficient to allow C43-4 to cross the nuclear membrane. To test this, we created a D7A mutation in C43-4 and observed that it was capable of entering the nucleus (**Figure 9A**), but did not slow the growth of cells like WT C43-4, did not increase CLS, and did not interact with Apc10 in a 2-hybrid assay (**Figures 9C-E**). The C43-4^D7A^ mutant peptide provided the evidence we required to demonstrate that C43-4 increased yeast CLS and slowed growth via an interaction with Apc10. This was further supported by the observations that C43-4^FITC^ could not increase CLS in any APC mutant tested (**Figure 10**), confirming that C43-4 required a fully functional APC. C43-4 could not increase the CLS of any of the 3 APC co-activator mutants tested, *cdc20-1*, *cdh1Δ* or *ama1Δ*, indicating that C43-4 can activate all forms of the APC. Finally, both the FITC-tagged and untagged C43-4 versions increased CLS to the same extent (**Figure 9E**), demonstrating that the FITC motif does not impact CLS.

The C43-4^D7A^ mutation was discovered due to sequence homology between C43-4 and Htz1, deduced from a BLAST search (**Supplemental Figure 13A**). Six C43-4 amino acids were homologous with a sequence stretch observed in Htz1 (**Figure 9B**), a histone variant that responds to stress (Srivatsan et al. 2018; Saatchi et al. 2019; Brewis et al. 2025). All 6 amino acids were individually converted to alanines in C43-4. In addition to D7A creating a loss-of-function C43-4 version, the R12A and T14A mutations also reversed the slow growth phenotype of C43-4 (**Figure 9D**). C43-4^R12A^ and C43-4^T14A^ both retained the ability to interact with Apc10, whereas C43-4^D7A^ did not (**Figure 9C**). We synthesized the D7A mutant for our control, but future work will involve synthesizing the R12A and T14A variants. R12 was found to be conserved in all the homologous proteins identified using BLAST (**Supplemental Figure 13A**) and defined the R in the RxxL motif, highlighting the importance of this motif for C43-4 function. In addition, C43-4 was found to share homology with several other proteins, 2 of which (Cst9 and Rpt1), also contained an RxxL domain. Neither Cst9 nor Rpt1 have been reported to be unstable or to interact with the APC, suggesting these 2 proteins could be novel APC targets. We focused on Htz1 and observed genetic and functional interplay between Htz1 and the APC (**Supplemental Figures 13B-D**). Strikingly, we found that the Apc10 subunit is unstable when grown in galactose, but was stabilized when *HTZ1* was overexpressed. Furthermore, the Htz1 protein was weakly expressed, but accumulated in *apc5^CA^* and *apc10Δ* mutants, suggesting that Htz1 may be partially targeted by the APC. This suggests a novel means of controlling APC function. Reducing Apc10 levels in stressed cells could streamline APC activity to target the stress response rather than the cell cycle. Future work will investigate these ideas.

Ama1 is a third APC activator that is specific to yeast and thought to be active only in meiosis (Pesin and Orr-Weaver 2008; Cooper and Strich 2011). As discussed above, the exchangeable APC activators (one of Cdc20, Cdh1, or Ama1) and Apc10 form a 2-component substrate recognition module for substrate recruitment (Vazquez-Fernandez et al. 2024). The D-box, comprised of the base RxxL sequence, is a motif encoded by many APC substrates that is necessary for recognition by the Apc10 2-component complex (Carroll et al. 2005; Barford 2011; Morgan 2013; Barford 2020). Strikingly, the C43-4 peptide contains an RxxL motif (**Table 2**). A recent study showed that peptides that mimic the D-box motif can bind to Cdc20 and inhibit APC^Cdc20^ activity (Eapen et al. 2025). We do not fully understand the mechanism at work with C43-4, but we suspect that C43-4 may mitigate Apc10-co-activator binding, possibly in the absence of substrate, prematurely activating the APC. Alternatively, C43-4 could replace Cdc20 in the co-receptor. This possibility was suggested by our observation that C43-4 could inhibit large tumor growth in mice induced by APCIN (VanGenderen et al. 2026), the APC inhibitor that works by blocking recruitment of CDC20 to the APC. Therefore, C43-4 may work in the absence of CDC20.

We have shown that C43-4 works in human breast cancer cells by resensitizing MDR cells to DOX, both *in vitro* and *in vivo* (VanGenderen et al. 2026). Here, we demonstrated that the lifespan-extending properties of C43-4 are indeed conserved. C43-4 was fused to GFP and expressed within worms (**Figures 11A**, **11B**). We found that, similar to yeast, C43-4 extended the lifespan of the *C. elegans* worm (**Figure 11C**). As discussed above, the APC works within the SNF1/Fkh1 feedforward loop in yeast (Harkness et al. 2004; Postnikoff et al. 2012; Malo et al. 2016). To test if this pathway of APC control was also conserved in worms, we measured lifespan in worms expressing the AMPK mutant *aak-2* and the FOXO3A mutant *daf-16*. In both cases, the longer lifespan induced by C43-4 was lost (**Figures 11D, 11E**), showing that in worms, APC activation by C43-4 likely works within the same AMPK/FOXO feed-forward loop as in yeast. Lastly, mutation of the AKT orthologue, *akt-1*, resulted in long life that could not be extended further by C43-4 (**Figure 11F**). This suggests that removal of the conserved AKT nutrient sensing kinase is important for APC-dependent lifespan increases. Consistent with this, our unpublished data indicates that the APC targets the nutrient sensor Sch9 (AKT in humans) for ubiquitin- and proteasome-dependent degradation when stress is present (unpublished).

In conclusion, we have discovered the first direct APC^Cdh1^ activator using a yeast 2-hybrid screen where Apc10 was the bait. The C43-4 peptide is the only one to require a functional APC^Cdh1^, identifying this peptide as a possible therapeutic for use in maintaining the health of differentiated, quiescent non-dividing cells. Maintaining the health of quiescent cells will inhibit or delay the transition of these cells to senescent cells, which occurs upon the accumulation of damaged DNA and proteins as cells age. Senescence is considered a key hallmark of aging, suggesting that the inhibition or delay of senescence, such as that provided by the C43-4 peptide, could delay aging. Protecting the health of differentiated quiescent cells could also be vital to protect against the development of cancer in differentiated cells and against neurodegeneration in neurons and the brain. Characterizing how C43-4 interacts with the APC, and possibly directly with Apc10, will provide the blueprint for developing further therapeutics that can be used to delay again and the onset of age-related disease.

## Supporting information

Supplemental data and will be used for the link to the file on the preprint site.

## ACKNOWLEDGEMENTS

We thank Saruul Uuganbayar for early assistance in the work. Dylan Bunko is also thanked for assistance with experiments. Meet Patel constructed the pRS416-HA-TrxA-C43-4 and YEplac181-HA-TrxA-C43-4 plasmids. Andrew Emili (U of Toronto) generously provided the *GAL_prom_-GST-HTZ1* construct. REH was supported by a USask CoMGRAD award. HMB was supported by an NSERC summer studentship. AK was supported by a USask College of Medicine BioMed summer award.

## AUTHOR CONTRIBUTIONS

REH performed the experiments in **Figures 1**, **2A**, **2B**, **2E**, **3B**-**E, 3G**, **4**, **6**, **7,** and **9D**. REH also performed the stats for the CLS experiments in **Figures 2** and **6**, and the stats for the spot dilutions in **Figures 3B**, and **4**, as well as the stats shown in the **Supplemental Figures**. REH also contributed to manuscript editing. SDP performed the experiments in **Figures 8**, **9E**, and **10**, and the stats for the CLS curves in **Figures 8-10**. NKS developed the nuclear fractionation assay, performed the experiments in **Figures 3F** and **9A**, and contributed to manuscript editing. AHH performed all the stats for the RLS experiments shown in the Supplemental Figures. AK created the C43-4 alanine mutants and performed the 2-hybrid assay shown in **Figure 9C**, HMB performed the experiments shown in **Supplemental Figures 13B-D**, and MEM supervised HMB and AK. CZV performed the *C. elegans* lifespan assays and created mutant C43-4 strains. BMW made the initial C43-4 worm strain (cloning, microinjection, integration, outcross). MW designed the experiments performed in **Figure 11** and supervised CZV and BMW. MW also contributed to editing of the manuscript. TAAH performed the RLS experiments shown in **Figures 2** and **5**, designed the 2-hybrid screen and the yeast experiments, wrote the grants supporting the work, and wrote and edited the manuscript. TAAH supervised REH, SDP, NKS, AHH, AK, HMB and MEM.

## DATA AVAILABILITY

No new databases were generated in this study. Strains and plasmids are available upon request.

## REFERENCES

Almeida A, Bolanos JP, Moreno S. 2005. Cdh1/Hct1-APC is essential for the survival of postmitotic neurons. J Neurosci. 25:8115–8121.

Arnason TG, Pisclevich MG, Dash MD, Davies GF, Harkness TA. 2005. Novel interaction between Apc5p and Rsp5p in an intracellular signaling pathway in Saccharomyces cerevisiae. Eukaryot Cell. 4:134–46.

Arnason TG et al. 2022. Activation of the Anaphase Promoting Complex Reverses Multiple Drug Resistant Cancer in a Canine Model of Multiple Drug Resistant Lymphoma. Cancers. 14: 4215.

Akbergenov R, Wolfer DP, Gillingham D, Shcherbakov D. 2025. Error-prone translation as a driver of proteostasis collapse and neurodegeneration. Neural Regen Res. Dec 30.

Arnold L, Höckner S, Seufert W. 2015. Insights into the cellular mechanism of the yeast ubiquitin ligase APC/C-Cdh1 from the analysis of in vivo degrons. Mol Biol Cell. 26: 843–58.

Bapat P, Nandy SK, Wangikar P, Venkatesh KV. 2006. Quantification of metabolically active biomass using Methylene Blue dye Reduction Test (MBRT): measurement of CFU in about 200 s. J Microbiol Methods. 65: 107–16.

Barford D. 2011. Structural insights into anaphase-promoting complex function and mechanism. Philos Trans R Soc Lond B Biol Sci. 366: 3605–24.

Barford D. 2020. Structural interconversions of the anaphase-promoting complex/cyclosome (APC/C) regulate cell cycle transitions. Curr Opin Struct Biol. 61: 86–97.

Baumer M, Braus GH, Irniger S. 2000. Two different modes of cyclin Clb2 proteolysis during mitosis in *Saccharomyces cerevisiae*. FEBS Lett. 468: 142–148.

Bobo-Jimenez V, et al. 2017. APC/C(Cdh1)-Rock2 pathway controls dendritic integrity and memory. Proc Natl Acad Sci. 114: 4513–4518.

Brenner S. 1974. The genetics of *Caenorhabditis elegans*. Genetics. 77:71–94.

Brewis HT, Stirling PC, Kobor MS. 2025. Characterizing the regulatory effects of H2A.Z and SWR1-C on gene expression during hydroxyurea exposure in *Saccharomyces cerevisiae*. PLoS Genet. 21: e1011566.

Brown A, Geiger H. 2018. Chromosome integrity checkpoints in stem and progenitor cells: transitions upon differentiation, pathogenesis, and aging. Cell Mol Life Sci. 75: 3771–3779.

Buschhorn BA et al. 2011. Substrate binding on the APC/C occurs between the coactivator Cdh1 and the processivity factor Doc1. Nat Struct Mol Biol. 18: 6–13.

Cappell SD, Chung M, Jaimovich A, Spencer SL, Meyer T. 2016. Irreversible APC(Cdh1) Inactivation Underlies the Point of No Return for Cell-Cycle Entry. Cell. 166: 167–80.

Carroll CW, Enquist-Newman M, Morgan DO. 2005. The APC subunit Doc1 promotes recognition of the substrate destruction box. Curr Biol. 15: 11–8.

Chen M, Gutierrez GJ, Ronai ZA. 2012. The anaphase-promoting complex or cyclosome supports cell survival in response to endoplasmic reticulum stress. PLoS One 7: e35520.

Choi M et al. 2017. TC Mps1 12, a novel Mps1 inhibitor, suppresses the growth of hepatocellular carcinoma cells via the accumulation of chromosomal instability. Br J Pharmacol. 174: 1810–1825.

Cooper KF, Mallory MJ, Egeland DB, Jarnik M, Strich R. 2000. Ama1p is a meiosis-specific regulator of the anaphase promoting complex/cyclosome in yeast. Proc Natl Acad Sci. 97: 14548–14553.

Cooper KF, Strich R. 2011. Meiotic control of the APC/C: similarities & differences from mitosis. Cell Div. 6: 16.

Cui Y et al. 2010. Degradation of the human mitotic checkpoint kinase Mps1 is cell cycle-regulated by APC-cCdc20 and APC-cCdh1 ubiquitin ligases. J Biol Chem. 285, 32988–32998

da Fonseca PC et al. 2011. Structures of APC/C(Cdh1) with substrates identify Cdh1 and Apc10 as the D-box co-receptor. Nature. 470: 274–8.

Delgado-Esteban M, Garcia-Higuera I, Maestre C, Moreno S, Almeida A. 2013. APC/C-Cdh1 coordinates neurogenesis and cortical size during development. Nat Commun. 4: 2879.

Eapen R et al. 2025. Development of D-box peptides to inhibit the anaphase-promoting complex/cyclosome. Elife 14: RP104238.

Egberink RO et al. 2023. Deciphering Structural Determinants Distinguishing Active from Inactive Cell-Penetrating Peptides for Cytosolic mRNA Delivery. Bioconjug Chem. 34: 1822–1834.

Eguren M et al. 2013. The APC/C cofactor Cdh1 prevents replicative stress and p53-dependent cell death in neural progenitors. Nat Commun. 4: 2880.

Fabrizio P, Pozza F, Pletcher SD, Gendron CM, Longo VD. 2001. Regulation of longevity and stress resistance by Sch9 in yeast. Science 292: 288–90.

Ferguson CJ et al. 2022. APC7 mediates ubiquitin signaling in constitutive heterochromatin in the developing mammalian brain. Mol Cell 82: 90–105.e13.

García-Higuera I et al. 2008. Genomic stability and tumor suppression by the APC/C cofactor Cdh1. Nat Cell Biol. 10: 802–811.

Garzón J et al. 2017. Shortage of dNTPs underlies altered replication dynamics and DNA breakage in the absence of the APC/C cofactor Cdh1. Oncogene. 36: 5808–5818.

Gietz RD, Schiestl RH. 2007. High-Efficiency Yeast Transformation Using the LiAc/SS Carrier DNA/PEG Method. Nat Protocols. 2: 31–34.

Greil C et al. 2016. The role of APC/C(Cdh1) in replication stress and origin of genomic instability. Oncogene. 35: 3062–70.

Greil C, Engelhardt M, Wäsch R. 2022. The Role of the APC/C and Its Coactivators Cdh1 and Cdc20 in Cancer Development and Therapy. Front Genet. 13: 941565

Guo J et al. 2022. Aging and aging-related diseases: from molecular mechanisms to interventions and treatments. Signal Transduct Target Ther. 7: 391.

Gutierrez JI, Edgar C, Tyler JK. 2026. Overexpression of Ssd1 and calorie restriction extend yeast replicative lifespan by preventing deleterious age-dependent iron uptake. Elife. 12: 14:RP108892.

Ha K et al. 2017. The anaphase promoting complex impacts repair choice by protecting ubiquitin signalling at DNA damage sites. Nat Commun. 8: 15751.

Han SK et al.2016. OASIS 2: online application for survival analysis 2 with features for the analysis of maximal lifespan and healthspan in aging research. Oncotarget. 7: 56147–56152.

Han SK, Kwon HC, Yang JS, Kim S, Lee SV. 2024. OASIS portable: User-friendly offline suite for secure survival analysis. Mol Cells. 47: 100011.

Harkness TA, Davies GF, Ramaswamy V, Arnason TG. 2002. The ubiquitin-dependent targeting pathway in Saccharomyces cerevisiae plays a critical role in multiple chromatin assembly regulatory steps. Genetics. 162: 615–32.

Harkness TA, Shea KA, Legrand C, Brahmania M, Davies GF. 2004. A functional analysis reveals dependence on the anaphase-promoting complex for prolonged life span in yeast. Genetics. 168: 759–74.

Harkness TA et al. 2005. Contribution of CAF-I to anaphase-promoting-complex-mediated mitotic chromatin assembly in *Saccharomyces cerevisiae*. Eukaryot Cell. 4: 673–84.

Harkness TAA. 2018. Activating the Anaphase Promoting Complex to Enhance Genomic Stability and Prolong Lifespan. Int J Mol Sci. 19: 1888.

Hobert O. 2002. PCR fusion-based approach to create reporter gene constructs for expression analysis in transgenic C. elegans. Biotechniques. 32: 728–30.

Hu D, Qiao X, Wu G, Wan Y. 2011. The emerging role of APC/CCdh1 in development. Semin Cell Dev Biol. 22: 579–85.

Hu X, Jin X, Cao X, Liu B. 2022. The Anaphase-Promoting Complex/Cyclosome Is a Cellular Ageing Regulator. Int J Mol Sci. 23: 15327.

Irniger S, Bäumer M, Braus GH. 2000. Glucose and ras activity influence the ubiquitin ligases APC/C and SCF in Saccharomyces cerevisiae. Genetics. 154: 1509–21.

Islam A, Turner EL, Menzel J, Malo ME, Harkness TA. 2011. Antagonistic Gcn5-Hda1 interactions revealed by mutations to the Anaphase Promoting Complex in yeast. Cell Div. 6: 13.

Isogai S, Tanahashi R, Takagi H, Nishimura A. 2025. Evaluation of synthetic peptides for direct protein delivery into *Saccharomyces cerevisiae*. Biosci Biotechnol Biochem. 89: 1525–1528.

Jeong Y et al. 2023. Targeting E3 ubiquitin ligases and their adaptors as a therapeutic strategy for metabolic diseases. Exp Mol Med. 55: 2097–2104.

Jiao R, Postnikoff S, Harkness TA, Arnason TG. 2015. The SNF1 Kinase Ubiquitin-associated Domain Restrains Its Activation, Activity, and the Yeast Life Span. J Biol Chem. 290: 15393–15404.

Johnston M, Flick JS, Pexton T. 1994. Multiple mechanisms provide rapid and stringent glucose repression of GAL gene expression in *Saccharomyces cerevisiae*. Mol Cell Biol. 14: 3834–41.

Kapanidou M, Curtis NL, Bolanos-Garcia VM. 2017. Cdc20: At the Crossroads between Chromosome Segregation and Mitotic Exit. Trends Biochem Sci. 42: 193–205.

Kastl J et al. 2015. Mad2 Inhibitor-1 (M2I-1): A Small Molecule Protein-Protein Interaction Inhibitor Targeting the Mitotic Spindle Assembly Checkpoint. ACS Chem Biol. 10: 1661–6.

Kaushik S, Cuervo A. 2015. Proteostasis and aging. Nat Med. 21: 1406–1415.

Kennedy BK et al. 2014. Geroscience: linking aging to chronic disease. Cell. 159: 709–13.

Kevei É, Pokrzywa W, Hoppe T. 2017. Repair or destruction-an intimate liaison between ubiquitin ligases and molecular chaperones in proteostasis. FEBS Lett. 591: 2616–2635.

Kotani S et al. 1998. PKA and MPF-activated polo-like kinase regulate anaphase-promoting complex activity and mitosis progression. Mol Cell 1: 371–80.

Kwolek-Mirek M, Zadrag-Tecza R. 2014. Comparison of methods used for assessing the viability and vitality of yeast cells. FEMS Yeast Res. 14: 1068–79.

Ladner CL, Yang J, Turner RJ, Edwards RA. 2004. Visible fluorescent detection of proteins in polyacrylamide gels without staining. Anal Biochem. 326: 13–20.

Lafranchi L et al. 2014. APC/C(Cdh1) controls CtIP stability during the cell cycle and in response to DNA damage. EMBO J. 33: 2860–79.

Lapresa R, Agulla J, Bolaños JP, Almeida A. 2022. APC/C-Cdh1-targeted substrates as potential therapies for Alzheimer’s disease. Front Pharmacol. 13: 1086540.

Li M et al. 2008. The adaptor protein of the anaphase promoting complex Cdh1 is essential in maintaining replicative lifespan and in learning and memory. Nat Cell Biol. 10: 1083–1089.

Li Y et al. 2018. Impact of Healthy Lifestyle Factors on Life Expectancies in the US Population. Circulation. 138: 345–355.

Li J et al. 2019. M2I-1 disrupts the in vivo interaction between CDC20 and MAD2 and increases the sensitivities of cancer cell lines to anti-mitotic drugs via MCL-1s. Cell Div. 15: 14:5.

Lima I, Borges F, Pombinho A, Chavarria D. 2025. The spindle assembly checkpoint: Molecular mechanisms and kinase-targeted drug discovery. Drug Discov Today. 30: 104355.

Lõoke M, Kristjuhan K, Kristjuhan A. 2011. Extraction of genomic DNA from yeasts for PCR-based applications. Biotechniques. 50: 325–8.

López-Otín C, Blasco MA, Partridge L, Serrano M, Kroemer G. 2023. Hallmarks of aging: an expanding universe. Cell. 186: 243–278.

Lubachowski M et al. 2024. Activation of the Anaphase Promoting Complex Restores Impaired Mitotic Progression and Chemosensitivity in Multiple Drug-Resistant Human Breast Cancer. Cancers. 16: 1755.

Malo ME, Postnikoff SD, Arnason TG, Harkness TA. 2016. Mitotic degradation of yeast Fkh1 by the Anaphase Promoting Complex is required for normal longevity, genomic stability and stress resistance. Aging. 8: 810–30.

Marescal O, Cheeseman IM. 2020. Cellular Mechanisms and Regulation of Quiescence. Dev Cell. 55: 259–271.

Menzel J et al. 2014. The anaphase promoting complex regulates yeast lifespan and rDNA stability by targeting Fob1 for degradation. Genetics. 196: 693–709.

Meyer HJ, Rape M. 2011. Processive ubiquitin chain formation by the anaphase-promoting complex. Semin. Cell Dev. Biol. 22: 544–50.

Morgan DO. 2013. The D box meets its match. Mol Cell. 50: 609–10.

Morimoto RI, Cuervo AM. 2014. Proteostasis and the aging proteome in health and disease. J Gerontol A Biol Sci Med Sci. 69: Suppl 1, S33–8.

Mouery BL et al. 2014. APC/C prevents a noncanonical order of cyclin/CDK activity to maintain CDK4/6 inhibitor-induced arrest. Proc Natl Acad Sci. 121: e2319574121.

Neumann NP, Lampen JO. 1967. Purification and properties of yeast invertase. Biochemistry. 6: 468–75.

Niccoli T, Partridge L. 2012. Ageing as a risk factor for disease. Curr Biol. 22: R741–52.

Novais P, Silva PMA, Amorim I, Bousbaa H. 2021. Second-Generation Antimitotics in Cancer Clinical Trials. Pharmaceutics. 13:1011.

Olshansky SJ, Willcox BJ, Demetrius L. 2024. Beltrán-Sánchez H. Implausibility of radical life extension in humans in the twenty-first century. Nat Aging. 4: 1635–1642.

Orlicky S, Tang X, Willems A, Tyers M, Sicheri F. 2003. Structural basis for phosphodependent substrate selection and orientation by the SCF^Cdc4^ ubiquitin ligase. Cell. 112: 243–56.

Ovadya Y et al. 2018. Impaired immune surveillance accelerates accumulation of senescent cells and aging. Nat Commun. 9: 5435.

Passmore LA et al. 2003. Doc1 mediates the activity of the anaphase-promoting complex by contributing to substrate recognition. EMBO J. 22: 786–96.

Pesin JA, Orr-Weaver TL. 2008. Regulation of APC/C activators in mitosis and meiosis. Annu Rev Cell Dev Biol. 24: 475–99.

Petropavlovskiy AA, Tauro MG, Lajoie P, Duennwald ML. 2020. A Quantitative Imaging-Based Protocol for Yeast Growth and Survival on Agar Plates. STAR Protoc. 1: 100182.

Pinto B, Silva JPN, Silva PMA, Barbosa DJ, Sarmento B, Tavares JC, Bousbaa H. 2023. Maximizing Anticancer Response with MPS1 and CENPE Inhibition Alongside Apoptosis Induction. Pharmaceutics. 16:56.

Postnikoff SD, Malo ME, Wong B, Harkness TA. 2012. The yeast forkhead transcription factors Fkh1 and Fkh2 regulate lifespan and stress response together with the anaphase-promoting complex. PLoS Genet. 8: e1002583.

Postnikoff SD, Harkness TA. 2014. Replicative and chronological life-span assays. Methods Mol Biol. 1163: 223–7.

Qin L et al. 2019. The pseudosubstrate inhibitor Acm1 inhibits the anaphase-promoting complex/cyclosome by combining high-affinity activator binding with disruption of Doc1/Apc10 function. J Biol Chem. 294: 17249–17261.

Reinders A, Bürckert N, Boller T, Wiemken A, De Virgilio C. 1998. *Saccharomyces cerevisiae* cAMP-dependent protein kinase controls entry into stationary phase through the Rim15p protein kinase. Genes Dev. 12: 2943–55.

Rodríguez C et al. 2019. A novel human Cdh1 mutation impairs anaphase promoting complex/cyclosome activity resulting in microcephaly, psychomotor retardation, and epilepsy. J Neurochem. 151: 103–115.

Saatchi F, Kirchmaier AL. 2019. Tolerance of DNA Replication Stress Is Promoted by Fumarate Through Modulation of Histone Demethylation and Enhancement of Replicative Intermediate Processing in *Saccharomyces cerevisiae*. Genetics. 212: 631–654.

Sackton KL et al. 2014. Synergistic blockade of mitotic exit by two chemical inhibitors of the APC/C. Nature. 514: 646–9.

Sampaio-Marques B, Ludovico P. 2018. Linking cellular proteostasis to yeast longevity. FEMS Yeast Res. 18.

Sanada F, Hayashi S, Morishita R. 2025. Targeting the hallmarks of aging: mechanisms and therapeutic opportunities. Front Cardiovasc Med. 12: 1631578.

Sansregret L et al. 2017. APC/C Dysfunction Limits Excessive Cancer Chromosomal Instability. Cancer Discov. 7: 218–233.

Schneider-Poetsch T et al. 2010. Inhibition of eukaryotic translation elongation by cycloheximide and lactimidomycin. Nat Chem Biol. 6: 209–217.

Skaar JR, Pagano M. 2008. Cdh1: a master G0/G1 regulator. Nat Cell Biol. 10: 755–7.

Song S, Lam EW, Tchkonia T, Kirkland JL, Sun Y. 2020. Senescent Cells: Emerging Targets for Human Aging and Age-Related Diseases. Trends Biochem Sci. 45: 578–592.

Srivatsan A et al. 2018. The Swr1 chromatin-remodeling complex prevents genome instability induced by replication fork progression defects. Nat Commun. 9: 3680.

Svirina A, Terterov I. 2021. Electrostatic effects in saturation of membrane binding of cationic cell-penetrating peptide. Eur Biophys J. 50: 15–23.

Szymiczek A et al. 2017. Inhibition of the spindle assembly checkpoint kinase Mps-1 as a novel therapeutic strategy in malignant mesothelioma. Oncogene. 36: 6501–6507.

Tabarraei H et al. 2023. CCR4-NOT subunit CCF-1/CNOT7 promotes transcriptional activation to multiple stress responses in Caenorhabditis elegans. Aging Cell. 22: e13795.

Thu KL et al. 2018. Disruption of the anaphase-promoting complex confers resistance to TTK inhibitors in triple-negative breast cancer. Proc Natl Acad Sci. 115: E1570–E1577.

Torchilin VP. 2008. Tat peptide-mediated intracellular delivery of pharmaceutical nanocarriers. Adv Drug Deliv Rev. 60: 548–58.

Turner EL et al. 2010. The *Saccharomyces cerevisiae* anaphase-promoting complex interacts with multiple histone-modifying enzymes to regulate cell cycle progression. Eukaryot Cell. 9: 1418–31.

Upadhyay A, Joshi V. 2024. Proteasome Activators and Ageing: Restoring Proteostasis Using Small Molecules. Subcell Biochem. 107: 21–41.

VanGenderen C, Harkness TAA, Arnason TG. 2020. The role of Anaphase Promoting Complex activation, inhibition and substrates in cancer development and progression. Aging. 12: 15818–15855.

VanGenderen C, et al. 2026. Restoration of chemosensitivity to drug resistant breast cancer cells through peptide activation of the Anaphase Promoting Complex. Sci Rep. in press.

Vazquez-Fernandez E et al. 2024. A comparative study of the cryo-EM structures of *Saccharomyces cerevisiae* and human anaphase-promoting complex/cyclosome (APC/C). Elife. 13: RP100821.

Visintin R, Prinz S, Amon A. 1997. CDC20 and CDH1: a family of substrate-specific activators of APC-dependent proteolysis. Science. 278: 460–3.

Wäsch R, Robbins JA, Cross FR. 2010. The emerging role of APC/CCdh1 in controlling differentiation, genomic stability and tumor suppression. Oncogene 29: 1–10.

Yamashita YM et al. 1996. 20S cyclosome complex formation and proteolytic activity inhibited by the cAMP/PKA pathway. Nature. 384: 276–9.

Zhang Y et al. 2025. APC/C coactivators Cdh1 and Cdc20: Mechanistic insights into cancer progression and therapeutic opportunities. Cell Signal. 136: 112171.

Zhou Z, He M, Shah AA, Wan Y. 2016. Insights into APC/C: from cellular function to diseases and therapeutics. Cell Div. 11: 9.

Zhou X, Xu R, Wu Y, Zhou L, Xiang T. 2023. The role of proteasomes in tumorigenesis. Genes Dis. 11: 101070.

