## Supplemental data and will be used for the link to the file on the preprint site. for "Discovery of a small peptide that increases yeast lifespan by enhancing the function of APC^Cdh1^ in nondividing quiescent cells"

### Supplemental Figure 1

Step 1 Run two PCR reactions using the pJG4-5 vector with peptide construct cloned in for Primers 1 & 2, and genomic DNA with a functional *LEU2* gene for Primers 3 & 4

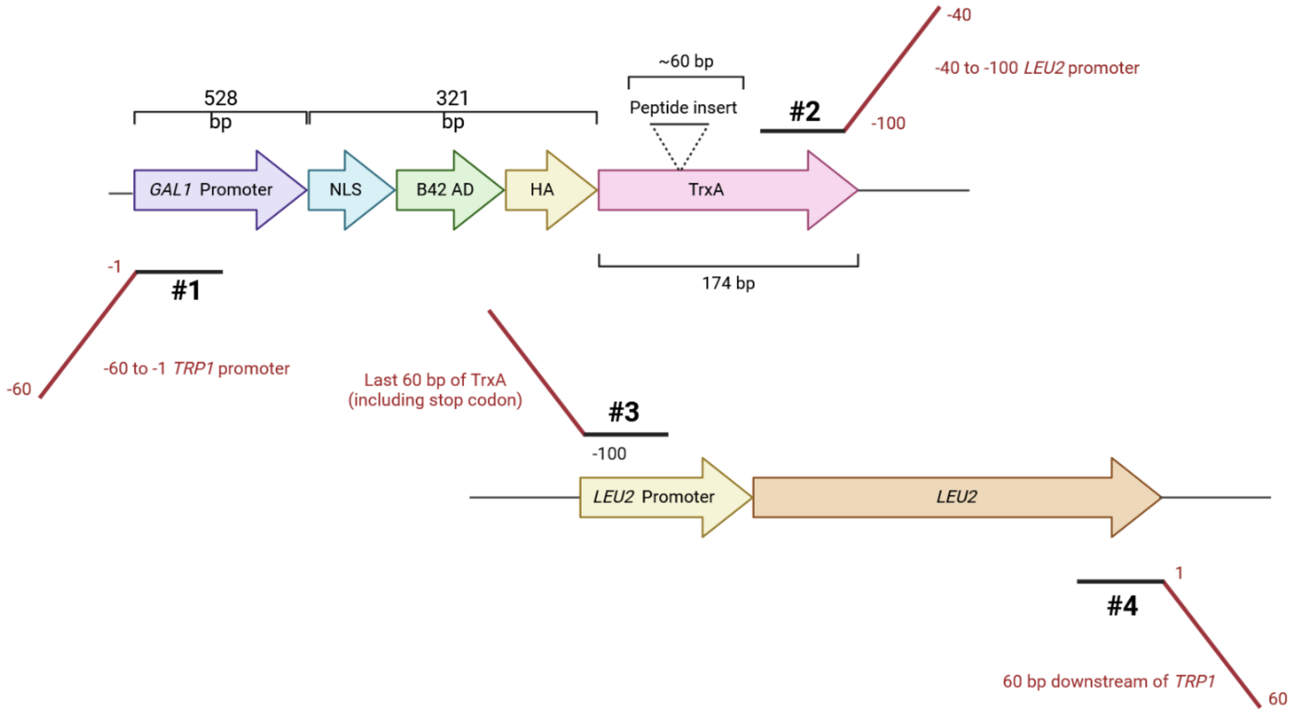

Step 2 Transform both PCR products into a strain lacking a functional *LEU2* gene

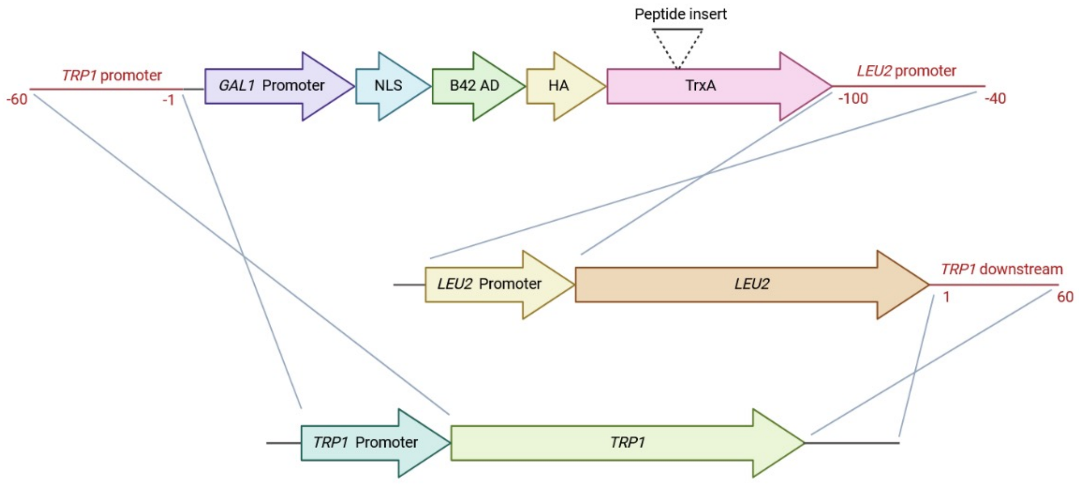

### Supplemental Figure 2

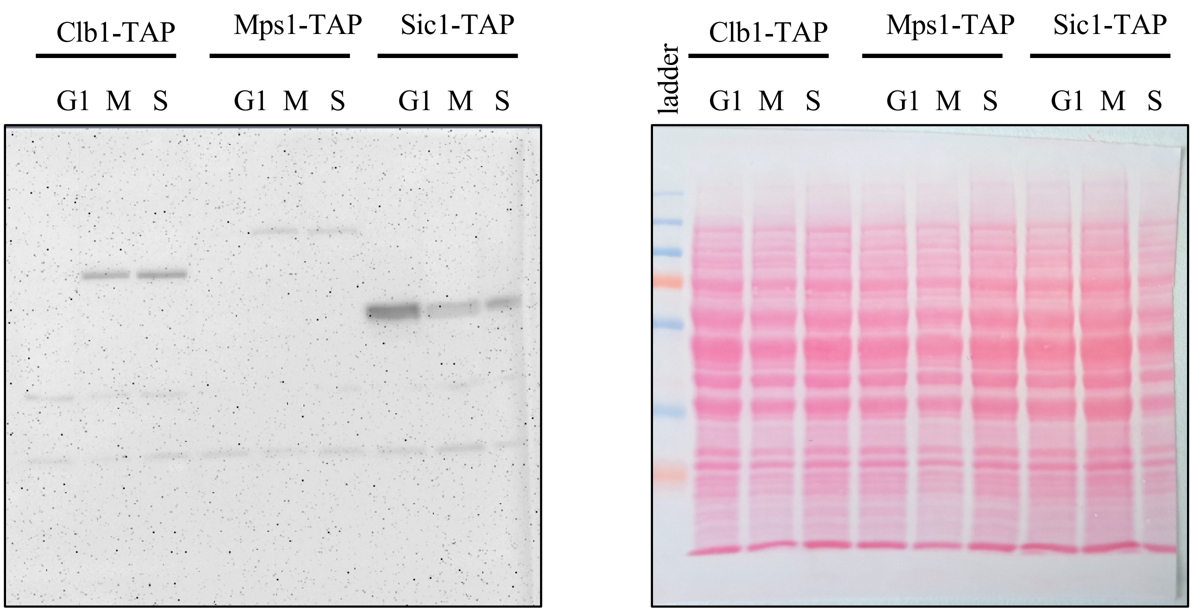

### Supplemental Figure 3

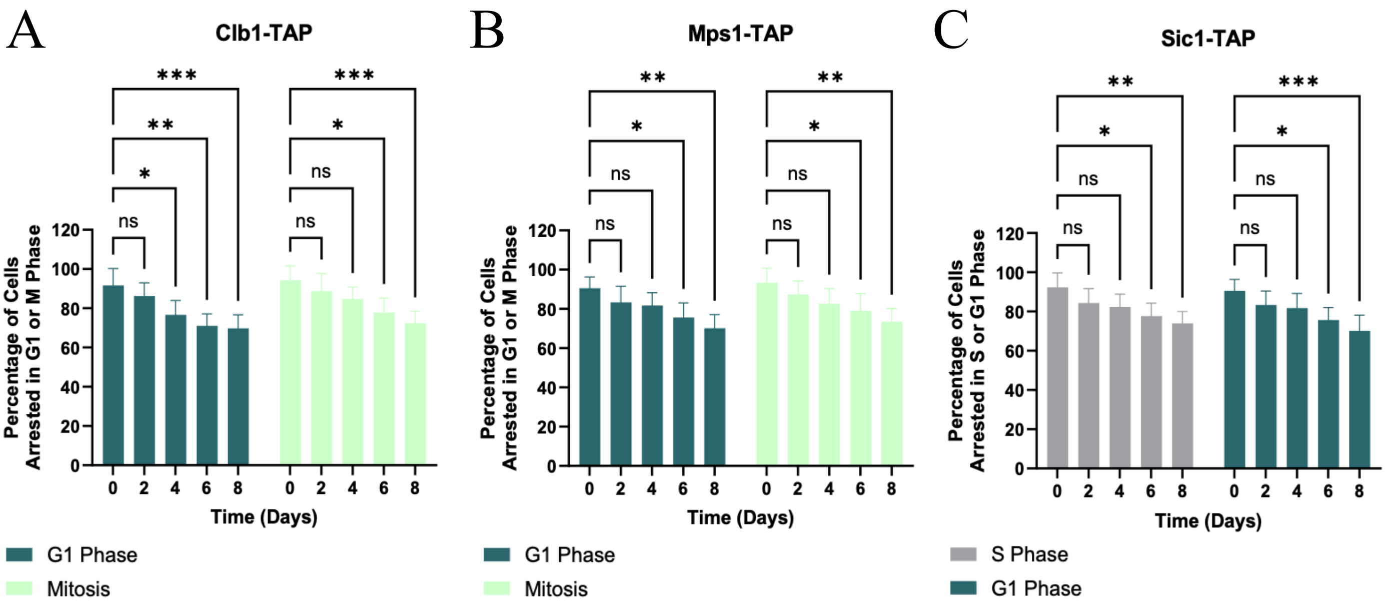

### Supplemental Figure 4

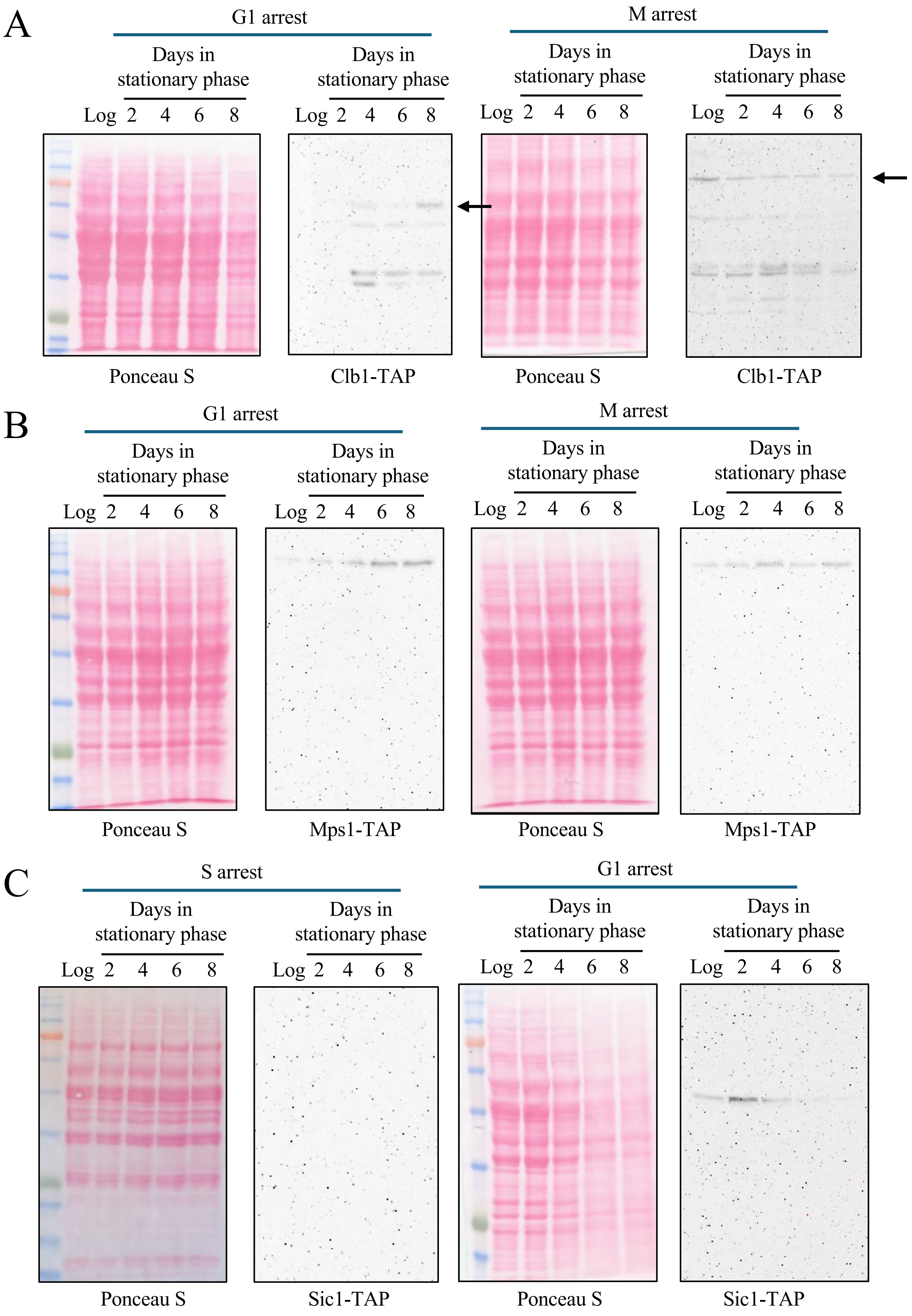

### Supplemental Figure 5

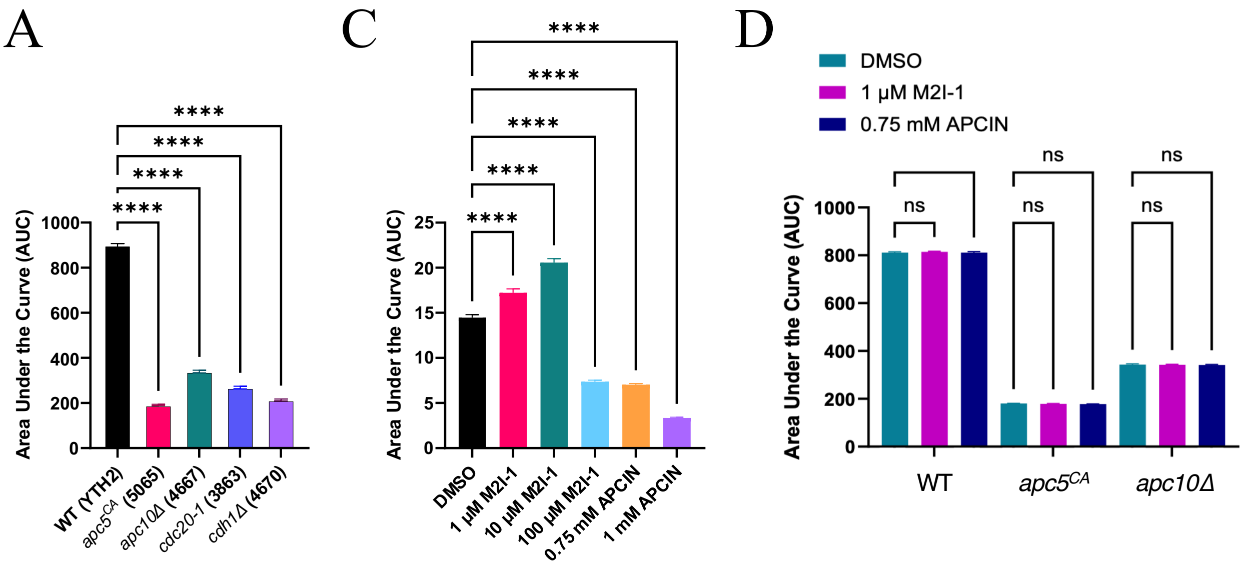

B

| Strain | Mean Survival (Days) | 95% Confidence Interval | Max survival <10% (Days) |
|---|---|---|---|
| **WT** | 13.26 ± 0.09225 | 12.93 to 13.61 | 23 |
| ***apc5$^{CA}$*** | 3.80 ± 0.6632 | 3.606 to 3.988 | 7 |
| ***apc10Δ*** | 6.25 ± 0.3796 | 5.558 to 6.912 | 15 |
| ***cdc20-1*** | 5.20 ± 0.5098 | 4.520 to 5.819 | 13 |
| ***cdh1Δ*** | 4.02 ± 0.5953 | 3.830 to 4.219 | 7 |

### Supplemental Figure 6

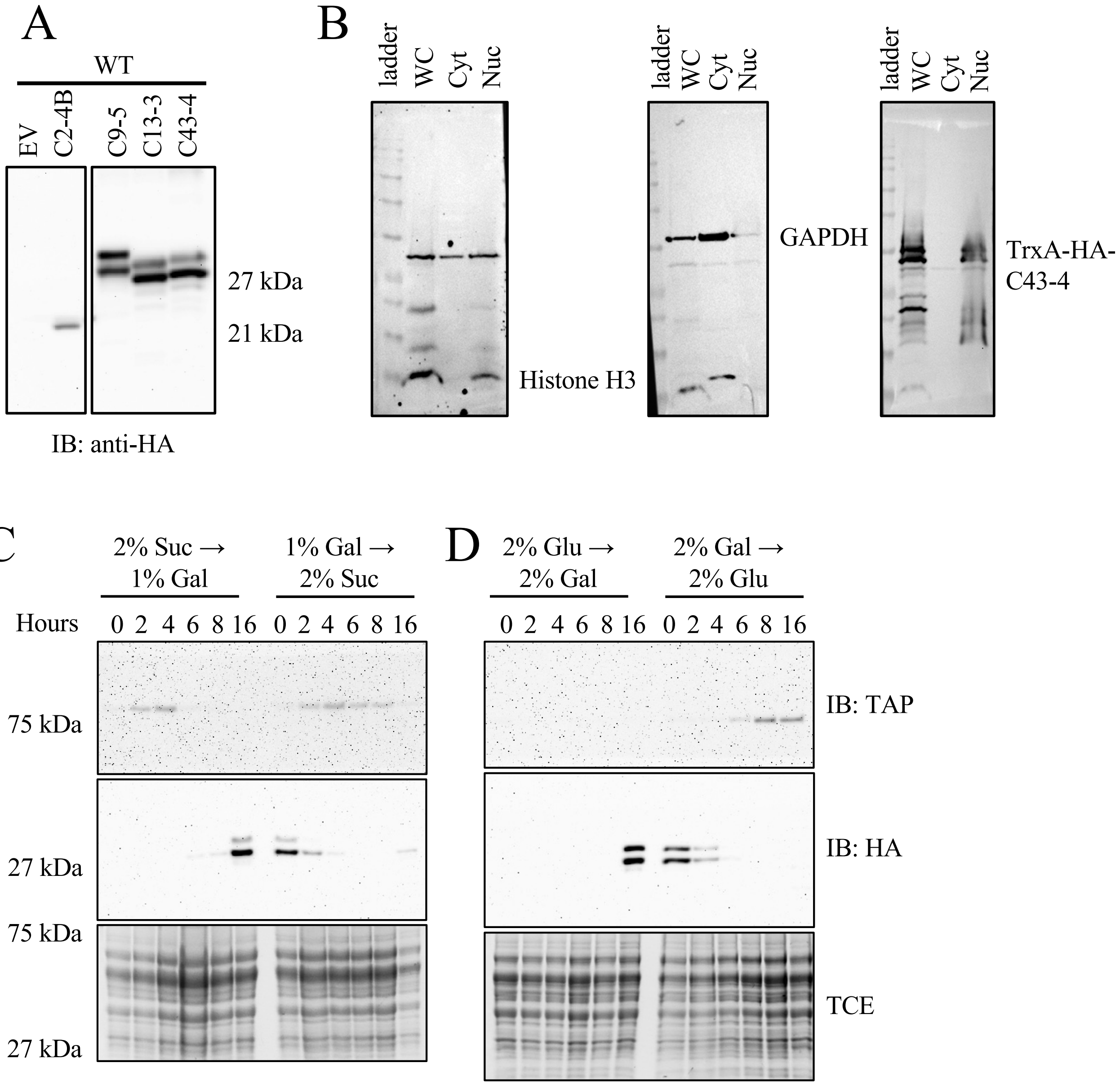

### Supplemental Figure 7

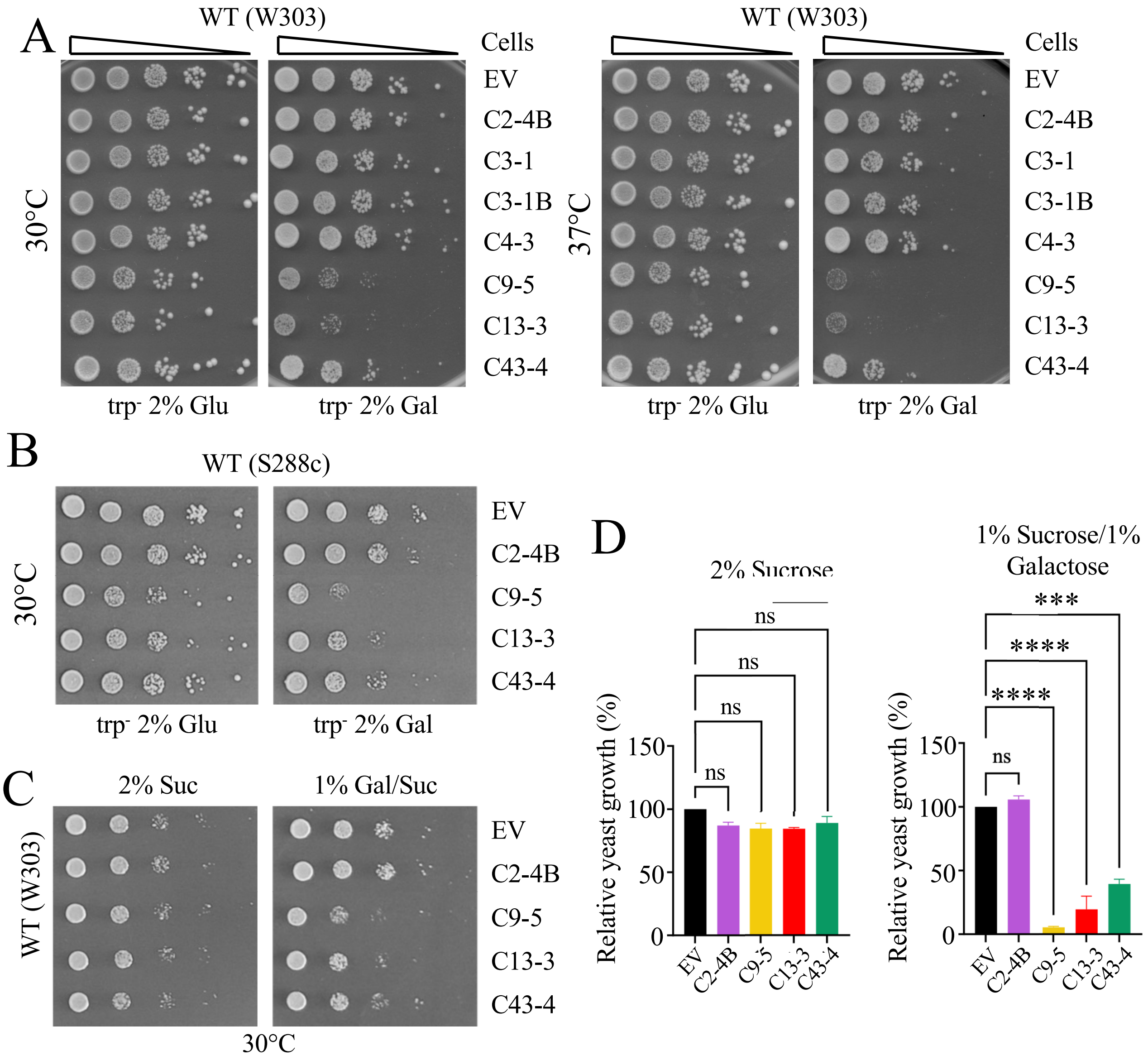

### Supplemental Figure 8

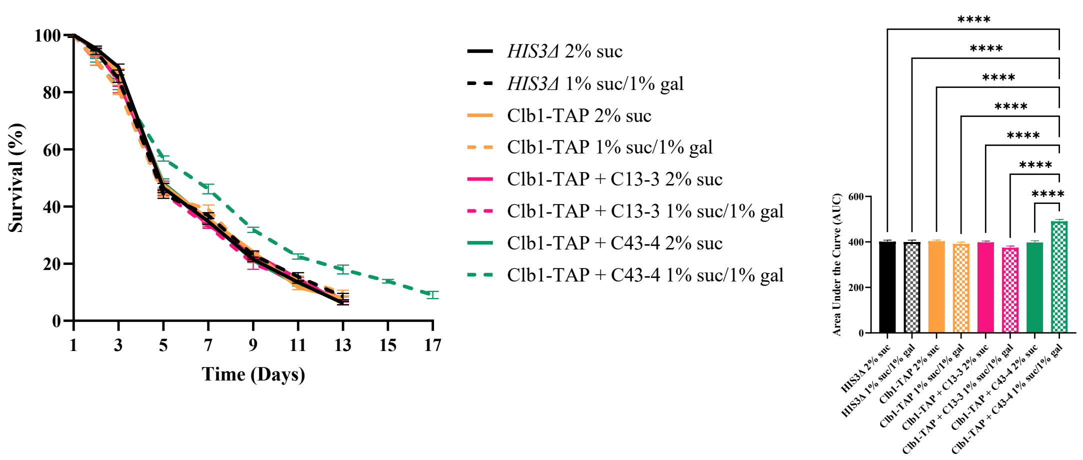

### Supplemental Figure 9

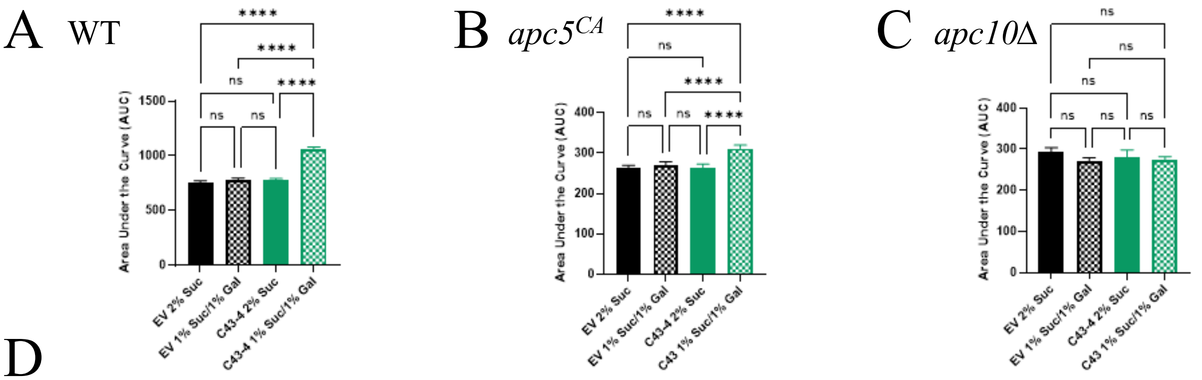

D

| Strain | Mean Survival (Days) | 95% Confidence Interval | Max survival (Days) |
|---|---|---|---|
| **WT EV 2% Suc** | 10.73 ± 0.1151 | 10.41 to 11.06 | 21 |
| **WT EV 1% Gal** | 10.92 ± 0.1049 | 10.62 to 11.22 | 21 |
| **WT C43 2% Suc** | 10.92 ± 0.1171 | 10.59 to 11.25 | 21 |
| **WT C43 1% Gal** | 14.38 ± 0.05722 | 14.10 to 14.67 | 27 |
| ***apc5$^{CA}$* EV 2% Suc** | 4.209 ± 0.7631 | 3.977 to 4.445 | 7 |
| ***apc5$^{CA}$* EV 1% Gal** | 4.254 ± 0.8992 | 3.977 to 4.539 | 7 |
| ***apc5$^{CA}$* C43 2% Suc** | 4.303 ± 0.8374 | 4.041 to 4.574 | 7 |
| ***apc5$^{CA}$* C43 1% Gal** | 5.011 ± 0.9553 | 4.538 to 5.485 | 9 |
| ***apc10Δ* EV 2% Suc** | 4.834 ± 0.4333 | 4.611 to 5.062 | 9 |
| ***apc10Δ* EV 1% Gal** | 4.634 ± 0.4421 | 4.418 to 4.852 | 9 |
| ***apc10Δ* C43 2% Suc** | 4.577 ± 0.5863 | 4.298 to 4.857 | 9 |
| ***apc10Δ* C43 1% Gal** | 4.659 ± 0.5458 | 4.384 to 4.937 | 9 |

Supplemental Figure 10

A

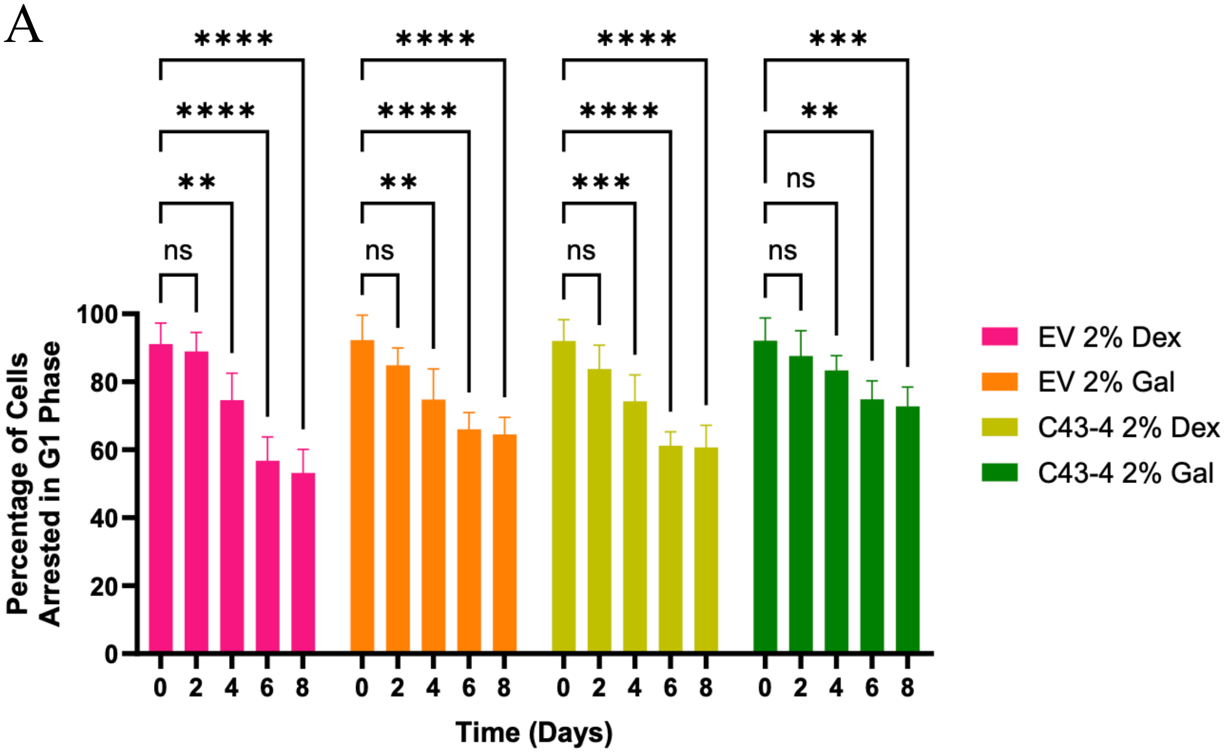

B

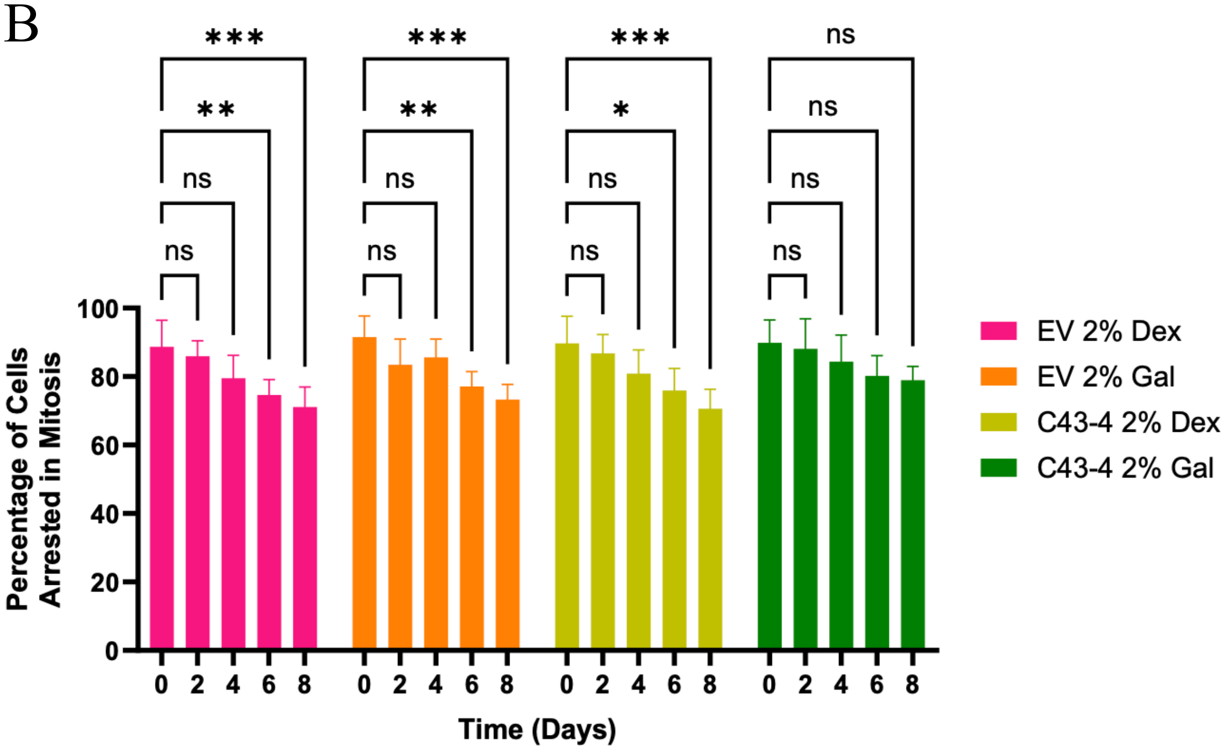

### Supplemental Figure 11

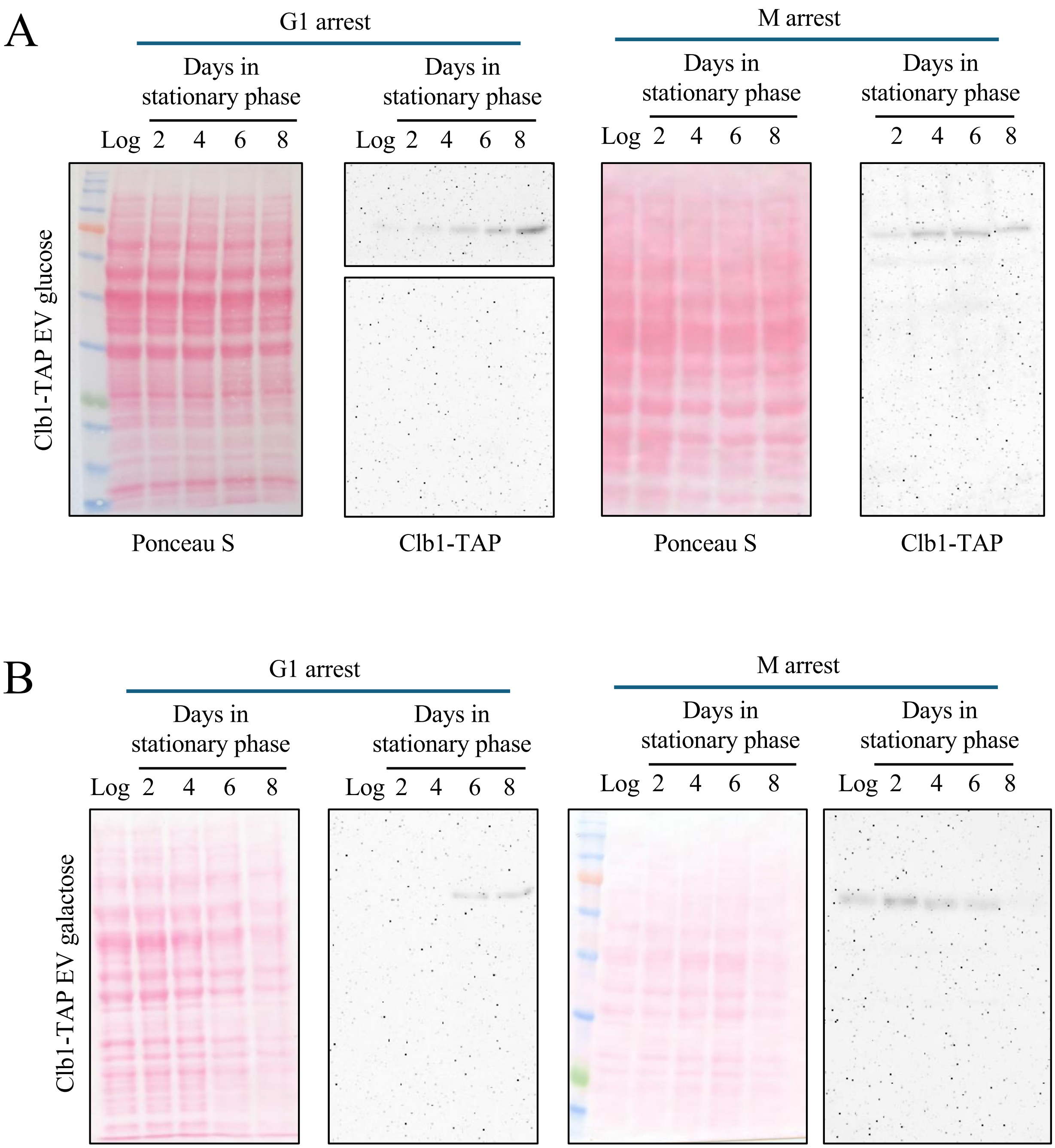

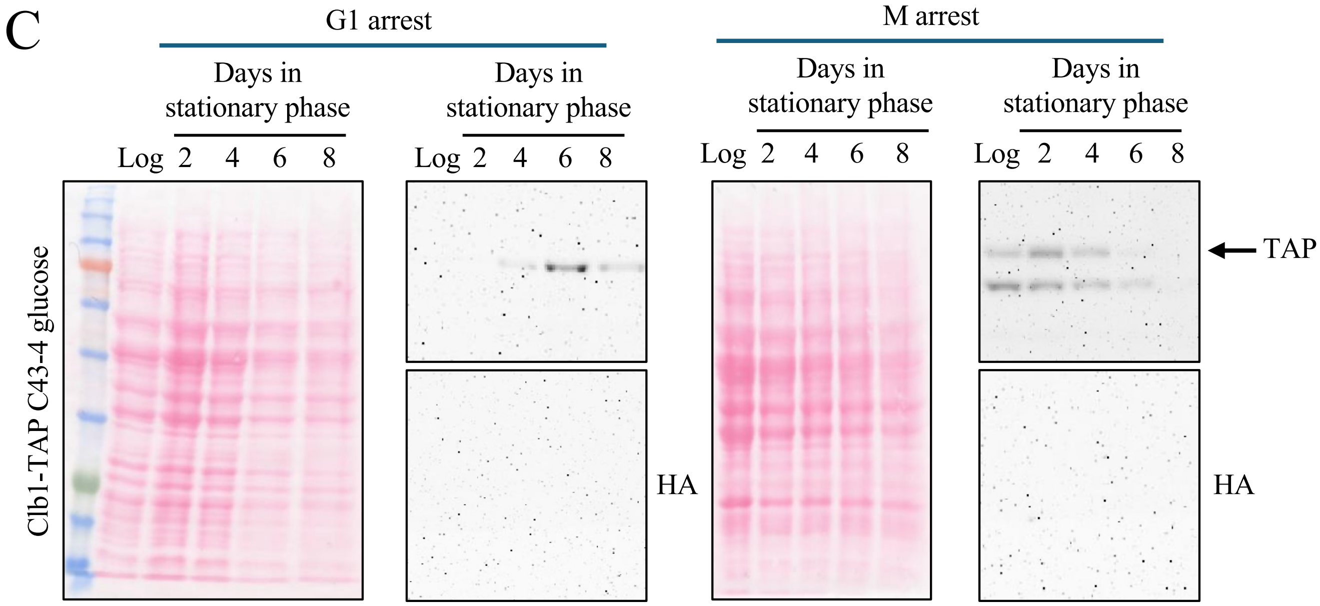
C
G1 arrest
M arrest
Days in stationary phase
Log 2 4 6 8
Clb1-TAP C43-4 glucose
TAP
HA

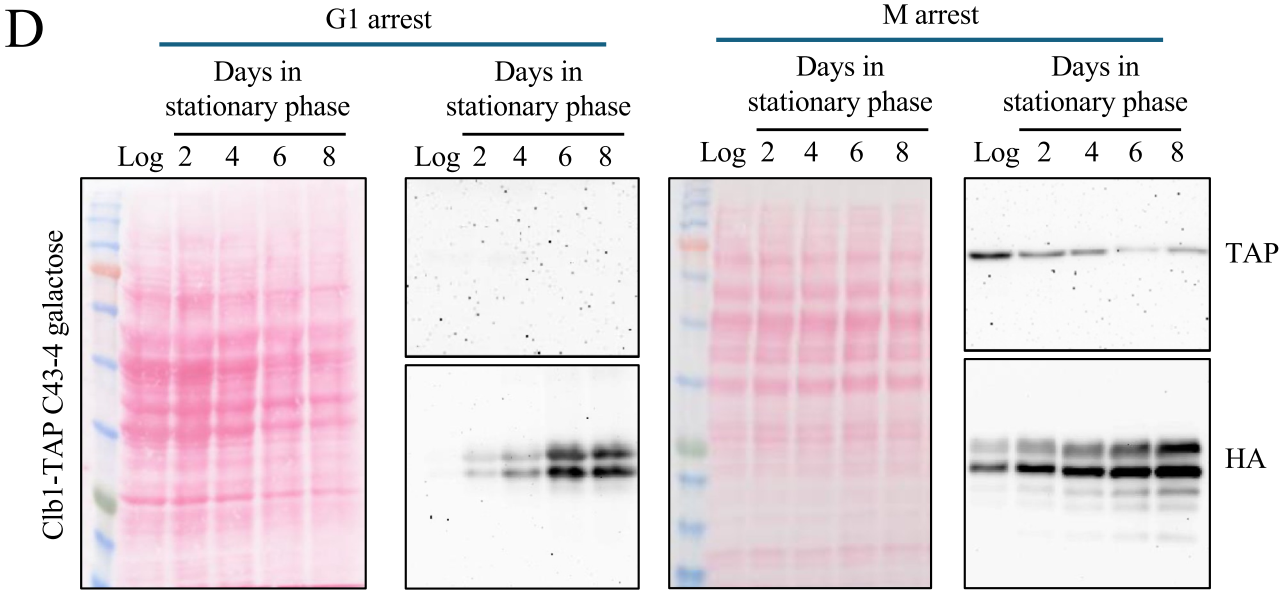
D
G1 arrest
M arrest
Days in stationary phase
Log 2 4 6 8
Clb1-TAP C43-4 galactose
TAP
HA

### Supplemental Figure 12

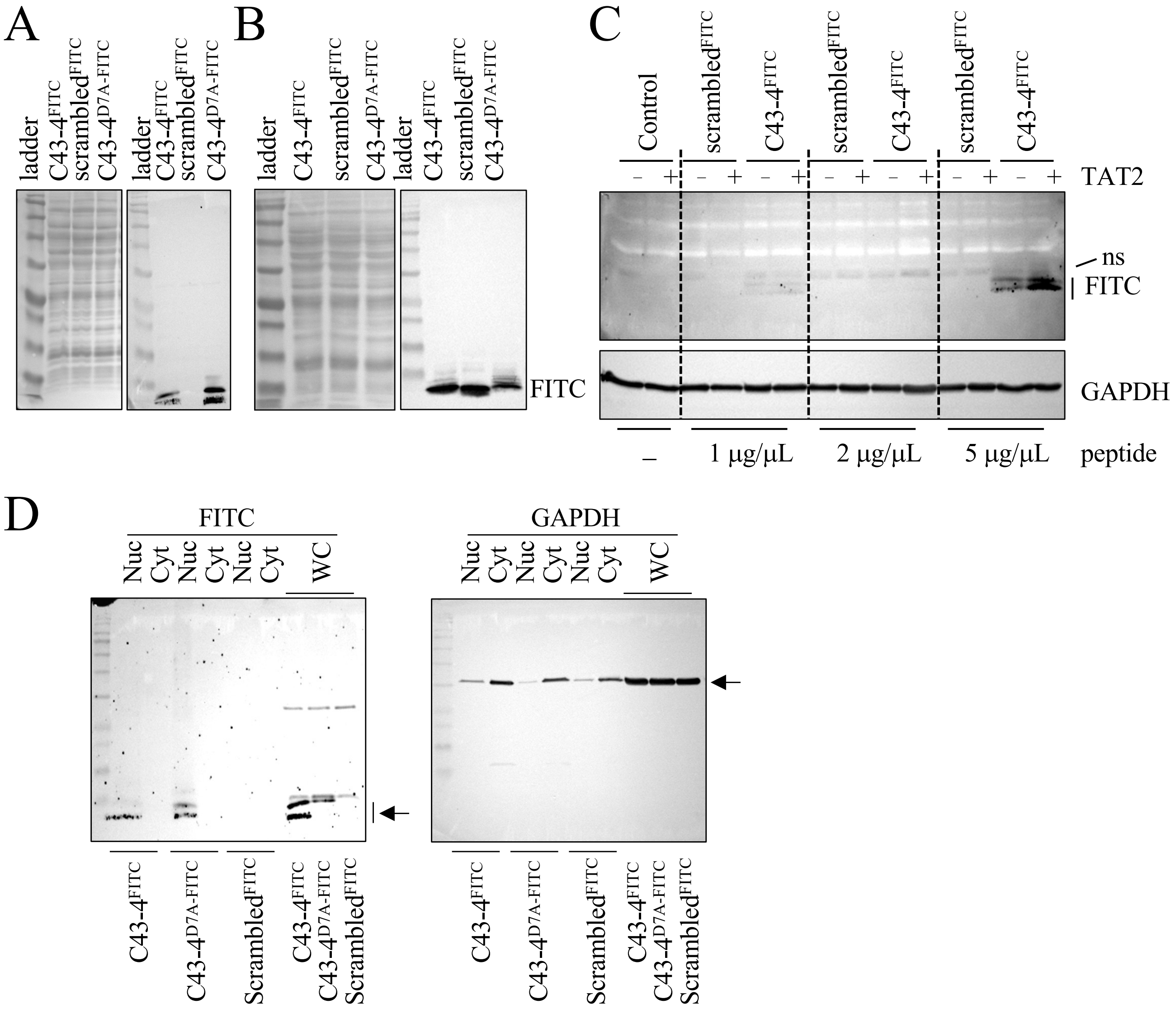

### Supplemental Figure 13

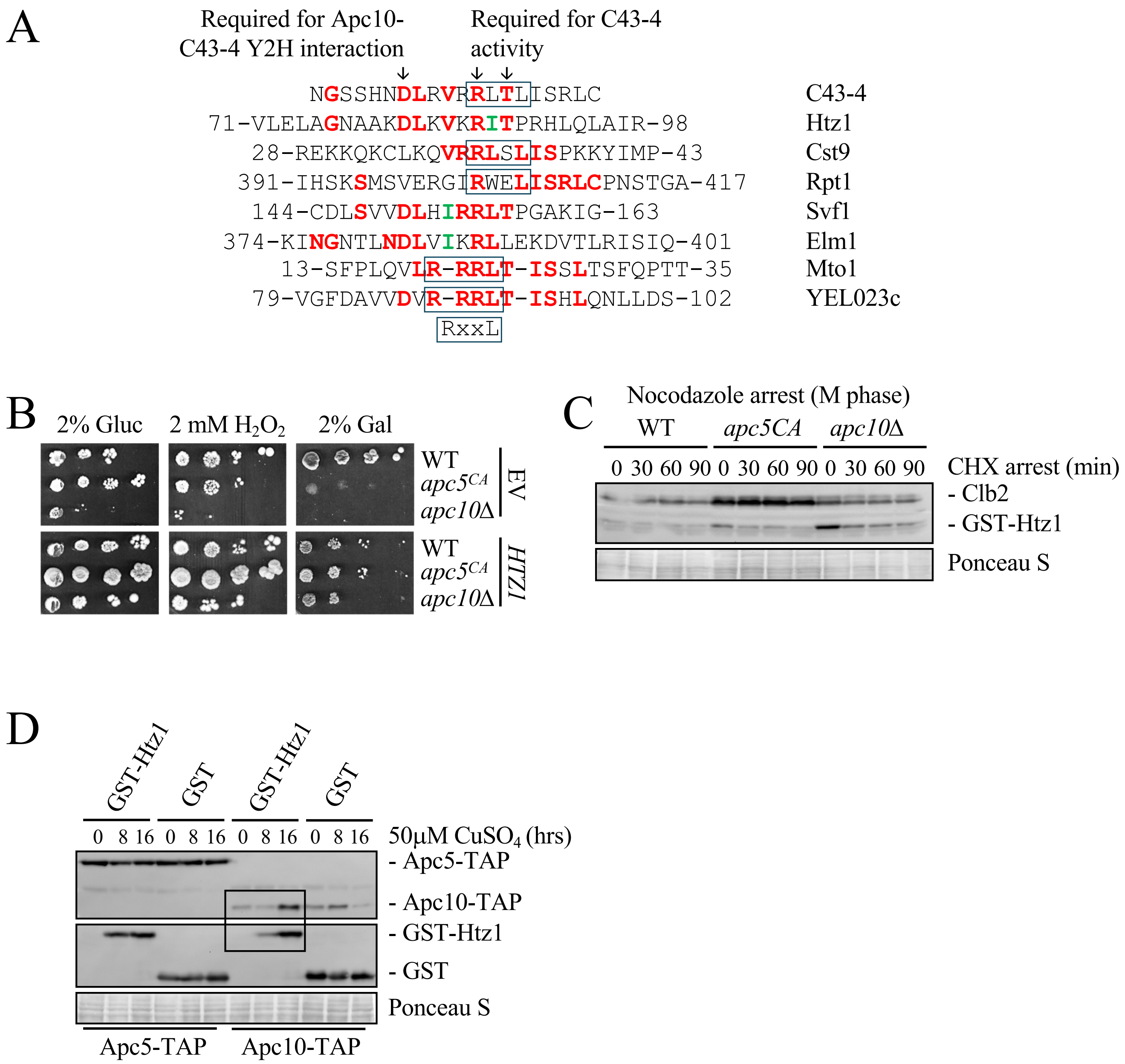

**Supplemental Table 1.** Data for **Figure 2C** using OASIS2. The log rank stats test was used to determine P-values.

| 2% Glucose | Mean Survival (generations) | 95% confidence interval | # subjects scored |
|---|---|---|---|
| WT + 0 uM M2I-1 | 21.43 ± 1.61 | 18.27 ~ 24.60 | 30 |
| WT + 0.1 uM M2I-1 | 23.69 ± 1.55 | 20.65 ~ 26.73 | 32 |
| WT + 1 uM M2I-1 | 27 ± 1.47 | 24.12 ~ 29.88 | 32 |
| WT + 10 uM M2I-1 | 29.41 ± 1.25 | 26.95 ~ 31.87 | 32 |

| Condition | $X^2$ | p-value |
|---|---|---|
| WT + 0 mM M2I vs WT + 0.1 mM M2I | 1.04 | 0.3087 |
| WT + 0 mM M2I vs WT + 1 mM M2I | 4.22 | 0.0399 |
| WT + 0 mM M2I vs WT + 10 mM M2I | 9.67 | 0.0019 |

**Supplemental Table 2.** Data for **Figure 2D** using the log rank test as part of the OASIS2 stats package.

| 2% Glucose | Mean Survival (generations) | 95% confidence interval | # subjects scored |
|---|---|---|---|
| *apc5* + 0 uM M2I-1 | 17.81 ± 0.73 | 16.37 ~ 19.25 | 32 |
| *apc5* + 0.1 uM M2I-1 | 20.47 ± 0.89 | 18.72 ~ 22.22 | 32 |
| *apc5* + 1 uM M2I-1 | 19.69 ± 0.87 | 17.98 ~ 21.40 | 32 |
| *apc5* + 10 uM M2I-1 | 21.44 ± 1.31 | 18.87 ~ 24.00 | 32 |

| Condition | $X^2$ | p-value |
|---|---|---|
| *apc5* + 0 µM M2I vs WT + 0 µM M2I | 8.69 | 0.0032 |
| *apc5* + 0 µM M2I vs *apc5* + 0.1 µM M2I | 6.81 | 0.009 |
| *apc5* + 0 µM M2I vs *apc5* + 1 µM M2I | 3.83 | 0.0502 |
| *apc5* + 0 µM M2I vs *apc5* + 10 µM M2I | 11.1 | 0.0009 |

**Supplemental Table 3.** Data for **Figure 5A** using a log rank test as part of the OASIS2 stats package.

| Trp- 2% Glucose | Mean Survival (generations) | 95% confidence interval | # subjects scored |
|---|---|---|---|
| WT + EV | 10.14 ± 0.68 | 8.79 ~ 11.48 | 51 |
| WT + C3-1B | 10.42 ± 0.67 | 9.11 ~ 11.73 | 57 |
| WT + C2-4B | 15.47 ± 0.72 | 14.07 ~ 16.88 | 57 |
| WT + C3-1 | 9.36 ± 0.49 | 8.39 ~ 10.33 | 50 |
| WT + C4-3 | 10.56 ± 0.8 | 8.98 ~ 12.13 | 54 |
| WT + C9-5 | 9.47 ± 0.39 | 8.70 ~ 12.13 | 53 |
| WT + C13-3 | 15.56 ± 0.95 | 13.70 ~ 17.42 | 41 |
| WT+ C43-3 | 10.91 ± 0.78 | 9.38 ~ 12.45 | 46 |

| Condition | $X^2$ | p-value |
|---|---|---|
| WT + EV vs WT + C3.1B | 0.21 | 0.647 |
| WT + EV vs WT + C2-4B | 19.62 | 0.0000094 |
| WT + EV vs WT + C3-1 | 1.32 | 0.2511 |
| WT + EV vs WT + C4-3 | 0.23 | 0.6279 |
| WT + EV vs WT + C9-6 | 1.32 | 0.2499 |
| WT + EV vs WT + C13-3 | 17.74 | 0.000025 |
| WT + EV vs WT + C43-4 | 0.29 | 0.578 |

**Supplemental Table 4.** Data for **Figure 5B** using a log rank test as part of the OASIS2 stats package.

| 2% Glucose | Mean Survival (generations) | 95% confidence interval | # subjects scored |
|---|---|---|---|
| WT + EV | 20.87 ± 1.45 | 18.03 ~ 23.72 | 31 |
| WT + C2.4B | 25.66 ± 1.56 | 22.61 ~ 28.71 | 32 |
| WT + C13.3 | 19.63 ± 1.53 | 16.63 ~ 22.62 | 32 |
| WT + C43.4 | 21.9 ± 1.61 | 18.76 ~ 25.05 | 31 |

| Condition | $X^2$ | p-value |
|---|---|---|
| WT + EV vs WT + C2-4B | 3.29 | 0.0696 |
| WT + EV vs WT + C13-3 | 0.14 | 0.713 |
| WT + EV vs WT + C43-4 | 0.28 | 0.5975 |

**Supplemental Table 5.** Data for **Figure 5C** using a log rank test as part of the OASIS2 stats package.

| | Mean Survival (generations) | 95% confidence interval | # subjects scored |
|---|---|---|---|
| WT + 2% Glu | 22 ± 1.74 | 18.58 ~ 25.42 | 32 |
| WT + 0.5% Gal/1.5% Suc | 24.47 ± 1.88 | 20.79 ~ 28.15 | 32 |
| WT + C43.4 2% Glu | 23.63 ± 1.45 | 20.77 ~ 26.48 | 32 |
| WT + C43.4 0.5%Gal/1.5% Suc | 29.66 ± 1.97 | 25.79 ~ 33.53 | 32 |

| Condition | $X^2$ | p-value |
|---|---|---|
| WT 2% Glu vs WT + 0.5% Gal | 0.96 | 0.3261 |
| WT 2% Glu vs C43.4 + 2% Glu | 0.11 | 0.7446 |
| WT 2% Glu vs C43.4 + 0.5% Gal | 7.66 | 0.0057 |

**Supplemental Table 6.** Mean and max lifespan, and 95% confidence intervals for CLS curves shown in **Figures 6A-E**.

| Strain | Mean Survival (Days) | 95% Confidence Interval | Max survival (Days) |
|---|---|---|---|
| **EV 2% Suc** | 5.065 ± 0.3029 | 4.906 to 5.225 | 9 |
| **EV 1% Gal** | 5.177 ± 0.1609 | 5.095 to 5.259 | 9 |
| **C2-4B 2% Suc** | 5.189 ± 0.3483 | 5.005 to 5.375 | 9 |
| **C2-4B 1% Gal** | 5.160 ± 0.2871 | 5.015 to 5.307 | 9 |
| **C9-5 2% Suc** | 5.169 ± 0.2938 | 5.014 to 5.324 | 9 |
| **C9-5 1% Gal** | 5.060 ± 0.2778 | 4.922 to 5.199 | 9 |
| **C13-3 2% Suc** | 5.216 ± 0.2339 | 5.092 to 5.341 | 9 |
| **C13-3 1% Gal** | 5.102 ± 0.3385 | 4.935 to 5.270 | 9 |
| **C43-4 2% Suc** | 5.243 ± 0.2187 | 5.128 to 5.359 | 9 |
| **C43-4 1% Gal** | 6.549 ± 0.3936 | 6.134 to 6.954 | 13 |
| **EV 2% Suc (Ura+)** | 6.318 ± 0.4558 | 5.871 to 6.765 | 13 |
| **EV 1% Gal (Ura+)** | 6.365 ± 0.5360 | 5.847 to 6.882 | 13 |
| **C43-4 2% Suc (Ura+)** | 6.533 ± 0.4193 | 6.132 to 6.936 | 13 |
| **C43-4 1% Gal (Ura+)** | 8.736 ± 0.3029 | 8.035 to 9.423 | 19 |

**Supplemental Table 7**. Statistical analysis performed using OASIS for lifespan of N2 wild type worms in the presence or absence of C43-4. Linked to **Figure 11C**.

| | Mean survival (Days) | 95% confidence interval | # of subjects scored | # of censors | P - value vs corresponding N2 |
|---|---|---|---|---|---|
| **Lifespan (Trial 1)** | | | | | $1.2 \times 10^{-6}$ |
| **N2** | 19.95 ± 0.48 | 19.00 ~ 20.90 | 106 | 3 | |
| **C43.4** | 23.45 ± 0.45 | 22.56 ~ 24.33 | 148 | 17 | |
| **Lifespan (Trial 2)** | | | | | 0.0002 |
| **N2** | 19.07 ± 0.44 | 18.20 ~ 19.94 | 120 | 18 | |
| **C43.4** | 23.22 ± 0.45 | 22.33 ~ 24.11 | 98 | 26 | |
| **Lifespan (Trial 3)** | | | | | 0.0002 |
| **N2** | 20.61 ± 0.53 | 19.58 ~ 21.65 | 98 | 13 | |
| **C43.4** | 23.96 ± 0.47 | 23.04 ~ 24.89 | 124 | 32 | |

**Supplemental Table 8**. **(A)** Statistical analysis performed using OASIS showing mean lifespans of N2 wild type and *daf-16* worms in the presence or absence of C43-4. **(B)** P - values comparing lifespan curves for the different combinations used. Linked to **Figure 11D**.

# A

| | Mean survival (Days) | 95% confidence interval | # of subjects scored | # of censors |
|---|---|---|---|---|
| **Lifespan (Trial 1)** | | | | |
| **N2** | 17.52 ± 0.26 | 17.01 ~ 18.03 | 186 | 2 |
| **C43-4** | 19.37 ± 0.35 | 18.69 ~ 20.06 | 159 | 7 |
| ***daf-16*(mu86)** | 14.05 ± 0.24 | 13.59 ~ 14.51 | 129 | 5 |
| **C43.4; *daf-16*(mu86)** | 15.73 ± 0.28 | 15.18 ~ 16.27 | 146 | 5 |
| **Lifespan (Trial 2)** | | | | |
| **N2** | 17.76 ± 0.3 | 17.17 ~ 18.36 | 176 | 1 |
| **C43-4** | 19.48 ± 0.31 | 18.86 ~ 20.09 | 164 | 7 |
| ***daf-16*(mu86)** | 14.65 ± 0.25 | 14.15 ~ 15.14 | 155 | 5 |
| **C43.4; *daf-16*(mu86)** | 16.35 ± 0.27 | 15.82 ~ 16.89 | 153 | 7 |
| **Lifespan (Trial 3)** | | | | |
| **N2** | 19.35 ± 0.38 | 18.60 ~ 20.10 | 133 | 2 |
| **C43-4** | 21.49 ± 0.41 | 20.69 ~ 22.29 | 132 | 8 |
| ***daf-16*(mu86)** | 15.26 ± 0.23 | 14.81 ~ 15.71 | 148 | 4 |
| **C43.4; *daf-16*(mu86)** | 17.5 ± 0.26 | 16.99~ 18.02 | 159 | 4 |

B

| Trial 1 condition | $X^2$ | P – value |
|---|---|---|
| N2 v.s. C43-4 | 23.54 | 0.0000012 |
| N2 v.s. *daf-16*(mu86) | 83.08 | 0 |
| N2 v.s. C43-4;*daf-16*(mu86) | 19 | 0.000013 |
| C43-4 v.s. *daf-16*(mu86) | 130.76 | 0 |
| C43-4 v.s. C43-4;*daf-16*(mu86) | 65.74 | 0 |
| *daf-16*(mu86) v.s. C43-4;*daf-16*(mu86) | 24.05 | 9.40E-07 |
| **Trial 2 condition** | $X^2$ | **P – value** |
| N2 v.s. C43-4 | 13.98 | 0.0002 |
| N2 v.s. *daf-16*(mu86) | 59.39 | 0 |
| N2 v.s. C43-4;*daf-16*(mu86) | 13.02 | 0.0003 |
| C43-4 v.s. *daf-16*(mu86) | 117.1 | 0 |
| C43-4 v.s. C43-4;*daf-16*(mu86) | 52.78 | 0 |
| *daf-16*(mu86) v.s. C43-4;*daf-16*(mu86) | 20.86 | 0.0000049 |
| **Trial 3 condition** | $X^2$ | **P – value** |
| N2 v.s. C43-4 | 13.6 | 0.0002 |
| N2 v.s. *daf-16*(mu86) | 80.58 | 0 |
| N2 v.s. C43-4;*daf-16*(mu86) | 20.27 | 0.0000067 |
| C43-4 v.s. *daf-16*(mu86) | 134.31 | 0 |
| C43-4 v.s. C43-4;*daf-16*(mu86) | 63.77 | 0 |
| *daf-16*(mu86) v.s. C43-4;*daf-16*(mu86) | 41.06 | 0 |

**Supplemental Table 9**. **(A)** Statistical analysis performed using OASIS showing mean lifespans of N2 wild type and *aat-2* worms in the presence or absence of C43-4. **(B)** P - values comparing lifespan curves for the different combinations used. Linked to **Figure 11E**.

# A

| | Mean survival (Days) | 95% confidence interval | # of subjects scored | # of censors |
|---|---|---|---|---|
| **Lifespan (Trial 1)** | | | | |
| **N2** | 13.92 ± 0.16 | 13.60 ~ 14.23 | 191 | 4 |
| **C43.4** | 14.92 ± 0.19 | 14.54 ~ 15.30 | 197 | 3 |
| **aak-2(ok524)** | 12.43 ± 0.17 | 12.10 ~ 12.76 | 164 | 7 |
| **C43.4; aak-2(ok524)** | 13.61 ± 0.17 | 13.28 ~ 13.94 | 160 | 3 |
| **Lifespan (Trial 2)** | | | | |
| **N2** | 16.3 ± 0.3 | 15.72 ~ 16.88 | 162 | 3 |
| **C43.4** | 17.95 ± 0.31 | 17.35 ~ 18.55 | 166 | 2 |
| **aak-2(ok524)** | 14.96 ± 0.19 | 14.58 ~ 15.34 | 175 | 8 |
| **C43.4; aak-2(ok524)** | 15.23 ± 0.23 | 14.78 ~ 15.67 | 146 | 21 |
| **Lifespan (Trial 3)** | | | | |
| **N2** | 17.76 ± 0.3 | 17.17 ~ 18.36 | 176 | 1 |
| **C43.4** | 19.62 ± 0.32 | 18.98 ~ 20.25 | 164 | 7 |
| **aak-2(ok524)** | 15.6 ± 0.21 | 15.18 ~ 16.01 | 189 | 16 |
| **C43.4; aak-2(ok524)** | 15.55 ± 0.28 | 14.99 ~ 16.10 | 140 | 10 |

B

| Trial 1 condition | $X^2$ | P – value |
|---|---|---|
| N2 v.s. C43-4 | 18.93 | 0.000014 |
| N2 v.s. *aak-2*(ok524) | 34.05 | 5.30E-09 |
| N2 v.s. C43-4;*aak-2*(ok524) | 0.98 | 0.3218 |
| C43-4 v.s. *aak-2*(ok524) | 82.52 | 0 |
| C43-4 v.s. C43-4;*aak-2*(ok524) | 28.74 | 8.30E-08 |
| *aak-2*(ok524) v.s. C43-4;*aak-2*(ok524) | 22.61 | 0.000002 |
| **Trial 2 condition** | $X^2$ | **P – value** |
| N2 v.s. C43-4 | 13.66 | 0.0002 |
| N2 v.s. *aak-2*(ok524) | 16.9 | 0.000039 |
| N2 v.s. C43-4;*aak-2*(ok524) | 8.63 | 0.0033 |
| C43-4 v.s. *aak-2*(ok524) | 75.5 | 0 |
| C43-4 v.s. C43-4;*aak-2*(ok524) | 52.38 | 0 |
| *aak-2*(ok524) v.s. C43-4;*aak-2*(ok524) | 1.32 | 0.2511 |
| **Trial 3 condition** | $X^2$ | **P – value** |
| N2 v.s. C43-4 | 13.98 | 0.0002 |
| N2 v.s. *aak-2*(ok524) | 40.44 | 0 |
| N2 v.s. C43-4;*aak-2*(ok524) | 28.12 | 1.10E-07 |
| C43-4 v.s. *aak-2*(ok524) | 100.01 | 0 |
| C43-4 v.s. C43-4;*aak-2*(ok524) | 74.26 | 0 |
| *aak-2*(ok524) v.s. C43-4;*aak-2*(ok524) | 0.02 | 0.8792 |

**Supplemental Table 10. (A)** Statistical analysis performed using OASIS showing mean lifespans of N2 wild type and *akt-1* worms in the presence or absence of C43-4. **(B)** P - values comparing lifespan curves for the different combinations used. Linked with **Figure 11F**.

A

| | Mean survival (Days) | 95% confidence interval | # of subjects scored | # of censors |
|---|---|---|---|---|
| **Lifespan (Trial 1)** | | | | |
| **N2** | 19.12 ± 0.36 | 18.41 ~ 19.83 | 133 | 2 |
| **C43.4** | 21.55 ± 0.4 | 20.76 ~ 22.34 | 135 | 8 |
| ***akt-1*(ok525)** | 23.02 ± 0.55 | 21.94 ~ 24.10 | 184 | 4 |
| **C43.4; *akt-1*(ok525)** | 23.79 ± 0.71 | 22.40 ~ 25.18 | 146 | 7 |
| **Lifespan (Trial 2)** | | | | |
| **N2** | 19.83 ± 0.32 | 19.20 ~ 20.46 | 162 | 7 |
| **C43.4** | 20.51 ± 0.4 | 19.73 ~ 21.28 | 134 | 7 |
| ***akt-1*(ok525)** | 22.65 ± 0.56 | 21.54 ~ 23.75 | 131 | 1 |
| **C43.4; *akt-1*(ok525)** | 22.97 ± 0.58 | 21.82 ~ 24.11 | 126 | 15 |
| **Lifespan (Trial 3)** | | | | |
| **N2** | 17.86 ± 0.29 | 17.28 ~ 18.43 | 156 | 4 |
| **C43.4** | 20.28 ± 0.39 | 19.51 ~ 21.05 | 124 | 11 |
| ***akt-1*(ok525)** | 22.75 ± 0.44 | 21.90 ~ 23.60 | 159 | 9 |
| **C43.4; *akt-1*(ok525)** | 23.74 ± 0.6 | 22.56 ~ 24.91 | 119 | 6 |

B

| Trial 1 condition | $X^2$ | P – value |
|---|---|---|
| N2 v.s. C43-4 | 21.03 | 0.0000045 |
| N2 v.s. *akt-1*(ok525) | 41.21 | 0 |
| N2 v.s. C43-4;*akt-1*(ok525) | 42.45 | 0 |
| C43-4 v.s. *akt-1*(ok525) | 13.89 | 0.0002 |
| C43-4 v.s. C43-4;*akt-1*(ok525) | 19.52 | 0.00001 |
| *akt-1*(ok525) v.s. C43-4;*akt-1*(ok525) | 3.27 | 0.0708 |
| **Trial 2 condition** | $X^2$ | **P – value** |
| N2 v.s. C43-4 | 3.59 | 0.0581 |
| N2 v.s. *akt-1*(ok525) | 34.23 | 4.90E-09 |
| N2 v.s. C43-4;*akt-1*(ok525) | 38.03 | 0 |
| C43-4 v.s. *akt-1*(ok525) | 17.2 | 0.000034 |
| C43-4 v.s. C43-4;*akt-1*(ok525) | 20.77 | 0.0000052 |
| *akt-1*(ok525) v.s. C43-4;*akt-1*(ok525) | 0.21 | 0.6454 |
| **Trial 3 condition** | $X^2$ | **P – value** |
| N2 v.s. C43-4 | 26.94 | 2.10E-07 |
| N2 v.s. *akt-1*(ok525) | 84.84 | 0 |
| N2 v.s. C43-4;*akt-1*(ok525) | 88.73 | 0 |
| C43-4 v.s. *akt-1*(ok525) | 22.2 | 0.0000025 |
| C43-4 v.s. C43-4;*akt-1*(ok525) | 33.04 | 9.00E-09 |
| *akt-1*(ok525) v.s. C43-4;*akt-1*(ok525) | 3.63 | 0.0567 |
